# The evolutionary history of Neandertal and Denisovan Y chromosomes

**DOI:** 10.1101/2020.03.09.983445

**Authors:** Martin Petr, Mateja Hajdinjak, Qiaomei Fu, Elena Essel, Hélène Rougier, Isabelle Crevecoeur, Patrick Semal, Liubov V. Golovanova, Vladimir B. Doronichev, Carles Lalueza-Fox, Marco de la Rasilla, Antonio Rosas, Michael V. Shunkov, Maxim B. Kozlikin, Anatoli P. Derevianko, Benjamin Vernot, Matthias Meyer, Janet Kelso

## Abstract

Ancient DNA has allowed the study of various aspects of human history in unprecedented detail. However, because the majority of archaic human specimens preserved well enough for genome sequencing have been female, comprehensive studies of Y chromosomes of Denisovans and Neandertals have not yet been possible. Here we present sequences of the first Denisovan Y chromosomes (*Denisova 4* and *Denisova 8*), as well as the Y chromosomes of three late Neandertals (*Spy 94a*, *Mezmaiskaya 2* and *El Sidrón 1253*). We find that the Denisovan Y chromosomes split around 700 thousand years ago (kya) from a lineage shared by Neandertal and modern human Y chromosomes, which diverged from each other around 370 kya. The phylogenetic relationships of archaic and modern human Y chromosomes therefore differ from population relationships inferred from their autosomal genomes, and mirror the relationships observed on the level of mitochondrial DNA. This provides strong evidence that gene flow from an early lineage related to modern humans resulted in the replacement of both the mitochondrial and Y chromosomal gene pools in late Neandertals. Although unlikely under neutrality, we show that this replacement is plausible if the low effective population size of Neandertals resulted in an increased genetic load in their Y chromosomes and mitochondrial DNA relative to modern humans.

## Introduction

Ancient DNA (aDNA) has transformed our understanding of human evolutionary history, revealing complex patterns of population migration, turnover and gene flow, including admixture from archaic humans into modern humans around 55 thousand years ago (kya) (*1–4*). The majority of insights into the relationships between archaic and modern humans have been based on autosomal sequences, which represent a composite of genealogies of any individual’s ancestors. Although mitochondrial DNA (mtDNA) and Y chromosomes only provide information about single maternal and paternal lineages, they offer a unique perspective on various aspects of population history such as sex-specific migration and other cultural phenomena (*5–7*). Furthermore, because of their lower effective population size (*N_e_*) compared to autosomal loci, coalescent times of mtDNA and Y chromosomes sampled from two populations provide an upper bound for the last time they experienced gene flow. In this respect, the mtDNA and autosomal sequences of Neandertals, Denisovans and modern humans have revealed puzzling phylogenetic discrepancies. Their autosomal genomes show that Neandertals and Denisovans are sister groups that split from a common ancestor with modern humans between 550–765 kya (*8*). In contrast, with the TMRCA with modern humans of 360-468 kya, mtDNAs of Neandertals are more similar to the mtDNAs of modern humans than to those of Denisovans (*9*). Intriguingly, ∼400 ky old early Neandertals from Sima de los Huesos were shown to carry mitochondrial genomes related to Denisovan mtDNAs, which is concordant with the autosomal relationships between these groups of archaic humans (*10, 11*). Based on these results it has been suggested that Neandertals originally carried a Denisovan-like mtDNA which was later completely replaced via gene flow from an early lineage related to modern humans (*9, 11*).

Y chromosomes of Neandertals and Denisovans could provide an important additional source of information about population splits and gene flow events between archaic and modern humans or populations related to them. However, with the exception of a small amount of Neandertal Y chromosome coding sequence (118 kb, (*12*)), none of the male Neandertals or Denisovans studied to date have yielded sufficient amounts of endogenous DNA to allow comprehensive studies of entire Y chromosomes.

## Specimens, DNA capture and genotyping

Previous genetic studies identified two male Denisovans, *Denisova 4 (*55–84 ky old) and *Denisova 8* (106–136 ky old) (*13, 14*), and two male late Neandertals, *Spy 94a* (38-39 ky old) and *Mezmaiskaya 2* (43-45 ky old) (*15*) (Figure 1A). To enrich for hominin Y chromosome DNA of these individuals, we designed DNA capture probes targeting ∼6.9 Mb of the non-recombining portion of the human Y chromosome sequence (Figure 1C, Supplementary Information). Using these probes, we performed hybridization capture on selected single-stranded DNA libraries from *Denisova 4*, *Denisova 8*, *Spy 94a,* and *Mezmaiskaya 2* (Supplementary Information). The captured DNA molecules were sequenced from both ends, overlapping reads were merged and aligned to the human reference genome (hg19/GRCh37) (Supplementary Information). Reads of at least 35 base-pair (bp) in length that aligned uniquely to the capture target regions were retained for further analysis.

**Figure 1.**
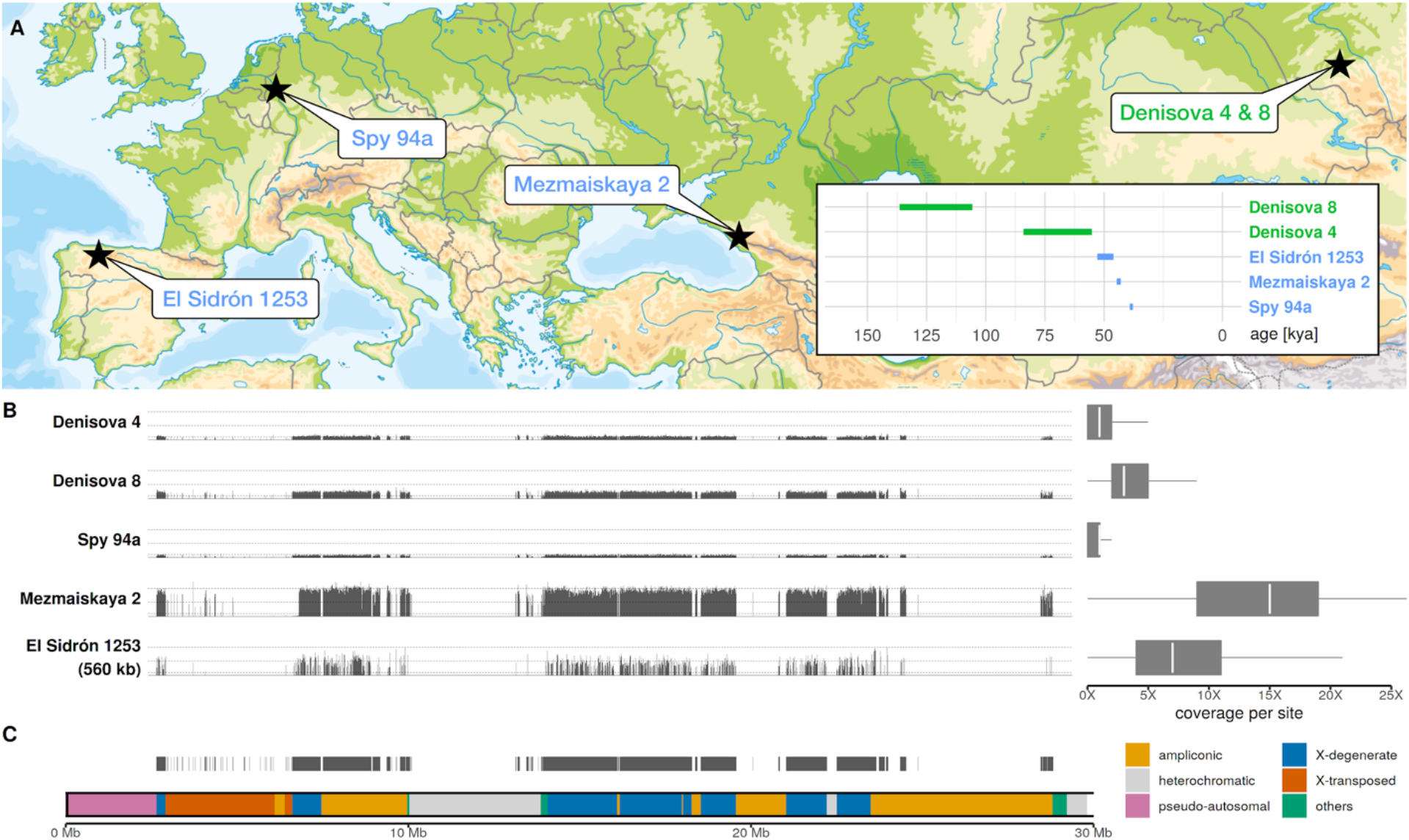
Geographical locations, ages and sequencing coverage of the male archaic humans in our study. **(A)** Locations of archaeological sites where the five archaic human specimens have been found. Estimates of the ages shown as an inset (*14–16*). (**B**) **Left** - Spatial distribution of sequencing coverage for each archaic human Y chromosome along the ∼6.9 Mb of capture target regions. The heights of the thin vertical bars represent average coverage in each target region. The chromosomal coordinates are aligned to match the Y chromosome structure depicted in panel C. **Right -** Distribution of coverage across all target sites for each archaic Y chromosome on the left. **(C)** Genomic structure of the portion of the human Y chromosome targeted for capture. Thin black vertical lines show the position of individual target capture regions. The coordinates of Y chromosome regions were taken from (*32*).

Of the total ∼6.9 Mb of Y chromosome capture target regions, we generated 1.4X for *Denisova 4*, 3.5X for *Denisova 8*, 0.8X for *Spy 94a* and 14.3X for *Mezmaiskaya 2* (Figure 1B, Table S4.1). In addition, we sequenced 7.9X coverage of a smaller subset of the Y chromosome of the ∼46-53-ky-old *El Sidrón 1253* Neandertal from Spain (Figure 1B, Table S4.1) (*16*) by capturing a set of previously generated double-stranded, UDG-treated libraries (*17*) using a ∼560 kb capture array designed to study modern human Y chromosome variation (*5*). This data provides an opportunity for validating our results with different sample preparation and capture strategies.

To call genotypes for the captured archaic human Y chromosomes, we leveraged the haploid nature of the human Y chromosome and implemented a consensus approach that requires 90% of the reads observed at each site to agree on a single allele, restricting to sites covered by at least three reads (Supplementary information; Table S5.1). This approach minimizes the impact of aDNA damage on genotyping accuracy (Figure S5.1) while allowing for a small proportion of sequencing errors, contamination or misalignment (Supplementary Information). To obtain a reference panel of modern human Y chromosomes for the analyses below, we applied the same genotype calling procedure to a set of previously published modern human Y chromosomes (*6, 18, 19*) (Supplementary Information).

## Archaic Y chromosome phylogeny

To determine the relationships between Denisovan, Neandertal and modern human Y chromosomes we constructed a neighbor-joining tree from the alignment of Y chromosome genotype calls (Supplementary Information). Unlike the rest of the nuclear genome, which puts Denisovans and Neandertals as sister groups to modern humans (*2*), we found that the Denisovan Y chromosomes form a separate lineage that split before Neandertal and modern human Y chromosomes diverged from each other (Figure 2A, 100% bootstrap support for both ancestral nodes). Notably, all three late Neandertal Y chromosomes cluster together and fall outside of the variation of present-day human Y chromosomes (Figure 2A, 100% bootstrap support), which includes the African Y chromosome lineage A00 known to have diverged from all other present-day human Y chromosomes around 250 kya (*6*).

**Figure 2.**
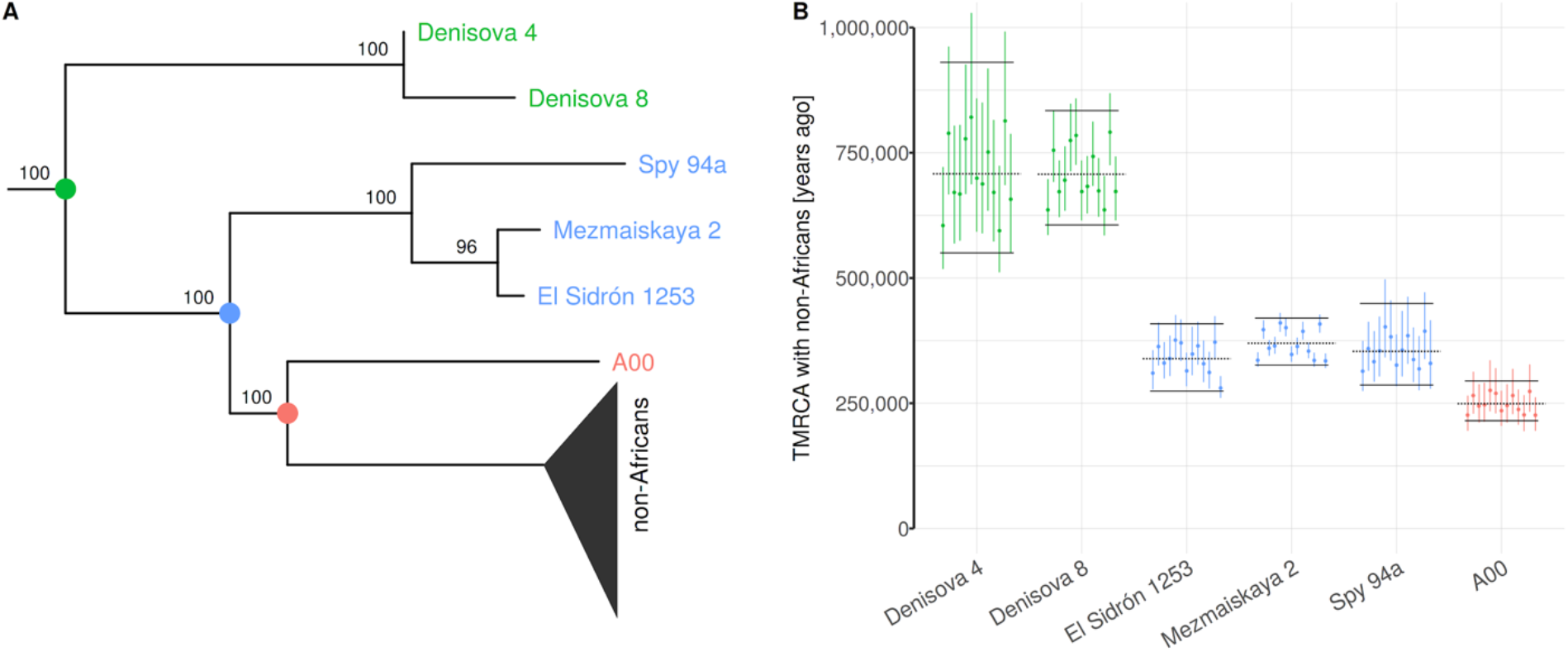
Phylogenetic relationships between archaic and modern human Y chromosomes. **(A)** Neighbor-joining tree based on the alignment of Y chromosome genotype calls, excluding C-to-T and G-to-A polymorphisms to mitigate the effects of aDNA damage. Numbers next to the internal nodes show bootstrap support for the three major clades (green - Denisovans, blue - Neandertals, red – deeply divergent African lineage A00) based on 100 bootstrap replicates. The tree was rooted using a chimpanzee Y chromosome as the outgroup. We note that the terminal branch lengths are not informative about the ages of specimens due to differences in sequence quality (Figure 1A). **(B)** Distributions of the times to the most recent common ancestor (TMRCA) between Y chromosomes listed along the x-axis and a panel of 13 non-African Y chromosomes. Each dot represents a TMRCA estimate based on a single non-African Y chromosome, with error bars showing approximate 95% C.I. based on resampling of branch counts (Supplementary Information). Black horizontal lines show the mean TMRCA calculated across the full non-African panel (dotted lines) with bootstrap-based 95% C.I. (solid lines).

## Ages of Y chromosomal ancestors

To estimate the time to the most recent common ancestor (TMRCA) of archaic and modern human Y chromosomes we followed an approach similar to that taken by Mendez *et al.*, expressing this TMRCA relative to the deepest known split within present-human Y chromosomes (African Y chromosome lineage A00, (*6, 12*), Supplementary Information). This has the advantage of not relying on private mutations on the archaic human branch which makes it robust to low coverage and aDNA damage which vary significantly between samples (Figure 2A, Table S4.1, Figure S5.1).

We first calculated the mutation rate in the total 6.9 Mb target regions to be 7.34×10^-10^ per bp per year (bootstrap CI: 6.27-8.46×10^-10^; Figure S7.1, Table S7.2, Supplementary Information). Using this mutation rate, we estimated the TMRCA of the African A00 lineage and a set of non-African Y chromosomes from the SGDP panel (*6, 19*) at ∼249 kya (bootstrap CI: 213-293 kya; Figure S7.1, Table S7.2, Supplementary information). These estimates are consistent with values inferred from larger-scale studies of present-day human Y chromosomes (*6, 18*), suggesting that the Y chromosomal regions we defined for capture are not unusual in terms of their mutation rate.

Second, assuming the A00 divergence time of 249 kya, we inferred TMRCAs between archaic Y chromosomes and present-day non-African Y chromosomes for each archaic individual at a time (Figure S7.4, Table S7.3, Supplementary information). We found that the two Denisovan Y chromosomes (*Denisova 4* and *Denisova 8*) split from the modern human lineage around 700 kya (CI: 607-833 kya for the higher coverage *Denisova 8*, Figure 2B, Table S7.3). In contrast, the three Neandertal Y chromosomes split from the modern human lineage about 350 kya: 353 kya for *Spy 94a* (286-449 kya), 369 kya for *Mezmaiskaya 2* (326-419 kya) and 339 kya for *El Sidrón 1253* (275-408 kya) (Figure 2B, Table S7.3). Additionally, we used the proportions of sharing of derived alleles with the high-coverage Mezmaiskaya 2 Y chromosome to estimate the TMRCA of the three Neanderthal Y chromosomes at around 100 kya (Figure S7.14 and S7.15). We validated the robustness of all TMRCA estimates by repeating the analyses using filters of varying levels of stringency and different genotype calling methods (Figures S7.9, S7.11, S7.13). Similarly, although we detected some evidence of capture bias in the data (Figure S4.5), we observed no significant differences between capture data and shotgun sequences or between individuals showing different read length distributions, indicating that the effect of technical biases on our inferences is negligible (Figure S7.11).

The Denisovan-modern human Y chromosome TMRCA estimates are in good agreement with population split times inferred from autosomal sequences, suggesting that the differentiation of Denisovan Y chromosomes from modern humans occurred through a simple population split (*20*). In contrast, the Neandertal-modern human Y chromosome TMRCAs are significantly younger than the inferred population split time (Figure 3A) and consistent with a time window for gene flow from a lineage related modern humans into Neandertals inferred from mtDNA (*9*) and autosomal sequences (*21, 22*).

**Figure 3.**
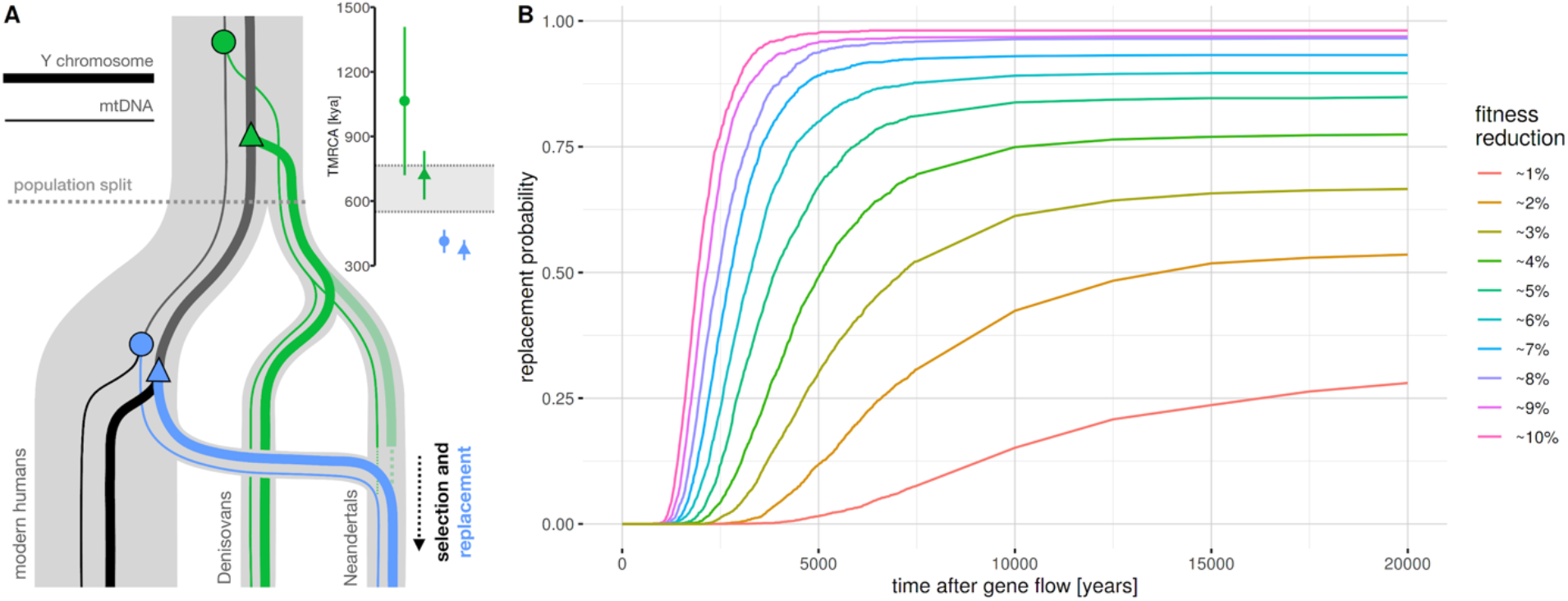
Proposed model for the replacement of Neandertal Y chromosomes and mtDNA. **(A)** Schematic representation of the relationships between archaic and modern human mtDNA (thin lines) and Y chromosomes (thick lines) based on current phylogenetic inferences, with the hypothesized time-window for selection. The semi-transparent Neandertal lineage indicates an as yet unsampled, hypothetical Y chromosome which was replaced by an early lineage related to modern human Y chromosomes. Positions of relevant most recent common ancestors with modern human lineages are shown for mtDNA (circle nodes) and Y chromosomes (triangle nodes). Inset shows TMRCA estimates for the four nodes in the diagram: Y chromosome TMRCAs as estimated by our study, mtDNA TMRCAs estimates from the literature (*9, 10*). The grey horizontal bar highlights the 95% C.I. for the population split time between archaic and modern humans (*8*). **(B)** Probability of replacement of a non-recombining, uniparental Neandertal locus as a function of time after gene flow, assuming a given level of fitness burden relative to its modern human counterpart. Trajectories are based on forward simulations across a grid of parameters (Figure S8.1-S8.3, Supplementary Information). Modern human introgression was simulated in a single pulse at 5%.

## Disagreement with the previous Y chromosome TMRCA

Our estimates of the Neandertal-modern human TMRCA, including those obtained using the larger amount of data we generated from the same individual (TMRCA of ∼339 kya, Figure 2B), are substantially younger than the previous estimate of ∼588 kya from the *El Sidrón 1253* Neandertal (*12*). The previous estimate was based on ∼3X coverage of 118 kb of exome capture sequence and, due to the limited amount of data, used SNPs supported even by single reads (*12, 17*). Although the TMRCA inference procedure used by (*12*) does not rely directly on the counts of private mutations on the archaic lineage (Figure S7.4, Supplementary Information), it can still be affected by erroneous genotype calls, which can lead to shared derived variants being converted to the ancestral state, increasing the apparent TMRCA. Indeed, when we applied our stricter filtering criteria to the *El Sidrón 1253* data analyzed previously, we arrived at TMRCA estimates for *El Sidrón 1253* that are consistent with the other Neandertals in our study (Figure S7.12).

## The probability of replacement

The phylogenetic relationships of archaic and modern human Y chromosomes are similar to the observations made from mtDNA genomes (*9, 10*), suggesting that both mtDNA and Y chromosomes of early Neandertals have been replaced via gene flow from an early lineage related to modern humans, possibly as a result of the same population contact (Figure 3A, (*9*)). Although such contact has been proposed, previous work suggests gene flow from modern humans into Neandertals on the order of only a few percent (*21, 23*). Assuming neutrality, the fixation probability of a locus is equal to its initial frequency in a population (*24*). Therefore, the joint probability of both Neandertal mtDNA and Y chromosome replacements by their modern human counterparts under neutrality is even lower. However, several studies have suggested that due to their low *N_e_* and reduced efficacy of purifying selection, Neandertals may have accumulated an excess of deleterious variation compared to modern humans (*25, 26*) and sequencing of the exomes of three Neandertals directly demonstrated that they carried more deleterious alleles than present-day humans (*17*). To explore the dynamics of introgression into Neandertals, we simulated the frequency trajectories of a non-recombining, uniparental locus under a model of purifying selection (Supplementary Information, (*27*)). We simulated lower *N_e_* on the Neandertal lineage after its split from modern humans and accumulation of deleterious variants on both lineages across a grid of several relevant parameters (such as the time of the split between both populations or the amount of sequence under negative selection). For each combination of parameters, we then calculated the ratio of fitnesses of average Neandertal and average modern Y chromosomes produced by the simulation, and traced the trajectory of introgressed modern human Y chromosomes in Neandertals over 100 ky following 5% admixture (Figures S8.1 and S8.2, Supplementary information).

We found that even a small reduction in fitness of Neandertal Y chromosomes compared to modern human Y chromosomes has a strong effect on the probability of a complete replacement by introgressed modern human Y chromosomes (Figure 3B). Specifically, even a 1% reduction in Neandertal Y chromosome fitness increases the probability of replacement after 20ky to ∼25% and a 2% reduction in fitness increases this probability to ∼50%. This fitness reduction measure is an aggregate over all linked deleterious mutations on the Y chromosome and integrates a number of biological parameters, only a subset of which we consider here (Figure S8.1 and S8.3). Importantly, we note that although we simulated introgression of Y chromosomes, the abstract measure of fitness reduction of a non-recombining, uniparental locus can also be generalized to the case of mtDNA introgression (Figure 3B).

These results show that a model of higher genetic load in Neandertals is compatible with an increased probability of replacement of Neandertal mtDNA and Y chromosomes with their introgressed modern human counterparts. Furthermore, given the crucial role of the Y chromosome in reproduction and fertility, and its haploid nature, it is possible that deleterious mutations or structural variants on the Y chromosome have a dramatically larger impact on fitness than we considered in our simulations (*28*).

## Conclusions

Our results show that the Y chromosomes of late Neandertals represent an extinct lineage related to modern human Y chromosomes that introgressed into Neandertals some time between ∼370 kya and ∼100 kya. The presence of this Y chromosome lineage in all late Neandertals makes it unlikely that genetic changes that accumulated in Neandertal and modern human Y chromosomes prior to the introgression lead to incompatibilities between these groups of humans. We predict that the ∼400 ky old Sima de los Huesos individuals, who are early Neandertals but carry a Denisovan-like mtDNA (*10, 11*), should also carry a Y chromosome lineage more similar to Denisovans than to later Neandertals. Although complete replacement of mtDNA and Y chromosomes might seem surprising given that limited modern human gene flow has been detected in the genomes of late Neandertals (*15, 21–23*), mitochondrial-autosomal discrepancies are predicted by population genetic theory, and are relatively common during interspecific hybridization in the animal kingdom (*29–31*). Our simulations show that differences in genetic loads in uniparental loci between the two hybridizing populations is a plausible driver of this phenomenon.

## Acknowledgements

We would like to thank Svante Pääbo, Mark Stoneking, Benjamin Peter, Montgomery Slatkin, Laurits Skov and Elena Zavala for helpful discussions and comments on the manuscript. Q.F. was supported by funding from the Chinese Academy of Sciences (XDB26000000), and the National Natural Science Foundation of China (91731303, 41925009,41630102). A.R. was funded by Spanish government (MICINN/FEDER), grant number CGL2016-75109-P. The reassessment of the Spy collection by H.R., I.C. and P.S. was supported by the Belgian Science Policy Office (BELSPO 2004-2007, MO/36/0112). M.S., M.K. and A.D. were supported by the Russian Foundation for Basic Research (RFBR 17-29-04206). This study was funded by the Max Planck Society and the European Research Council (grant agreement number 694707). M.P., J.K. analyzed data. M.H., Q.F., E.E. performed laboratory experiments. H.R., I.C., P.S., L.V.G., V.B.D., C.L.-F., M.d.l.R., A.R., M.V.S., M.B.K., A.P.D. provided samples. B.V., M.M., J.K. supervised the project. M.P., J.K. wrote and edited the manuscript with input from all co-authors.

## Data and materials availability

All sequence data are available from the European Nucleotide Archive under accession numbers xxx (*Mezmaiskaya 2*), xxx (*Spy 94a*), xxx (*Denisova 4*), xxx (*Denisova 8*) and xxx (*El Sidrón 1253*).

Complete source code for data processing and simulations, as well as Jupyter notebooks with all analyses and results can be found at https://github.com/bodkan/archaic-ychr. All data is available from https://bioinf.eva.mpg.de/archaic-ychr.

## Supplementary Information

### 1. Y chromosome DNA capture design

To design a set of DNA capture probes, we identified regions of the human Y chromosome that are uniquely mappable with short sequence reads. Starting from the entire human Y chromosome reference sequence (version *hg19*), we removed regions that overlap those found by the Tandem Repeats Finder (*1*) and those identified by a previously described mappability track as regions that may result in ambiguous alignment of short reads (so called *“map35_50%”* filter, (*2*)). We then removed any regions that were shorter than 99 bp of continuous sequence. In total, this process yielded 6,912,728 bp (∼6.9 Mb) of the Y chromosome suitable for use as an ancient DNA capture target.

We designed 52 bp oligonucleotide probes by tiling the identified 6.9 Mb of target sequence with 52 bp fragments in steps of 3 bp. This resulted in 2,049,846 individual oligonucleotide probes. To verify that the probe sequences are unique genome-wide, we aligned each probe to the complete hg19 reference sequence and confirmed that they all aligned only to their expected position on the Y chromosome with mapping quality of at least 30. The files containing the coordinates of target regions, as well as the coordinates and sequences of all capture probes, including 8 bp adapters, are freely available from https://bioinf.eva.mpg.de/archaic-ychr.

Following the approach taken by Fu *et al*. (*3*), 60 bp oligonucleotides containing the probe sequences as well as an 8 bp universal linker sequence were synthesized on three One Million Feature Arrays (Agilent Technologies), converted into probe libraries and amplified. Single-stranded biotinylated DNA probes were generated using a linear amplification reaction with a single biotinylated primer (*3*).

We also co-analyzed data from two additional captures carried out previously: (i) ∼120 kb of Y chromosome sequence from the *El Sidrón 1253* Neandertal that was targeted as a part of an exome capture study (*4*) and has been analyzed previously (*5*), and (ii) a larger amount of data (∼560 kb) from the same *El Sidrón 1253* individual which we captured using probes designed for a previously published set of Y chromosome target regions (*6*). The file containing the coordinates of target regions and coordinates and sequences of all capture probes, including 8 bp adapters, is freely available from http://bioinf.eva.mpg.de/archaic-ychr.

The features of our new capture design, as well as a comparison with the Y chromosome target regions on the exome capture (*4, 5*) and the ∼560 kb capture (*6*) are reported in Table S1.1.

**Table S1.1.**
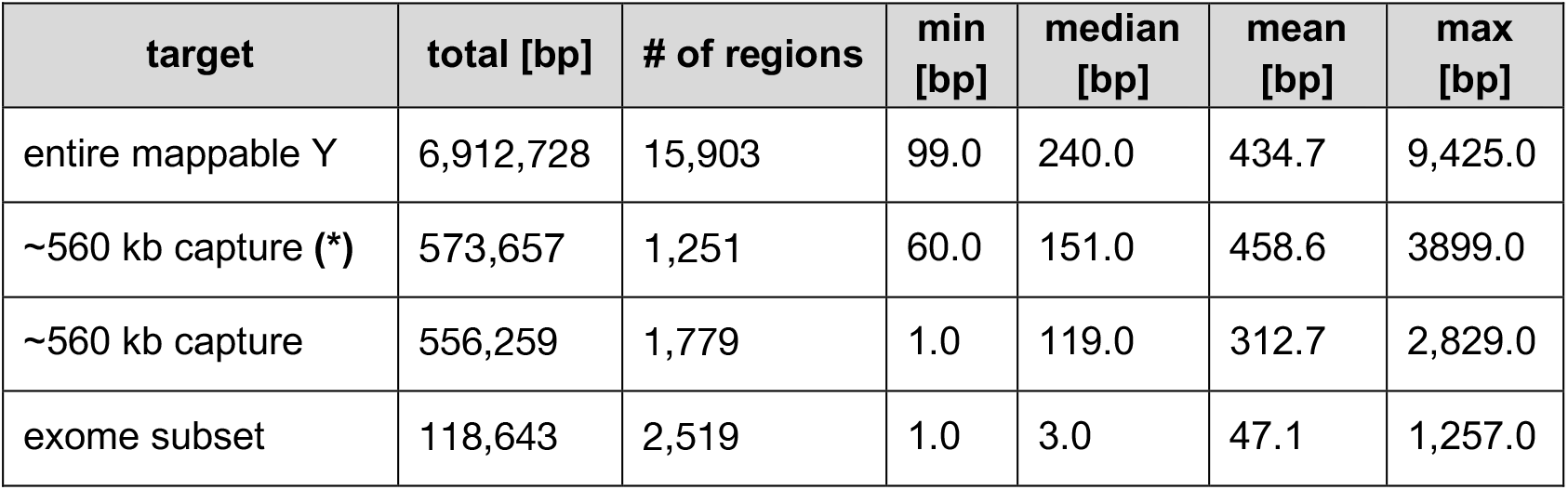
Characteristics of the three sets of Y chromosome capture targets analyzed in our study. “Exome subset” refers to a Y chromosome subset of the exome capture sequence generated by Castellano *et al.* and analyzed by Mendez *et al.* (called “filter 1”) (*4, 5*), *“∼560 kb capture”* refers to target regions originally designed for studying present-day human Y chromosome variation (*6*), star (*) signifies statistics before intersecting the original set of target regions with the *“map35_50%”* filter (*2*), *“entire mappable Y”* represents capture regions targeting the entire mappable portion of the human Y chromosome designed for our study.

| target | total [bp] | # of regions | min [bp] | median [bp] | mean [bp] | max [bp] |
| --- | --- | --- | --- | --- | --- | --- |
| entire mappable Y | 6,912,728 | 15,903 | 99.0 | 240.0 | 434.7 | 9,425.0 |
| ~560 kb capture (*) | 573,657 | 1,251 | 60.0 | 151.0 | 458.6 | 3899.0 |
| ~560 kb capture | 556,259 | 1,779 | 1.0 | 119.0 | 312.7 | 2,829.0 |
| exome subset | 118,643 | 2,519 | 1.0 | 3.0 | 47.1 | 1,257.0 |

### 2. Sampling, DNA extraction, library preparation and capture

Samples of 15.4 mg and 14.9 mg of tooth powder from *Denisova 8* were used for DNA extraction using a silica-based method (*7*) with modifications as described in (*8*). Ten mg of the tooth powder from *Denisova 4* were used for a silica-based DNA extraction that is optimized for the recovery of extremely short DNA fragments (*9*). Four samples of Mezmaiskaya 2 bone powder, ranging between 3.2 mg and 17.5 mg were treated with 0.5% hypochlorite solution to minimize microbial and present-day human DNA contamination (*8*) before DNA was extracted either manually (*7*) or on an automated liquid handling platform (Bravo NGS workstation B, Agilent Technologies) (*10*). See Table S2.1 for an overview of the DNA extracts and libraries generated in this and previous studies and the experimental conditions used.

In addition to existing single-stranded libraries for *Spy94a* and *Mezmaiskaya 2* (*11*), new single-stranded DNA libraries for *Mezmaiskaya 2*, *Denisova 4* and *Denisova 8* were prepared from DNA extracts made for this study (Table S2.1). Two of the single-stranded DNA libraries for *Denisova 8* (A9461 and A9462) were prepared manually using 10 µL of each extract as an input (*12*). All other single-stranded DNA libraries were prepared using either 10 µL or 30 µL of extract as input (*13*) on an automated liquid handling platform (Bravo NGS workstation B, Agilent Technologies) (*14*). All new libraries were prepared without UDG treatment (non-UDG treated libraries).

In order to monitor the efficiency of library preparation, a control oligonucleotide was spiked into each aliquot of a DNA extract used for library preparation (*9*). Quantitative PCR was used to determine the total number of unique library molecules and the number of oligonucleotides that were successfully converted to library molecules (*9, 13*) (Table S2.1). Each library was tagged with two unique index sequences (*15*) and amplified into plateau with AccuPrime Pfx DNA polymerase (Life Technologies) (*16*) according to the modifications detailed in (*8*). Fifty microlitres (half of the total volume) of each of the amplified libraries were purified on an automated liquid handling platform (Bravo NGS workstation B, Agilent Technologies) using SPRI beads (*14*). A NanoDrop 1000 Spectrophotometer (NanoDrop Technologies) was used to determine the concentrations of the purified libraries.

In solution hybridization capture of the Y chromosome was performed in two successive rounds of capture as described previously (*3*), using the Y chromosome probe set designed in the present study and single-stranded libraries prepared in this and previous studies. In addition, we performed hybridization capture on 40 double-stranded libraries prepared in a previous study from the *El Sidrón 1253* Neandertal (see Table S1 in (*4*)) using a smaller ∼560 kb Y chromosomal probe set that was also designed previously (*6*).

### 3. Sequencing and data processing

#### 3.1. Newly generated archaic human Y chromosomes

All captured libraries were sequenced on the Illumina HiSeq 2500 platform in a double index configuration (2×76 cycles) (*15*), and base calling was done using Bustard (Illumina). Adapters were trimmed and overlapping paired-end reads were merged using *leeHom* (*17*). The Burrows-Wheeler Aligner (BWA) (*18*) with parameters adjusted for alignment of ancient DNA (`“-n 0.01 -o 2 -l 16500”`) was used to align the sequenced fragments to the human reference genome version hg19/GRCh37. Only reads showing perfect matches to the expected index sequence combinations were retained for subsequent analyses. PCR duplicates were removed using the *bam-rmdup* program, which can be downloaded in source form from https://github.com/mpieva/biohazard-tools. DNA fragments that were at least 35 base pairs (bp) long and had a mapping quality of at least 25 were extracted using *samtools* (*19*). Each processed and filtered BAM file (one for each archaic human Y chromosome) was intersected with a BED file of the appropriate Y chromosome target (full ∼6.9 Mb capture, ∼120 kb exome capture or ∼560 kb capture).

#### 3.2. Previously published archaic human sequences

In addition to the new capture data generated here, we analyzed previously published shotgun sequences of the *Spy 94a* and *Mezmaiskaya 2* individuals (*11*), as well as exome capture data of the *El Sidrón 1253* individual (*4*). For comparisons to our capture data, we generated BAM files for *Spy 94a* and *Mezmaiskaya 2* shotgun sequences and the *El Sidrón 1253* exome capture by filtering the published data to minimum read length of 35 bp and mapping quality 25, keeping only sequences aligned to the set of appropriate target capture regions (∼6.9 Mb capture target for *Spy 94a* and *Mezmaiskaya 2*, ∼118 kb capture target for *El Sidrón 1253, Table S1.1).*

#### 3.3. Previously published modern human sequences

For comparisons with modern human Y chromosomes, we downloaded 19 BAM files of African and non-African Y chromosomes published by the Simons Genome Diversity Project (SGDP) (*20*), two Y chromosomes representing the African A00 lineage (*21*) and the Y chromosome of a ∼45,000-year-old hunter-gatherer *Ust’-Ishim* (*22*). Because the two individuals from which the A00 Y chromosomes were sequenced are closely and each is only about half of the coverage of the other modern human Y chromosomes (Table S4.3), we followed the approach of the original A00 publication and merged the two A00 Y chromosomes into a single BAM file (*21, 23*). All individual BAM files (one for each modern human Y chromosome) were then filtered to retain reads with a minimum length of 35 bp and mapping quality of at least 25, and alignment to the appropriate set of Y chromosome target capture regions (Table S1.1).

### 4. Coverage and measures of ancient DNA quality

#### 4.1. Coverage

Sequencing coverage was calculated using *bedtools* (*24*). To get coverage for a given individual in a given set of target regions, we ran the command `bedtools coverage - a <BED> -b <BAM> -d`, which reports the coverage for each position in a BED file in the last column of its output. We removed sites with coverage higher than the 98% quantile of the entire distribution in each of the individuals in our study. Figure 1B (spatial distribution and overall distribution) and Tables S4.1, S4.2 and S4.3 summarise the values of coverage at sites with less than 98% quantile of the overall distribution in a sample.

**Table S4.1.**
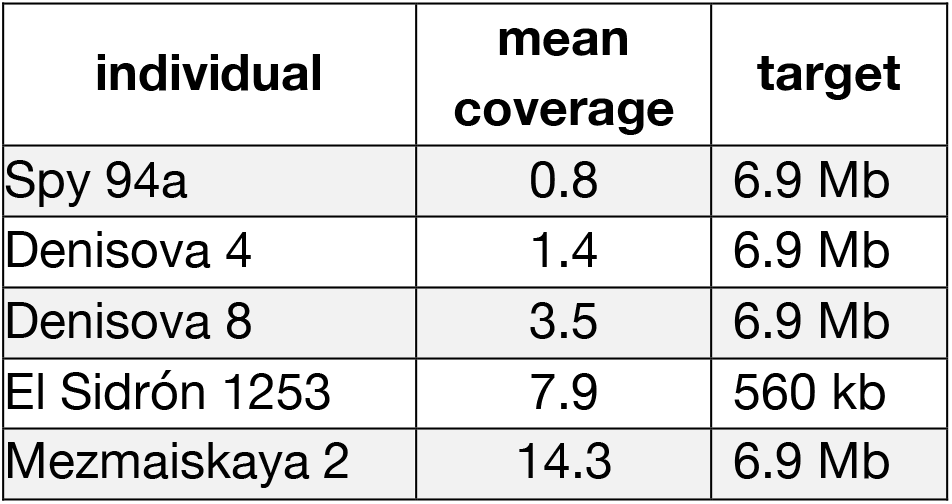
Mean coverage of archaic human Y chromosomes sequenced in this study. Sites with coverage higher than 98% quantile of the entire distribution were excluded from the calculation.

**Table S4.2.** Mean coverage of previously published archaic human Y chromosome sequences. The coverage reported for *Spy 94a* and *Mezmaiskaya 2* shotgun sequence data is that of sequences overlapping the 6.9 Mb Y capture regions. The *El Sidrón 1253* libraries were captured using an exome capture array (*4, 5*) and the coverage reported here is for the ∼118 kb exome target capture regions. For each individual, sites with coverage higher than 98% quantile of the entire distribution were excluded from the calculation.

| name | mean coverage | target | study |
| --- | --- | --- | --- |
| Spy 94a (shotgun) | 0.5 | 6.9 Mb | Hajdinjak *et al.*, 2018 |
| Mezmaiskaya 2 (shotgun) | 0.8 | 6.9 Mb | Hajdinjak *et al.*, 2018 |
| El Sidrón 1253 (capture) | 3.2 | 118 kb | Castellano *et al.*, 2014, Mendez *et al.*, 2016 |

**Table S4.3.** Mean coverage of modern human Y chromosomes in capture target regions. Coverage is reported using sequences within the 6.9 Mb target capture regions. For each individual, sites with coverage higher than 98% quantile of the entire distribution were excluded from the calculation.

| name | mean coverage | target | study |
| --- | --- | --- | --- |
| A00-1 | 8.8 | 6.9 Mb | Karmin *et al.*, 2015 |
| A00-2 | 12.0 | 6.9 Mb | Karmin *et al.*, 2015 |
| S_Mandenka-1 | 16.3 | 6.9 Mb | Mallick *et al.*, 2016 |
| S_Yoruba-2 | 17.0 | 6.9 Mb | Mallick *et al.*, 2016 |
| S_Finnish-2 | 17.0 | 6.9 Mb | Mallick *et al.*, 2016 |
| S_Punjabi-1 | 17.1 | 6.9 Mb | Mallick *et al.*, 2016 |
| S_Sardinian-1 | 18.1 | 6.9 Mb | Mallick *et al.*, 2016 |
| S_Dai-2 | 19.8 | 6.9 Mb | Mallick *et al.*, 2016 |
| S_Gambian-1 | 20.1 | 6.9 Mb | Mallick *et al.*, 2016 |
| Ust'-Ishim | 20.1 | 6.9 Mb | Fu *et al.*, 2014 |
| S_Mbuti-1 | 20.3 | 6.9 Mb | Mallick *et al.*, 2016 |
| S_Dinka-1 | 20.8 | 6.9 Mb | Mallick *et al.*, 2016 |
| S_Han-2 | 20.8 | 6.9 Mb | Mallick *et al.*, 2016 |
| A00 | 20.9 | 6.9 Mb | merged A00-1 and A00-2 |
| S_BedouinB-1 | 21.7 | 6.9 Mb | Mallick *et al.*, 2016 |
| S_French-1 | 21.9 | 6.9 Mb | Mallick *et al.*, 2016 |
| S_Karitiana-1 | 22.2 | 6.9 Mb | Mallick *et al.*, 2016 |
| S_Turkish-1 | 22.5 | 6.9 Mb | Mallick *et al.*, 2016 |
| S_Saami-2 | 22.6 | 6.9 Mb | Mallick *et al.*, 2016 |
| S_Ju_hoan_North-1 | 22.7 | 6.9 Mb | Mallick *et al.*, 2016 |
| S_Papuan-2 | 23.2 | 6.9 Mb | Mallick *et al.*, 2016 |
| S_Thai-1 | 25.1 | 6.9 Mb | Mallick *et al.*, 2016 |
| S_Burmese-1 | 29.2 | 6.9 Mb | Mallick *et al.*, 2016 |

#### 4.2. Patterns of ancient DNA damage

To check for the presence of genuine ancient DNA sequences, we looked for an increased rate of deamination-induced substitutions, an important signature of ancient DNA damage (*25*). We counted substitution frequencies for each individual BAM file (one BAM file per individual Y chromosome) and found that molecules from single-stranded libraries that were not treated by uracil-DNA glycosylase (UDG) enzyme (those from *Spy 94a*, *Mezmaiskaya 2*, *Denisova 4* and *Denisova 8*) show highly elevated frequencies of C-to-T substitutions towards the ends of molecules, as well as C-to-T substitutions throughout the molecules (Figure S4.1). As is characteristic of double-stranded libraries treated with the UDG enzyme, the deamination substitution frequency signal in the capture data from *El Sidrón 1253* UDG-treated libraries is much less pronounced and present only at the terminal positions of DNA fragments as both C-to-T and G-to-A substitutions (Figure S4.2). For comparison, Figure S4.3 shows DNA damage patterns from previously published shotgun sequences of *Spy 94a* and *Mezmaiskaya 2* individuals.

**Figure S4.1.**
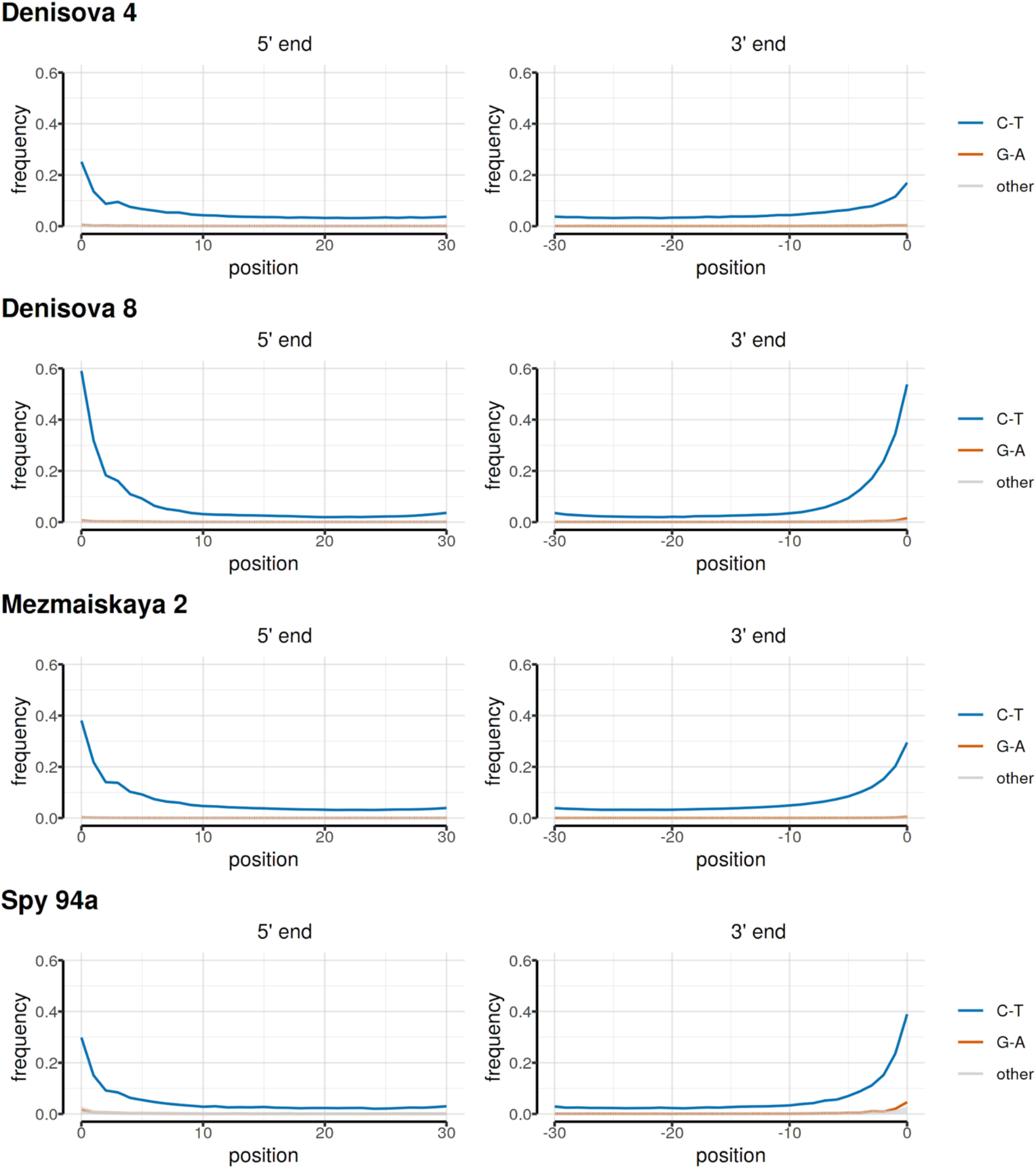
Patterns of ancient DNA damage in non-UDG-treated sequences captured using the 6.9 Mb capture.

**Figure S4.2.**
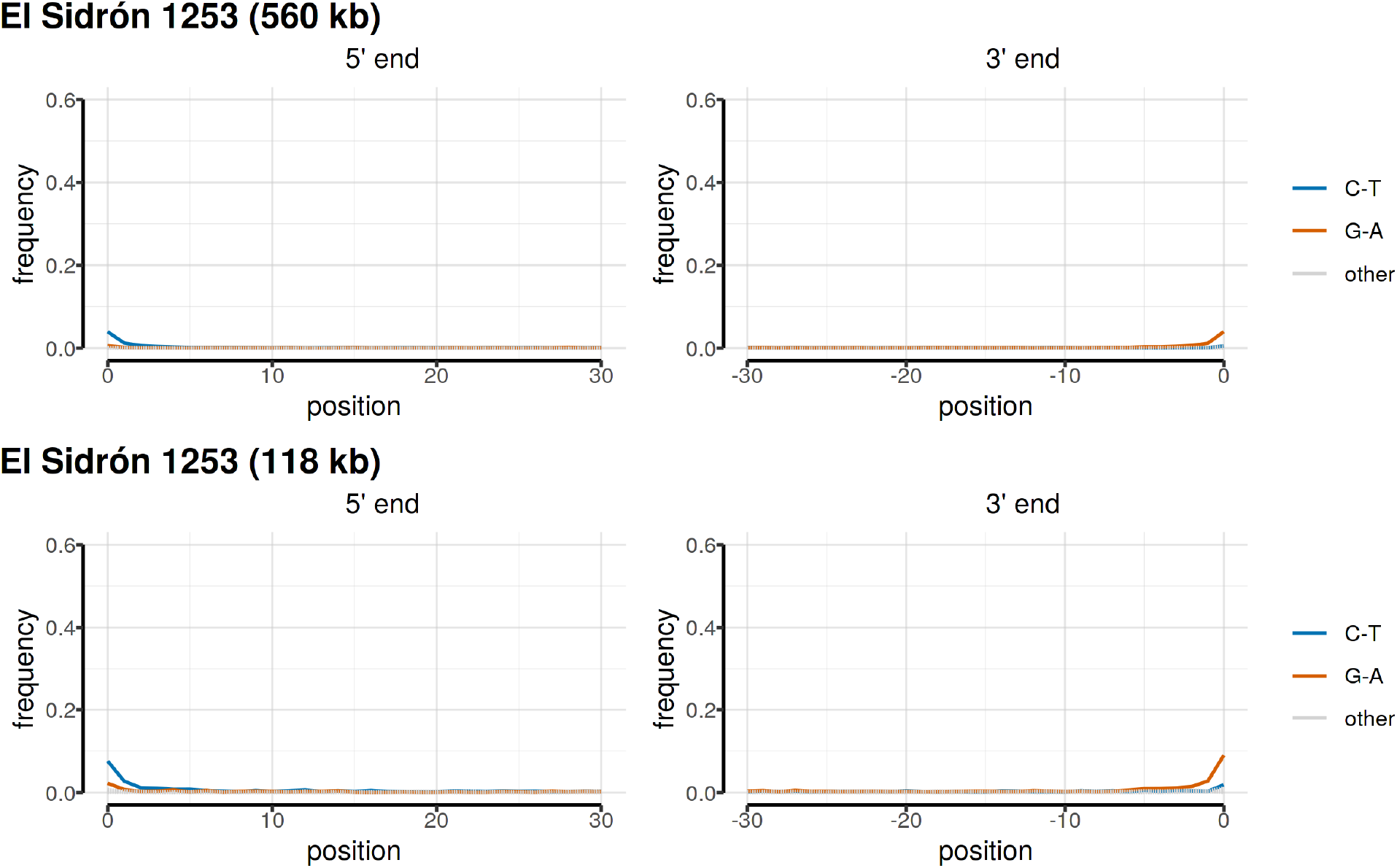
Patterns of ancient DNA damage in UDG-treated sequences from the *El Sidrón 1253* individual. Top row shows deamination patterns in the 560 kb capture generated for our study (SI 1), bottom row shows deamination patterns in previously published Y chromosome sequences from the exome capture of the same individual (*4, 5*).

**Figure S4.3.**
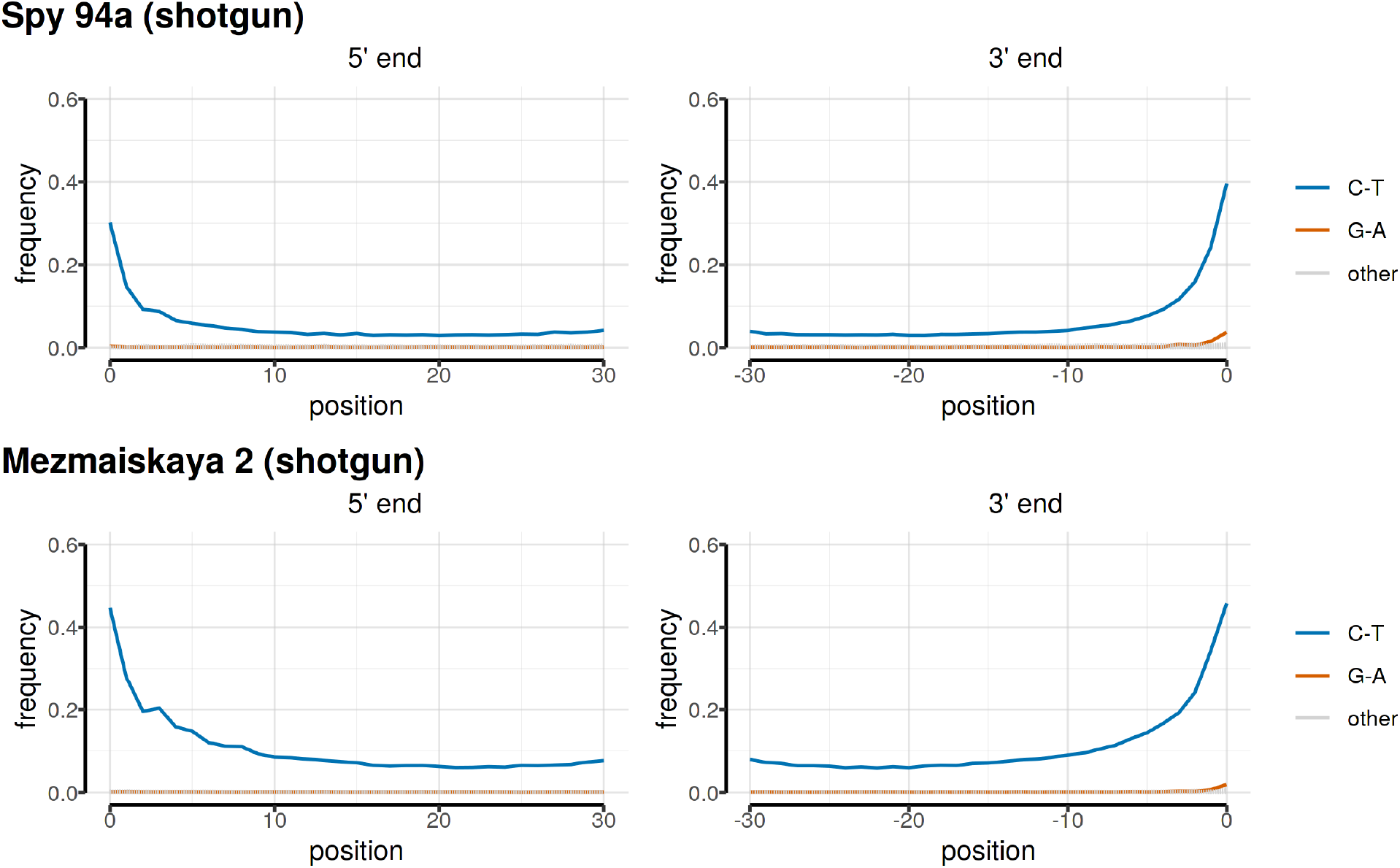
Patterns of ancient DNA damage in non-UDG-treated shotgun sequences of *Spy 94a* and *Mezmaiskaya 2*. Figures show results from previously published sequence data (*11*), using sequences within the 6.9 Mb Y chromosome capture target.

#### 4.3. Read length distribution

We calculated read lengths for each final processed BAM file using *samtools view* and *awk*. As expected for ancient sequences, archaic human Y chromosome fragments are very short (Figure S4.3, Table S4.4). We note that *Denisova 8* shows an even more extreme reduction in read length compared to the other captured archaic human Y chromosomes (Figure S4.3, Table 4.4), consistent with the fact that the *Denisova 8* specimen is possibly nearly twice as old as the other archaic humans in our study (Figure 1A).

**Figure S4.3.**
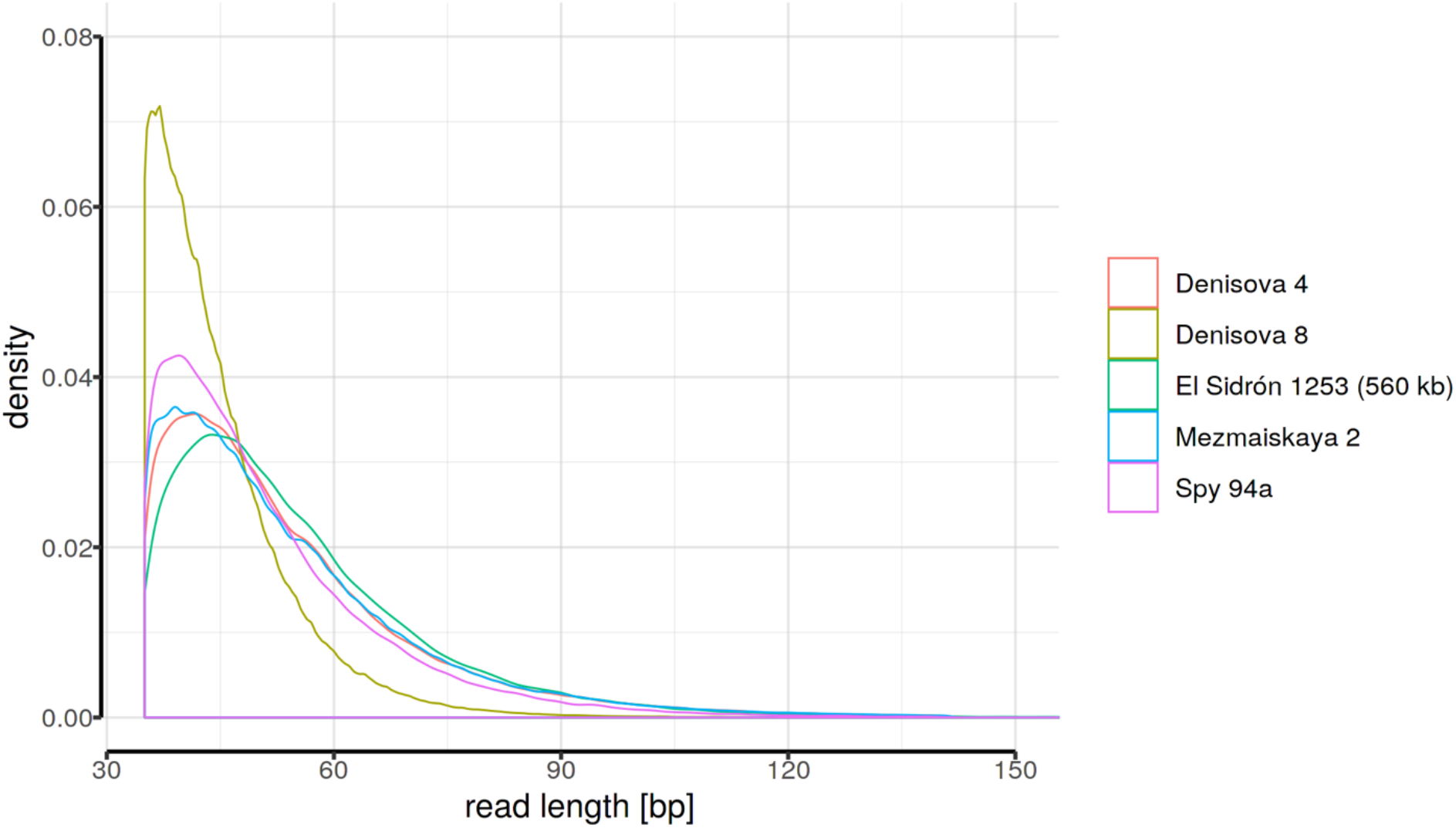
Distributions of read lengths calculated using sequences within the 6.9 Mb target regions.

**Table S4.4.**
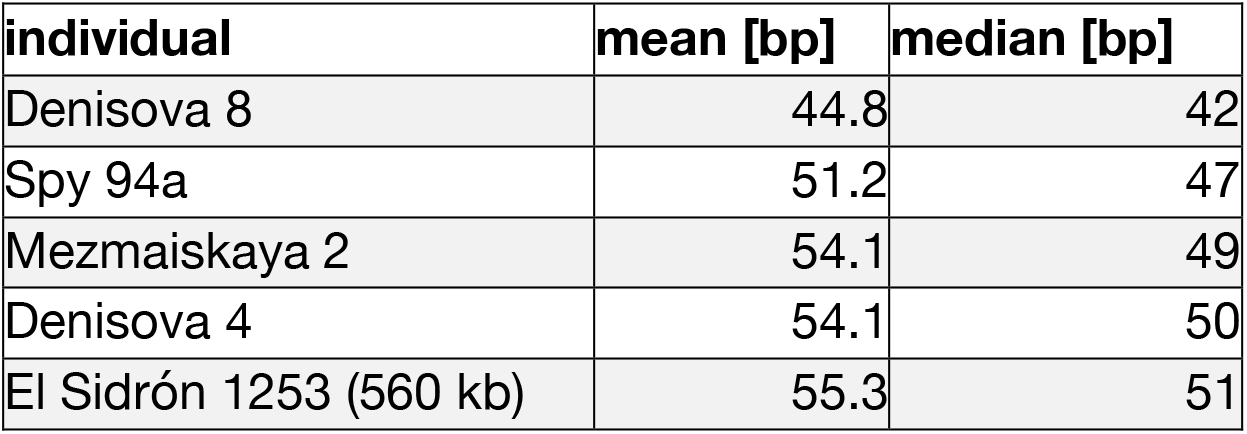
Mean and median values of read length distributions in Figure S4.3.

#### 4.4. Modern human contamination

Our consensus-based genotype calling strategy (described in SI 5) is designed to remove the effect of modern human contamination under the assumption that contaminant reads at a given position never represent more than 90% of the total number of reads. To validate that this approach achieves the desired effect, we assessed the frequency of modern human-derived SNPs at positions informative about modern human contamination in the final archaic human Y chromosome genotype calls.

To define these informative positions, we used genotypes of present-day human Y chromosomes and identified ancestral states by determining which sites carry the same allele in chimpanzee and two present-day African lineages *A00* and *S_Ju_hoan_North-1* (*20, 21*) (red branches in Figure S4.4). We then further restricted these sites to those in which a different allele is observed in all 13 non-African individuals from the SGDP panel (*20*). These represent alleles derived on the non-African Y chromosome lineage (blue branches in Figure S4.4). This conditioning led to a total of 268 informative positions. Given that all archaic human Y chromosomes are expected to carry the ancestral state at these sites because they all fall basal to modern human Y chromosomes (Figure 2A), observing a derived allele at any of these informative sites implies the presence of a modern human contaminant allele, double mutation or an erroneous SNP call. We note that although the 13 non-African Y chromosomes that we used to define the potential ‘contaminant-derived states’ may not represent the true contaminant population, the contaminating population would still share the same derived states due to the non-recombining nature of human Y chromosomes.

Using this set of 268 informative positions, we found that the five archaic human Y chromosomes carry the ancestral state at all informative positions except for a single position in the *Spy 94a* individual which shows a derived allele out of the total 16 informative sites available (Table S4.5). This shows that the consensus genotype calling method is efficient in mitigating the effect of modern human contaminant reads on the final set of Y chromosome genotype calls.

**Figure S4.4.**
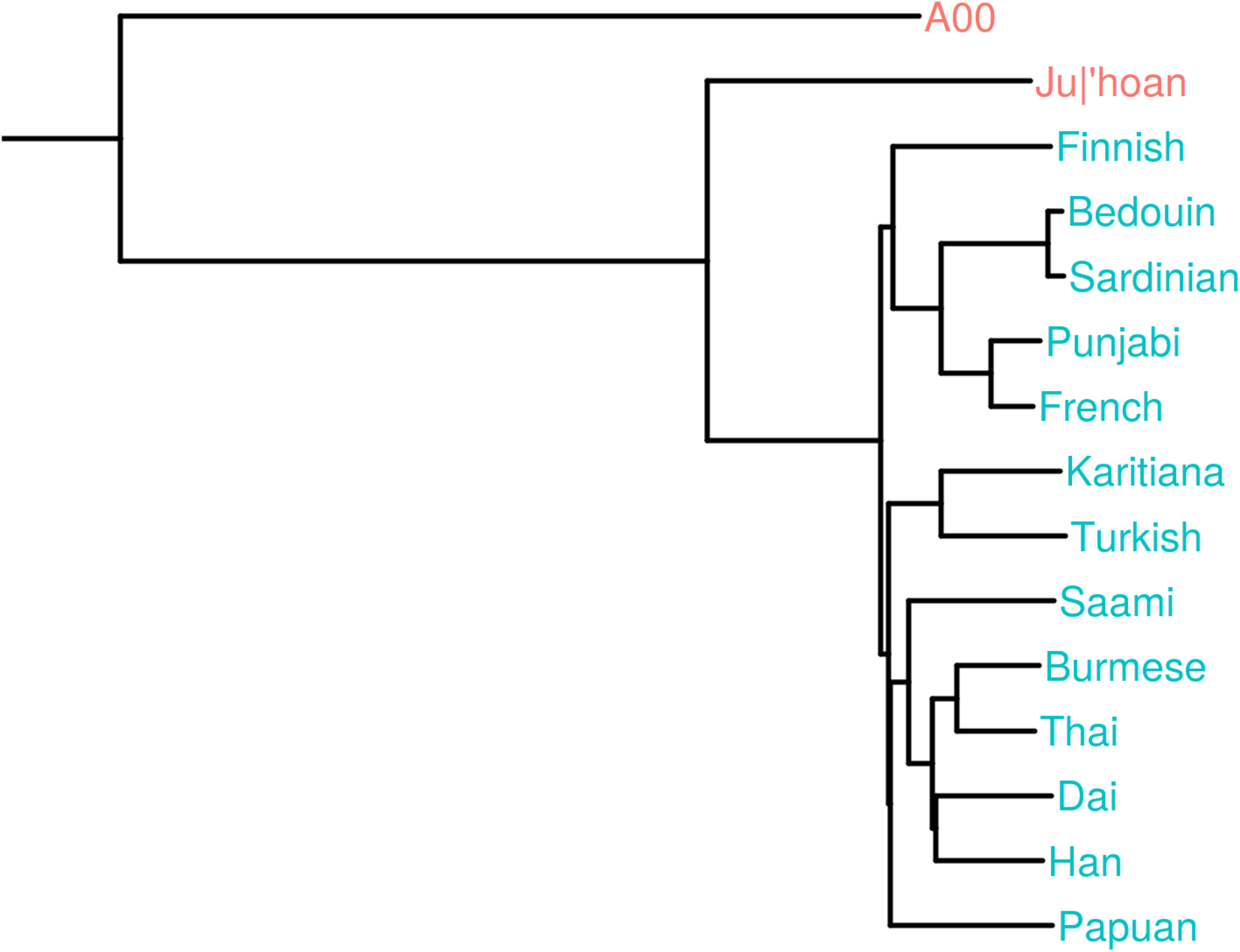
Phylogenetic tree demonstrating the definition of positions informative about modern human contamination. Red branches represent lineages which we required to carry one state (together with the chimpanzee) at positions where the blue lineages carry a different state. Therefore, red branches represent the ancestral states and blue branches represent the derived state. The tree was rooted with a chimpanzee Y chromosome as an outgroup (cropped for plotting purposes).

**Table S4.5.** Counts and proportions of potential ‘contaminant-derived’ non-African alleles in all archaic human Y chromosomes.

| individual | ancestral<br>count | derived<br>count | total | derived/total |
| --- | --- | --- | --- | --- |
| Spy 94a | 15 | 1 | 16 | 0.0625 |
| Mezmaiskaya 2 | 189 | 0 | 189 | 0.0000 |
| Denisova 4 | 14 | 0 | 14 | 0.0000 |
| Denisova 8 | 90 | 0 | 90 | 0.0000 |
| El Sidrón 1253 (560 kb) | 29 | 0 | 29 | 0.0000 |

### 4.5. Capture bias and reference (mapping) bias

Because the probes we designed for Y chromosome DNA enrichment are based on the human reference genome sequence (SI 1), we were concerned about the effect of capture bias on our inferences, specifically on the observed differences in divergence times between Denisovan and Neandertal Y chromosomes with respect to present-day humans (Figure 2). An earlier study of reference bias in published aDNA data sets has found minor but significant allelic imbalances at heterozygous sites from a baseline expectation of 50% ratio between reference and alternative alleles (*26*). Because this approach is not applicable for haploid Y chromosomes, we instead looked for departures of the observed number of sites without any genomic coverage from the theoretical expectation.

To build an intuition about this expectation, let’s first consider a case of a truly random distribution of sequencing reads in a complete absence of capture or reference bias. In such a situation, the count of reads observed at any site can be modeled as a random variable which follows a Poisson distribution with a parameter λ, where λ represents the average coverage observed across all sites. In a mathematical notation, letting *X* be this count of reads: *X∼ Poisson*(λ). Then, given some value of λ, the expected proportion of sites that are not covered by any sequencing reads can be expressed as *Poisson*(*X* = 0,*λ*), i.e. as the probability of observing zero reads at any site given the overall average coverage of λ. As an example, assuming 1-fold sequencing coverage we would expect to see (*X* = 0, λ = 1) = 0.3678794 ∼ 37% of the target sites not to be covered by any read at all, just by random chance. Importantly, however, capture bias or reference bias will manifest by some regions of the genome being underrepresented in terms of captured molecules or mapped reads. Therefore, the presence and magnitude of this bias in a given DNA enrichment experiment can be detected by estimating the difference between the proportion of sites without any sequencing coverage from the theoretical Poisson expectation.

The results, shown in Figure S4.5 and Table S4.6, demonstrate that there is both reference and capture bias in our data and offers several interesting insights. First, we see a comparable effect of bias in all capture data (4-6% departure from the theoretical Poisson expectation) regardless of which capture array was used for the enrichment procedure (i.e., the full 6.9 Mb capture array, the 560 kb capture array or the exome capture array, Figure S4.5). Furthermore, comparisons of capture and shotgun sequences of *Spy 94a* and *Mezmaiskaya 2* show that the majority of bias must be due to the capture procedure itself (i.e. failure to capture molecules). This is because the underlying true biological divergences of *Spy 94a* and *Mezmaiskaya 2* to the reference genome (which cause a failure to map reads due to an increased number of substitutions – i.e., a reference bias) must be the same for both capture and shotgun sequences from these individuals. Crucially, however, despite the differences in bias between capture and shotgun sequences, both datasets lead to the same estimates of TMRCA with present-day human Y chromosomes (Figure S7.11). Furthermore, although we see dramatically different phylogenetic relationships of Denisovan and Neandertal Y chromosomes with respect to modern humans (Figure 2A), both groups of archaic human capture sequences display comparable magnitudes of both sources of bias (Figure S4.5). Therefore, although undoubtedly present, capture and reference biases cannot result in the observed differences in divergence times between archaic human Y chromosomes and present-day human Y chromosomes (Figure 2).

**Figure S4.5.**
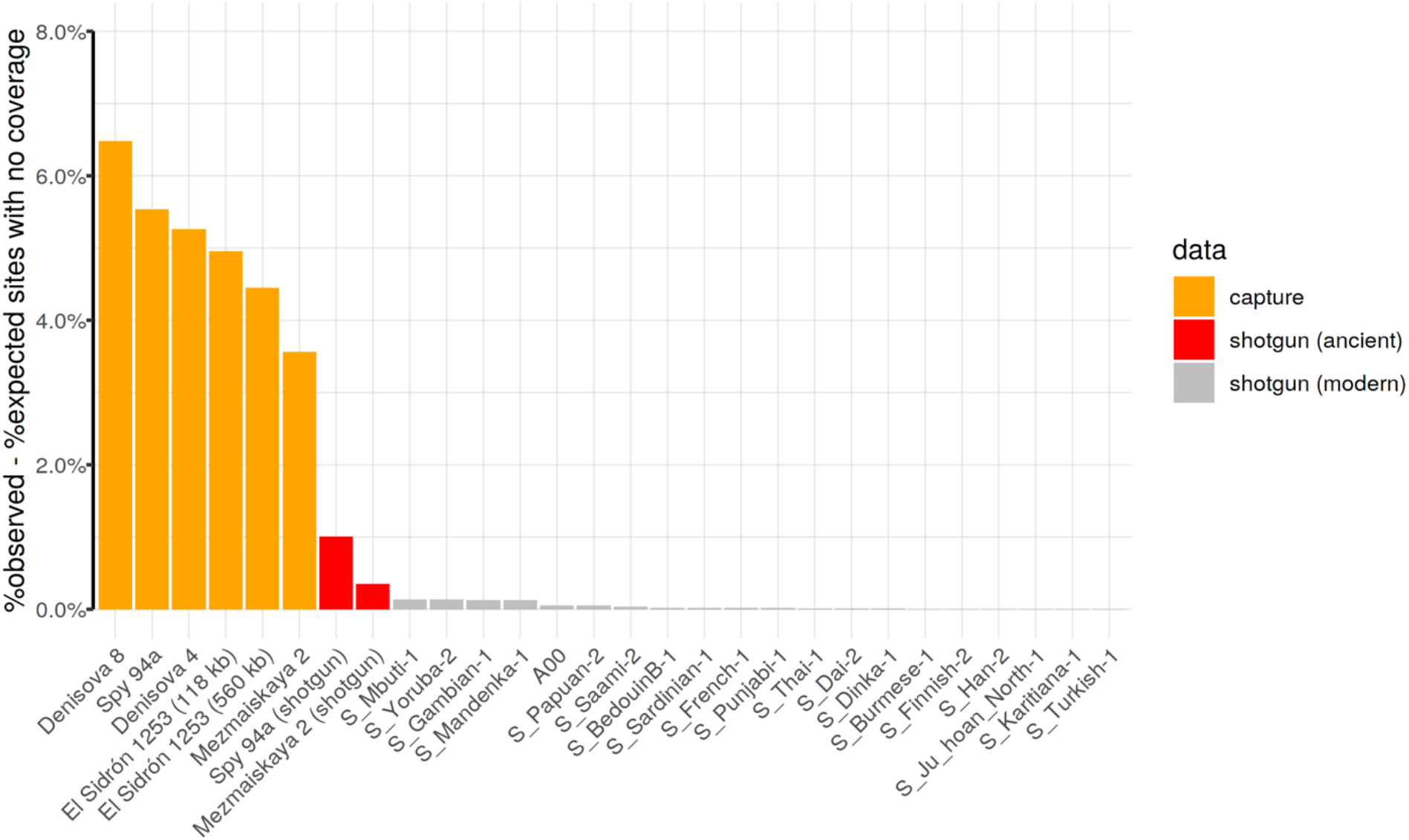
Differences between expected and observed counts of sites without any sequencing coverage. Exact counts are reported in Table S4.6. Data were filtered according to the criteria described in SI 4.1.

**Table S4.6.** Proportions of expected and observed sites without any coverage.

| name | coverage | observed | expected | difference | data |
| --- | --- | --- | --- | --- | --- |
| Denisova 8 | 3.4820355 | 0.09555 | 0.03074 | 0.06481 | capture |
| Spy 94a | 0.8253622 | 0.49341 | 0.43808 | 0.05533 | capture |
| Denisova 4 | 1.3717806 | 0.30622 | 0.25365 | 0.05257 | capture |
| El Sidrón 1253 (118 kb) | 3.2121215 | 0.08978 | 0.04027 | 0.04951 | capture |
| El Sidrón 1253 (560 kb) | 7.9165032 | 0.04479 | 0.00036 | 0.04443 | capture |
| Mezmaiskaya 2 | 14.3494002 | 0.03561 | 0.00000 | 0.03561 | capture |
| Spy 94a (shotgun) | 0.5052873 | 0.61337 | 0.60333 | 0.01004 | shotgun (ancient) |
| Mezmaiskaya 2 (shotgun) | 0.8248822 | 0.44174 | 0.43829 | 0.00345 | shotgun (ancient) |
| S_Yoruba-2 | 16.9616115 | 0.00129 | 0.00000 | 0.00129 | shotgun (modern) |
| S_Mbuti-1 | 20.3360989 | 0.00129 | 0.00000 | 0.00129 | shotgun (modern) |
| S_Mandenka-1 | 16.3312730 | 0.00127 | 0.00000 | 0.00127 | shotgun (modern) |
| S_Gambian-1 | 20.1210272 | 0.00127 | 0.00000 | 0.00127 | shotgun (modern) |
| A00 | 20.8671570 | 0.00050 | 0.00000 | 0.00050 | shotgun (modern) |
| S_Papuan-2 | 23.1529894 | 0.00048 | 0.00000 | 0.00048 | shotgun (modern) |
| S_Saami-2 | 22.6298230 | 0.00033 | 0.00000 | 0.00033 | shotgun (modern) |
| S_BedouinB-1 | 21.7140764 | 0.00016 | 0.00000 | 0.00016 | shotgun (modern) |
| S_Sardinian-1 | 18.1189362 | 0.00015 | 0.00000 | 0.00015 | shotgun (modern) |
| S_Punjabi-1 | 17.1469680 | 0.00013 | 0.00000 | 0.00013 | shotgun (modern) |
| S_French-1 | 21.9031161 | 0.00013 | 0.00000 | 0.00013 | shotgun (modern) |
| S_Thai-1 | 25.1203313 | 0.00010 | 0.00000 | 0.00010 | shotgun (modern) |
| S_Dai-2 | 19.8120096 | 0.00004 | 0.00000 | 0.00004 | shotgun (modern) |
| S_Dinka-1 | 20.7590166 | 0.00004 | 0.00000 | 0.00004 | shotgun (modern) |
| S_Finnish-2 | 17.0155033 | 0.00003 | 0.00000 | 0.00003 | shotgun (modern) |
| S_Han-2 | 20.7680004 | 0.00003 | 0.00000 | 0.00003 | shotgun (modern) |
| S_Karitiana-1 | 22.1963029 | 0.00003 | 0.00000 | 0.00003 | shotgun (modern) |
| S_Turkish-1 | 22.4638799 | 0.00003 | 0.00000 | 0.00003 | shotgun (modern) |
| S_Ju_hoan_North-1 | 22.7025717 | 0.00003 | 0.00000 | 0.00003 | shotgun (modern) |
| S_Burmese-1 | 29.2087547 | 0.00003 | 0.00000 | 0.00003 | shotgun (modern) |

### 5. Genotype calling

#### 5.1. Consensus genotype calling

The haploid nature of the human Y chromosome alleviates many issues inherent to genotype calling of diploid genomes. Most importantly, given that only one allele is expected to be present at each site of a non-recombining portion of the Y chromosome, observing more than one allele at a site must be the result of sequencing errors, DNA damage, contamination, or misalignment of reads. While such issues present a significant problem for calling diploid genotypes, by making it challenging to distinguish true heterozygous calls from erroneously called heterozygous genotypes (*27, 28*), they are less of an issue for haploid genotyping.

To call genotypes of the archaic and modern human Y chromosomes in our study, we applied a conservative approach to produce a consensus of sequencing reads. For each Y chromosome BAM file, we performed a pileup of reads at each site (disabling base quality recalibration), filtering out reads with mapping quality less than 25, ignoring bases with base quality less than 20 and removing reads carrying indels at a pileup position. Then, under the assumption that alleles introduced due to DNA damage, sequencing errors, misalignments, or contamination will be in a minority at each site, we called the allele supported by 90% of the reads in a pileup as the haploid genotype for that site. For further analyses, we additionally restricted to genotype calls supported by at least three reads, as described in section 5.3. This genotype calling procedure has been implemented as part of a functionality of a program which is freely available for download at https://github.com/bodkan/bam-caller.

#### 5.2. Genotype calling using snpAD

The consensus genotype calling approach described in the previous section is quite conservative and does not incorporate an explicit model of DNA damage and sequencing errors. To validate the robustness of our consensus-based results, we compared them to genotype calls generated using *snpAD*, an aDNA-specific genotype caller (*28*). A major caveat of this approach is the fact that *snpAD* has been designed for calling diploid genotypes and accurate results requires at least 4X genomic coverage (*28*). Therefore, its genotype calling model has not been tested on low coverage, haploid chromosomes such as those generated in our study. While recognizing these limitations, we used *snpAD* to call genotypes of all four archaic Y chromosomes captured for the 6.9 Mb target regions (*Denisova 4*, *Denisova 8*, *Spy 94a* and *Mezmaiskaya* 2), discarded any sites which were called as heterozygous (likely the result of errors, contamination or aDNA damage), and converted all homozygous genotype calls to a haploid state.

In accordance with *snpAD*’s more sophisticated model of aDNA damage patterns, we have found that the number of successfully genotyped sites is higher than those generated by our simpler consensus-based genotype calling approach, but only marginally so (Table S5.1). Furthermore, although the rates of C-to-T and G-to-A SNP frequencies observed in the final set of genotype calls of the high coverage *Mezmaiskaya 2* is very close to the baseline expectation for present-day DNA, the remaining low coverage archaic Y chromosomes are still affected by aDNA damage and show an excess of falsely called genotypes (Figures S5.1 and S5.2). We note that this is not unexpected, because the coverage of these individuals is much lower than what is recommended by for *snpAD* (*28*). Overall, we did not observe significant differences between *snpAD*-based and consensus-based genotypes in terms of the inferred times to the most recent common ancestor (TMRCA) and, in fact, we found that both lead to the same conclusions (Figure S7.11). Based on these analyses we concluded that our conservative 90% cutoff for consensus genotype calling method is appropriate and decided to use it for all analyses.

#### 5.3. Minimum coverage filtering

The majority of libraries analyzed in our study have not been treated with the uracil-deglycosylase (UDG) enzyme (SI 2). Unlike UDG-treated libraries, non-UDG libraries retain an increased deamination signal throughout the molecules (Figures S4.1 and S4.2) which poses a significant challenge for distinguishing false substitutions caused by aDNA damage from true polymorphisms (*28*).

For a given sequencing read carrying a putative substitution, it is not straightforward to decide whether this substitution represents a true polymorphism or error. Given enough sequencing coverage, this issue can be mostly overcome by observing a sufficient number of bases from reads that do not carry a deamination-induced substitution, integrating evidence from multiple reads at a site (*28*). However, as our data is of relatively low coverage (Figure 1B, Table S4.1), we were concerned by selecting an appropriate lower coverage cutoff to minimize the impact of false polymorphisms on our inferences. Specifically, if the same nucleotide is observed in a majority of reads mapped to the same genomic position, it is unlikely that this would be the result of aDNA damage, sequencing errors or contamination, as these occur mostly at relatively low frequencies in the individuals in our study (Figure S4.1 and Table S4.5). To get a sense of the frequency of calling false polymorphisms as a function of coverage, we calculated the proportions of observed genotypes in each archaic human Y chromosome given a certain coverage filtering cutoff and compared those to the baseline expectation for present-day human DNA. As expected, allowing SNPs supported by only one read leads to a significant excess of C-to-T and G-to-A SNP (Figure S5.1), a consequence of the presence of aDNA damage (Figure S4.1). We found that increasing the minimum coverage cutoff to two reads causes the rate of aDNA-induced SNPs to drop significantly towards the baseline expectation, but going beyond requiring the support of three reads for each SNP does not lead to further improvement in accuracy of genotype calling (Figure S5.1). Because of this and because each additional increase in required minimum coverage is at the expense of the final number of available sites, we settled on a minimum coverage cutoff of 3 reads.

It is important to note that despite the residual presence of false aDNA substitutions in the final set of filtered genotype calls, manifesting as increased frequencies of C-to-T and G-to-A SNPs compared to present-day DNA (Figure S5.1), comparisons of archaic-modern human TMRCA estimates obtained using the full set of genotype calls and those based on genotypes restricted to non-C-to-T/G-to-A SNPs did not reveal any significant differences (Figure S7.9). This is partially due to very low rates of residual false SNPs that pass through the filtering, but mostly because our TMRCA estimators are quite insensitive to private mutations on the archaic lineage (SI 7). A second validation of our coverage filter follows from the fact that the two Denisovan and all three Neandertal Y chromosomes lead to the same TMRCA estimates with modern human Y chromosomes despite differences in coverage and rates of aDNA damage (Figures 1B, 2B and S5.1). Both of these factors would affect genotyping accuracy if not handled appropriately, and would introduce noise in TMRCA estimates estimated for individual Y chromosomes.

Complete counts of Y chromosome positions passing the filters for all individuals are reported in Tables 5.1 and 5.2 (counts for archaic and modern human individuals, respectively, in 6.9 Mb target regions) and Tables 5.3 and 5.4 (counts for archaic and modern human individuals, respectively, in 560 kb target regions).

**Figure S5.1.**
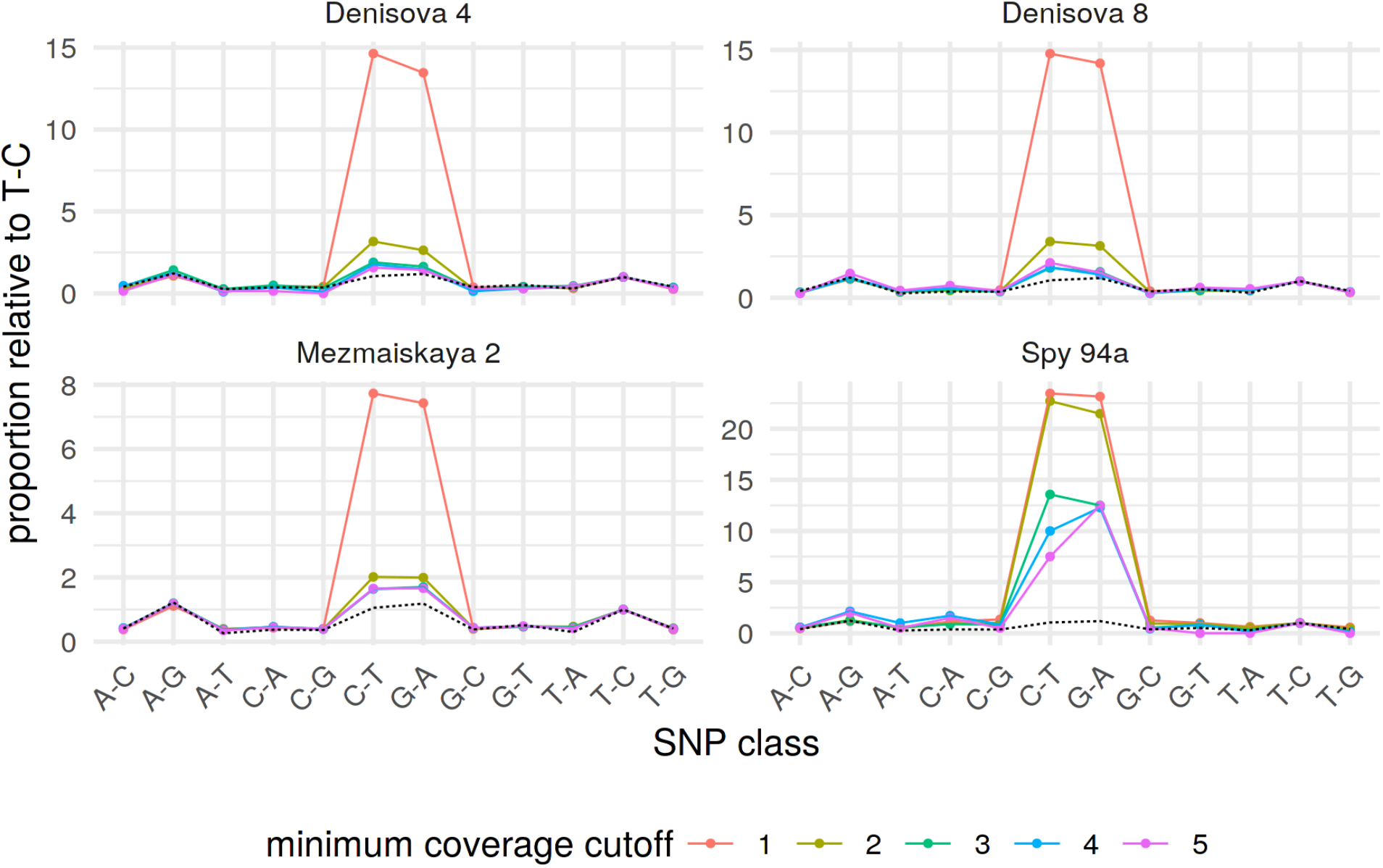
Frequencies of observed polymorphisms normalized by the observed frequency of T-C polymorphism. SNP classes were counted for each archaic human Y chromosome (one panel each) and all counts were normalized by dividing them with observed counts of T-C SNPs. Dotted line shows an expectation based on SNP proportions observed in Y chromosomes of the following SGDP individuals: *S_French_1*, *S_Papuan_2*, *S_Burmese_1*, *S_Thai_1*, *S_Sardinian_1*.

**Figure S5.2.**
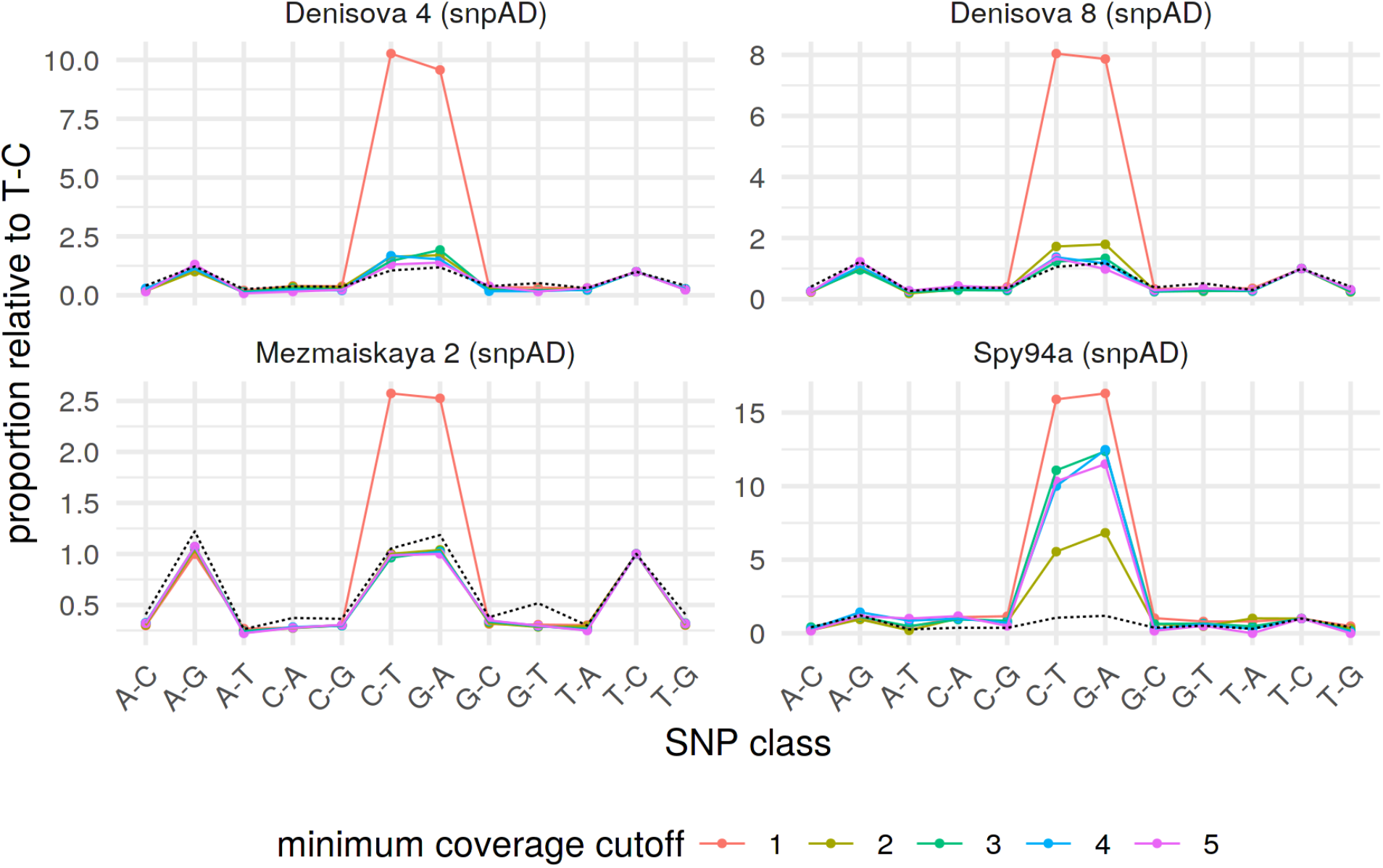
Frequencies of observed polymorphisms normalized by the observed frequency of T-C polymorphism. SNP classes were counted for each archaic human Y chromosome (one panel each) and all counts were normalized by dividing them with observed counts of T-C SNPs. Dotted line shows an expectation based on SNP proportions observed in Y chromosomes of the following SGDP individuals: *S_French_1*, *S_Papuan_2*, *S_Burmese_1*, *S_Thai_1*, *S_Sardinian_1*.

**Figure S5.3.**
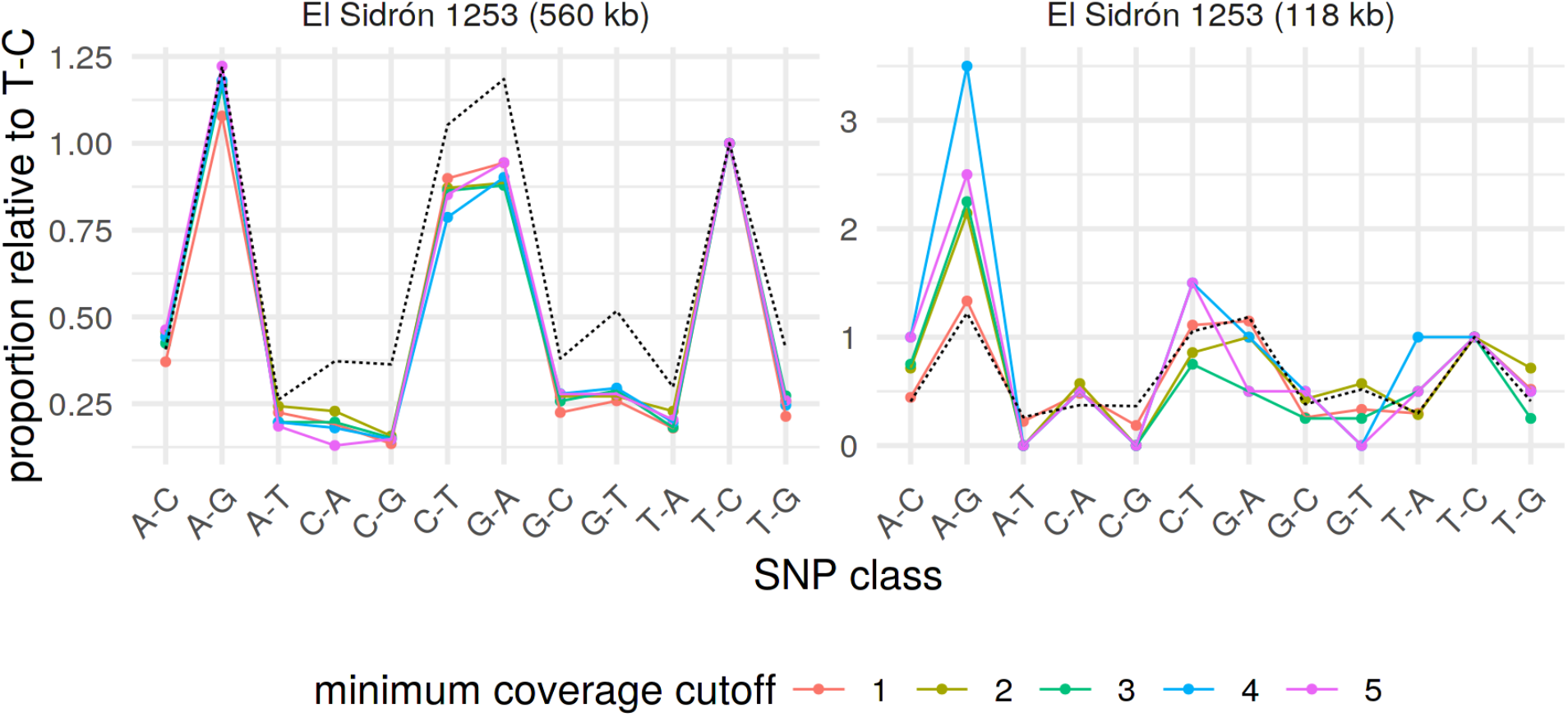
Frequencies of observed polymorphisms normalized by the frequency of T-C polymorphism. SNP classes were counted for each archaic human Y chromosome (one panel each) and all counts were normalized by dividing them with observed counts of T-C SNPs. Dotted line shows an expectation based on SNP proportions observed in Y chromosomes of the following SGDP individuals: *S_French_1*, *S_Papuan_2*, *S_Burmese_1*, *S_Thai_1*, *S_Sardinian_1*.

**Table S5.1.** Counts of sites for each archaic human Y chromosome in 6.9 Mb capture regions which passed the filtering for minimum depth of 3 reads in addition to other filtering and genotype calling criteria. Multiple records for the same individual indicate different versions of the data (shotgun sequences as opposed to capture) or different ways of calling genotypes (consensus genotype calling or genotype calling using snpAD). Reported are numbers for all sites and for sites excluding C-T and G-A polymorphisms. The proportions are calculated relative to the total number of available sites (6,912,728; SI 1).

| name | all sites |  | excluding C-to-T/G-to-A |  |
| --- | --- | --- | --- | --- |
|  | count | proportion | count | proportion |
| Spy 94a (shotgun) | 2,664,787 | 38.5% | 2,636,405 | 38.1% |
| Spy 94a | 3,502,380 | 50.7% | 3,470,671 | 50.2% |
| Spy 94a (snpAD) | 3,557,124 | 51.5% | 3,522,759 | 51.0% |
| Denisova 4 | 4,731,302 | 68.4% | 4,705,860 | 68.1% |
| Denisova 4 (snpAD) | 4,824,242 | 69.8% | 4,798,037 | 69.4% |
| Denisova 8 | 5,851,332 | 84.6% | 5,828,356 | 84.3% |
| Denisova 8 (snpAD) | 6,260,242 | 90.6% | 6,236,272 | 90.2% |
| Mezmaiskaya 2 (shotgun) | 3,822,106 | 55.3% | 3,762,768 | 54.4% |
| Mezmaiskaya 2 | 6,348,948 | 91.8% | 6,346,684 | 91.8% |
| Mezmaiskaya 2 (snpAD) | 6,669,912 | 96.5% | 6,667,441 | 96.5% |

**Table S5.2.** Counts of sites for each modern human Y chromosome (consensus genotype calls of shotgun data) in 6.9 Mb capture regions which passed the filtering for minimum depth of 3 reads in addition to other filtering and genotype calling criteria. Reported are numbers for all sites and for sites excluding C-T and G-A polymorphisms. The proportions are calculated relative to the total number of available sites (6,912,728).

| name | all sites |  | excluding C-to-T/G-to-A |  |
| --- | --- | --- | --- | --- |
|  | count | proportion | count | proportion |
| A00-1 | 6,852,077 | 99.12% | 6,851,175 | 99.11% |
| A00-2 | 6,867,616 | 99.35% | 6,866,716 | 99.33% |
| A00 | 6,873,078 | 99.43% | 6,872,184 | 99.41% |
| Ust'-Ishim | 6,888,071 | 99.64% | 6,887,986 | 99.64% |
| S_Yoruba-2 | 6,897,101 | 99.77% | 6,896,820 | 99.77% |
| S_Mandenka-1 | 6,898,162 | 99.79% | 6,897,879 | 99.79% |
| S_Mbuti-1 | 6,898,694 | 99.80% | 6,898,400 | 99.79% |
| S_Gambian-1 | 6,899,210 | 99.80% | 6,898,929 | 99.80% |
| S_Papuan-2 | 6,904,393 | 99.88% | 6,904,208 | 99.88% |
| S_Thai-1 | 6,905,023 | 99.89% | 6,904,846 | 99.89% |
| S_BedouinB-1 | 6,905,031 | 99.89% | 6,904,820 | 99.89% |
| S_Saami-2 | 6,905,291 | 99.89% | 6,905,102 | 99.89% |
| S_Sardinian-1 | 6,905,558 | 99.90% | 6,905,352 | 99.89% |
| S_Ju_hoan_North-1 | 6,906,228 | 99.91% | 6,905,872 | 99.90% |
| S_Karitiana-1 | 6,906,409 | 99.91% | 6,906,258 | 99.91% |
| S_Dai-2 | 6,906,414 | 99.91% | 6,906,248 | 99.91% |
| S_Burmese-1 | 6,906,438 | 99.91% | 6,906,260 | 99.91% |
| S_Dinka-1 | 6,906,507 | 99.91% | 6,906,225 | 99.91% |
| S_French-1 | 6,906,914 | 99.92% | 6,906,738 | 99.91% |
| S_Punjabi-1 | 6,907,065 | 99.92% | 6,906,885 | 99.92% |
| S_Finnish-2 | 6,907,088 | 99.92% | 6,906,889 | 99.92% |
| S_Han-2 | 6,907,603 | 99.93% | 6,907,417 | 99.92% |
| S_Turkish-1 | 6,907,751 | 99.93% | 6,907,717 | 99.93% |

**Table S5.3.**
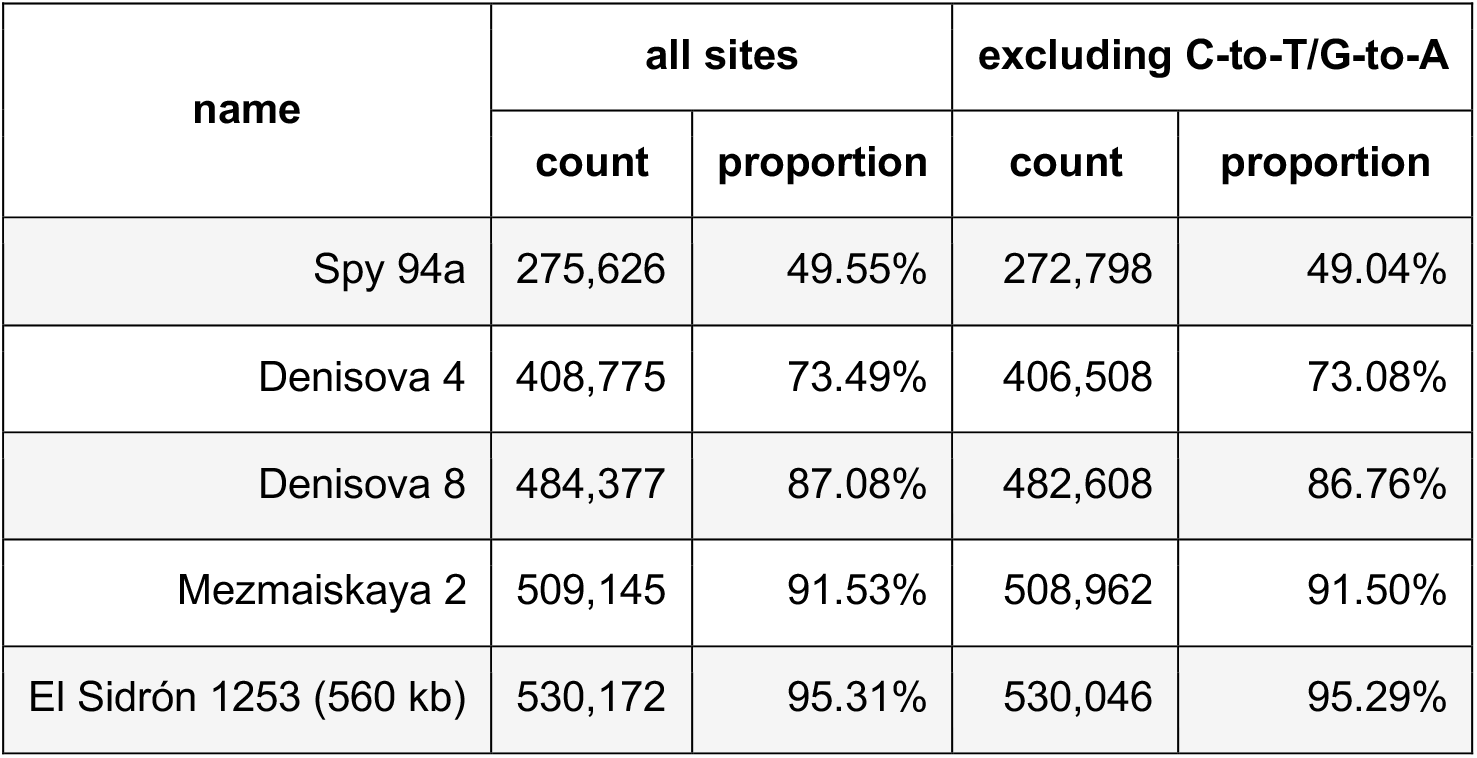
Counts of sites for each archaic human Y chromosome (in 560 kb capture regions) which passed the filtering for minimum depth of 3 reads in addition to other filtering and genotype calling criteria. Reported are numbers for all sites and for sites excluding C-to-T and G-to-A polymorphisms. The proportions are calculated relative to the total number of available sites (556,259).

| name | all sites |  | excluding C-to-T/G-to-A |  |
| --- | --- | --- | --- | --- |
|  | count | proportion | count | proportion |
| Spy 94a | 275,626 | 49.55% | 272,798 | 49.04% |
| Denisova 4 | 408,775 | 73.49% | 406,508 | 73.08% |
| Denisova 8 | 484,377 | 87.08% | 482,608 | 86.76% |
| Mezmaiskaya 2 | 509,145 | 91.53% | 508,962 | 91.50% |
| El Sidrón 1253 (560 kb) | 530,172 | 95.31% | 530,046 | 95.29% |

**Table S5.4.**
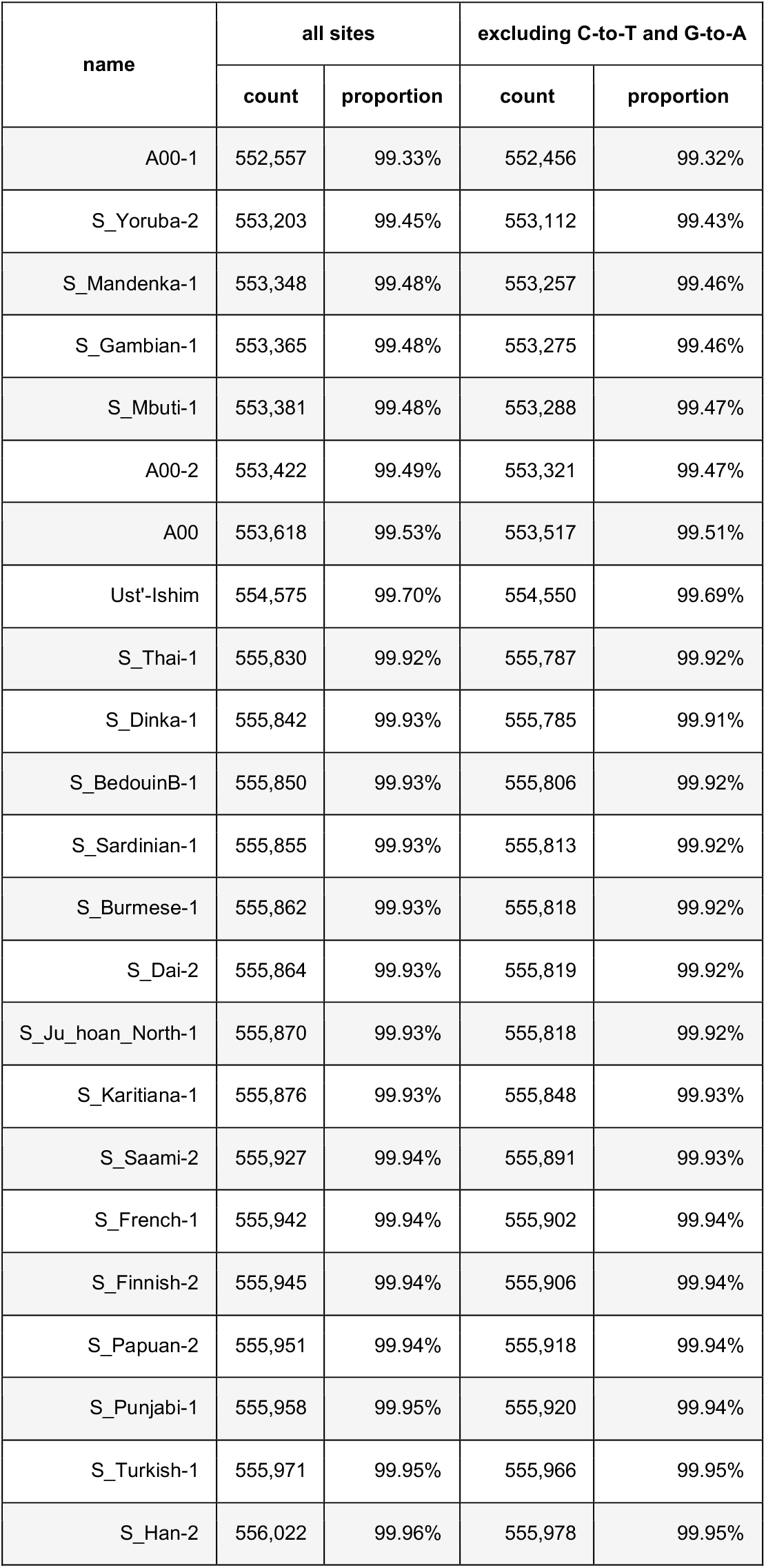
Counts of sites for each modern human Y chromosome (in 560 kb capture regions) which passed the filtering for minimum depth of 3 reads in addition to other filtering and genotype calling criteria. Reported are numbers for all sites and for sites excluding C-to-T and G-to-A polymorphisms. The proportions are calculated relative to the total number of available sites (556,259).

| name | all sites |  | excluding C-to-T and G-to-A |  |
| --- | --- | --- | --- | --- |
|  | count | proportion | count | proportion |
| A00-1 | 552,557 | 99.33% | 552,456 | 99.32% |
| S_Yoruba-2 | 553,203 | 99.45% | 553,112 | 99.43% |
| S_Mandenka-1 | 553,348 | 99.48% | 553,257 | 99.46% |
| S_Gambian-1 | 553,365 | 99.48% | 553,275 | 99.46% |
| S_Mbuti-1 | 553,381 | 99.48% | 553,288 | 99.47% |
| A00-2 | 553,422 | 99.49% | 553,321 | 99.47% |
| A00 | 553,618 | 99.53% | 553,517 | 99.51% |
| Ust'-Ishim | 554,575 | 99.70% | 554,550 | 99.69% |
| S_Thai-1 | 555,830 | 99.92% | 555,787 | 99.92% |
| S_Dinka-1 | 555,842 | 99.93% | 555,785 | 99.91% |
| S_BedouinB-1 | 555,850 | 99.93% | 555,806 | 99.92% |
| S_Sardinian-1 | 555,855 | 99.93% | 555,813 | 99.92% |
| S_Burmese-1 | 555,862 | 99.93% | 555,818 | 99.92% |
| S_Dai-2 | 555,864 | 99.93% | 555,819 | 99.92% |
| S_Ju_hoan_North-1 | 555,870 | 99.93% | 555,818 | 99.92% |
| S_Karitiana-1 | 555,876 | 99.93% | 555,848 | 99.93% |
| S_Saami-2 | 555,927 | 99.94% | 555,891 | 99.93% |
| S_French-1 | 555,942 | 99.94% | 555,902 | 99.94% |
| S_Finnish-2 | 555,945 | 99.94% | 555,906 | 99.94% |
| S_Papuan-2 | 555,951 | 99.94% | 555,918 | 99.94% |
| S_Punjabi-1 | 555,958 | 99.95% | 555,920 | 99.94% |
| S_Turkish-1 | 555,971 | 99.95% | 555,966 | 99.95% |
| S_Han-2 | 556,022 | 99.96% | 555,978 | 99.95% |

### 6. Inferring phylogenetic relationships

To resolve the phylogenetic relationships of each archaic human Y chromosome to all other Y chromosome sequences, we merged the VCF files with genotype calls from each individual (including the chimpanzee) into a single VCF file and converted the genotypes to the FASTA format using a custom Python script (available on our Github repository: https://github.com/bodkan/archaic-ychr). To mitigate biases introduced by low coverage and characteristics of aDNA damage (*29*), we excluded all C-to-T and G-to-A polymorphisms and applied the same filters for each individual as for all other analyses in our study. Finally, we excluded monomorphic sites and sites carrying private changes on the chimpanzee lineage to reduce the size of the final alignment file.

To construct a neighbor-joining phylogenetic tree (Figure 2A), we utilized the functionality provided by R packages *ape* and *phangorn* (*30, 31*). First, we calculated the distance matrix between all Y chromosome pairs in the FASTA file with the function `dist.dna`, using the model of simple pairwise differences (`model = “raw”`) and excluding sites with missing data specific to each pair (`pairwise.deletion = TRUE`). We then provided this distance matrix to the `nj` function and rooted the resulting neighbor-joining tree using the function `midpoint` from the *phangorn* package. Bootstrap confidence numbers for the neighbor-joining tree were calculated using *ape*’s `boot.phylo` function over 100 replicates. After inspecting the resulting phylogenetic tree, we found that the private branch leading to the *Denisova 4* had a negative length (value = −0.00088). Given that negative branch lengths are a relatively common artefact of the neighbor-joining algorithm and do not affect the reliability of the generated tree we followed the recommendation to set the branch length to zero (*32*). We note that this does not have any impact on our conclusions, because the change involves a private branch whose length is not meaningful given the discrepancies between sample dates and implied tree tip dates (Figures 1A and 2A, (*29*)). The final trees were annotated and plotted using the R package *ggtree* (*33*).

### 7. Estimating the TMRCA of archaic and modern human Y chromosomes

Given that most of the Y chromosome capture data analyzed in our study is of relatively low coverage (Figure 1B, Table S4.1), care needs to be taken when estimating phylogenetic parameters such as the time to the most recent common ancestor (TMRCA). Similarly, low coverage and the associated reduction in the accuracy of genotype calls render the inferred aDNA branch lengths unreliable (*29*). Any phylogenetic method of choice must be therefore robust to sequencing errors and incorrect branch lengths. We also observe discordances between sample dates and implied molecular tip dates, likely due to residual genotype calling errors (compare Figure 1A vs Figure 2A). We therefore estimated TMRCAs between archaic and modern human Y chromosomes using a method inspired by the analysis of the *El Sidrón 1253* Neandertal coding sequence (*5*). Instead of using polymorphisms on the archaic human lineage, this method relies on first estimating the TMRCA of a pair of high-coverage African and non-African Y chromosomes (*TMRCA_AFR_*) which is then used to extrapolate the deeper divergence time between archaic and modern human Y chromosomes. We describe the method in the sections below, detailing our modifications and improvements.

#### 7.1. TMRCA of Africans and non-Africans *TMRCA_AFR_*

The original study by Mendez *et al*. estimated the TMRCA between the A00 African Y chromosome lineage and the hg19 Y chromosome as a representative of non-African Y chromosomes (*5, 21*). In order to get a better sense of the uncertainty and noise in our TMRCA estimates, we expanded the present-day Y chromosome reference panel to 13 non-African and 6 African Y chromosomes from the SGDP data set (Table S7.1) (*20*).

In the first step, we estimated mutation rate in the 6.9 Mb capture target using the high-coverage Y chromosome of *Ust’-Ishim*, a 45,000 years old hunter-gatherer from Siberia (*22*). We counted derived mutations missing on the *Ust’-Ishim* branch compared to those observed in the panel of 13 non-African Y chromosomes, and used this branch-shortening to calculate mutation rate assuming generation time of 25 years (Figure S7.1, Table S7.2). In the second step, we counted mutations accumulated on an African lineage and a non-African lineage since their split from each other and calculated the TMRCA of both (in units of years ago) using the mutation rate estimated in the first step. Importantly, we discovered that the branch-lengths in Africans are as much as 13% shorter compared to non-Africans (Figure S7.3), which is consistent with significant branch length variability discovered in previous studies and suggested to be a result of various demographic and selection processes (*35, 36*). To keep our methodology consistent throughout our analyses, we estimated the TMRCA of African and non-African Y chromosomes as the length of the non-African Y lineage (sum of branch lengths *a* + *d* in Figure S7.1). Encouragingly, we found that our mutation rate and TMRCA*_AFR_* point estimates (Table S7.2) match closely those based on a large panel of present-day Y chromosomes (*21*). Most importantly, using the A00 lineage as a representative of the deepest known split among present-day human Y chromosomes we inferred a TMRCA*_A00_* of ∼249 years ago (point estimate based on an estimated mutation rate of 7.34×10^-10^ per bp per year), which is comparable to the TMRCA*_A00_* of ∼254 years ago estimated by Karmin et al., 2015. Therefore, our more restricted 6.9 Mb capture target gives TMRCA estimates consistent with those obtained from the full Y chromosome shotgun data (*21*).

**Figure S7.1.**
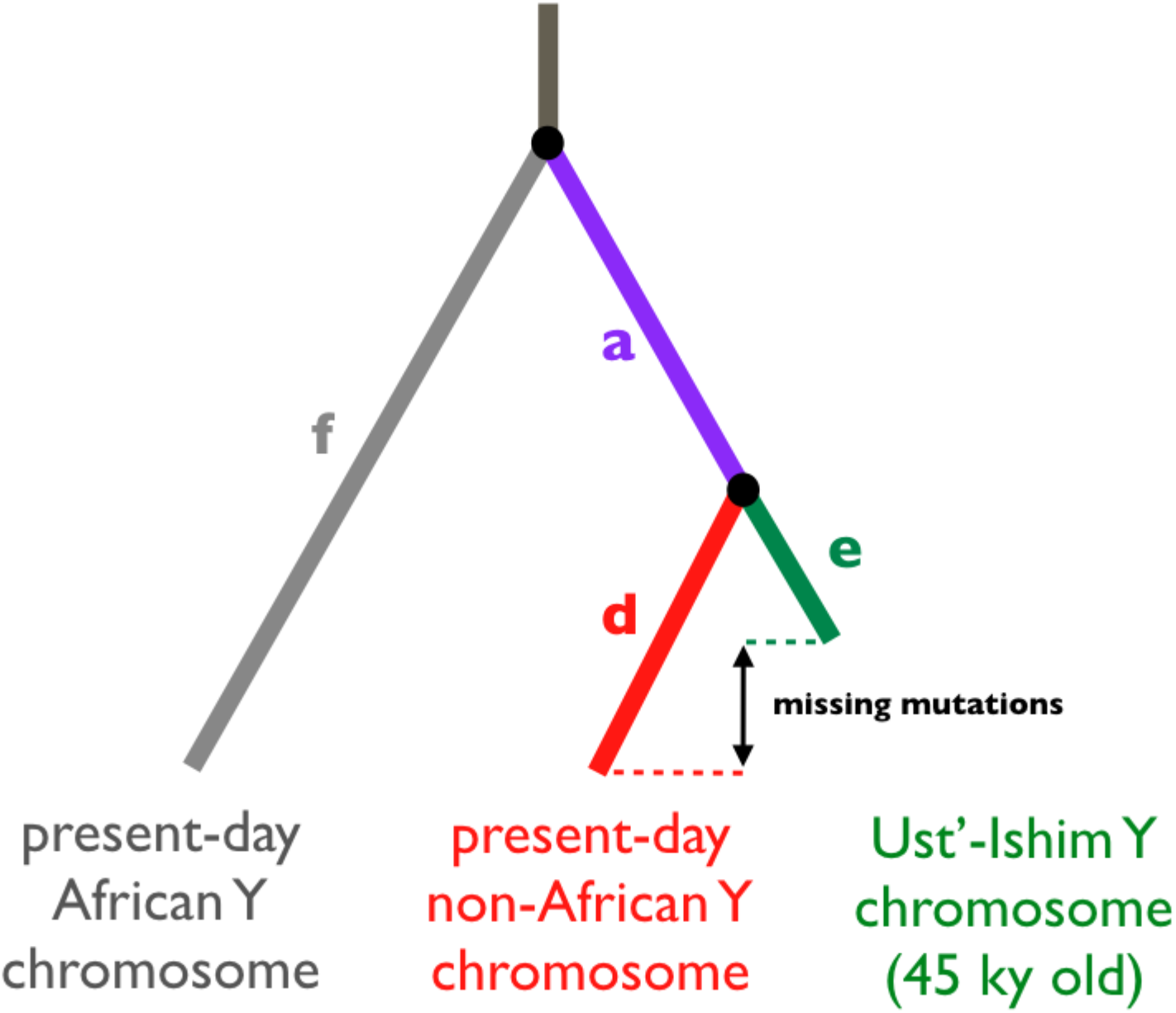
Schematic of the branch-counting method to estimate the mutation rate and split times of African and non-African Y chromosomes. Accurate knowledge of the age of the *Ust’-Ishim* individual (*22*) makes it possible to estimate the mutation rate within the 6.9 Mb target capture regions. We use the number of mutations missing on the *Ust’-Ishim* lineage since this individual died (45 kya) and compare it to another non-African Y chromosome, i.e. the quantity *d - e*. We used this mutation rate to calculate the TMRCA between a pair of non-African and African Y chromosomes as the total length of the branches *a + d*.

**Figure S7.2.**
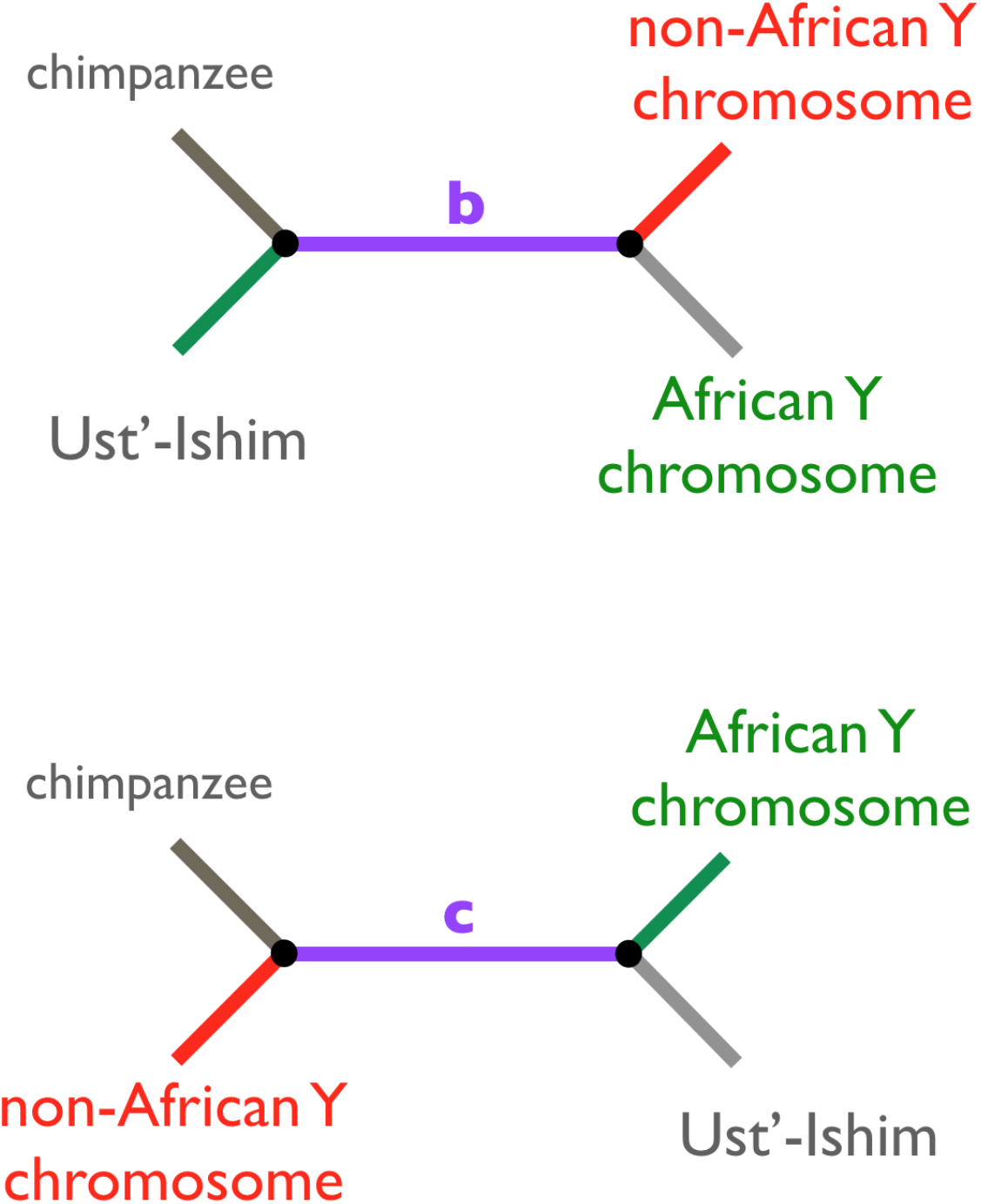
Two alternative site patterns which are discordant with the true phylogenetic relationship. Given that the true phylogenetic relationship is that shown in Figure S7.1, these alternative branch patterns must be a result of double mutations or genotype calling errors.

**Figure S7.3.**
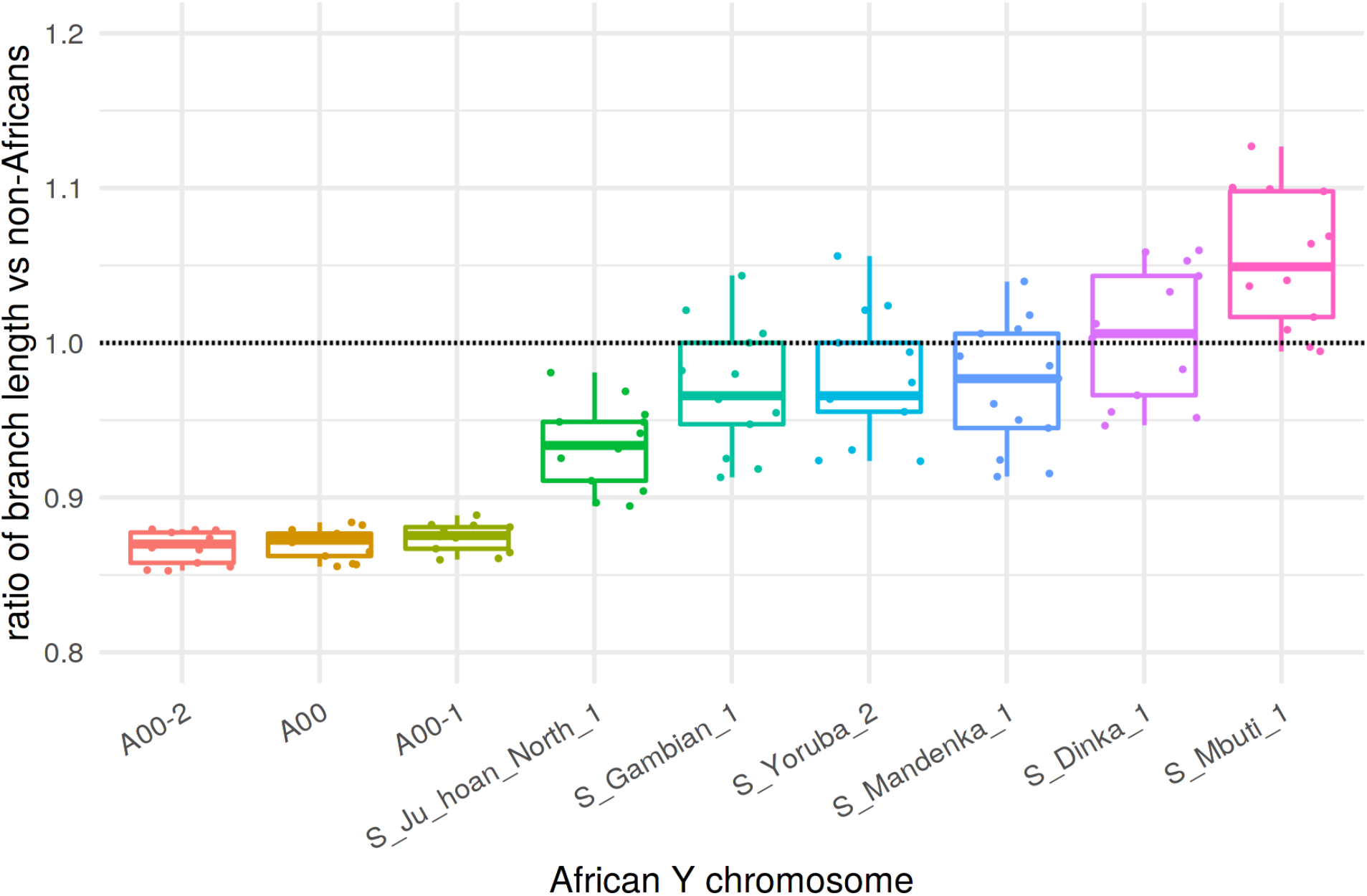
Branch length differences between African Y chromosomes and a panel of 13 non-African Y chromosomes. Ratios were calculated by creating an alignment of chimpanzee, African and non-African Y chromosomes and taking the ratio of the number of derived alleles observed in an African (x-axis) and the number of derived alleles in each of the individual non-Africans (dots, Table S7.1). “A00” represents a merge of sequences of two lower coverage Y chromosomes, A00-1 and A00-2 (Table S4.3).

**Table S7.1.** Haplogroups of present-day human Y chromosomes used in our reference panel. Haplogroup names were taken from the SGDP annotation table (*20*). Haplogroups of the African panel are highlighted in gray. The A00 individual represents a merge of lower coverage sequences of two individuals, here named A00-1 and A00-2 (Table S4.3).

| individual | haplogroup |
| --- | --- |
| S_Finnish-2 | I1a1b5 |
| S_Sardinian-1 | J1b2b |
| S_BedouinB-1 | J1b2b |
| S_Punjabi-1 | J2a1 |
| S_French-1 | J2a1b1 |
| S_Turkish-1 | R1b1a2a1a2c1g |
| S_Karitiana-1 | Q1a2a1a1 |
| S_Saami-2 | N1c1a1a2a |
| S_Burmese-1 | O3a1c2 |
| S_Thai-1 | O3a2c |
| S_Dai-2 | O2a1 |
| S_Han-2 | O1a1 |
| S_Papuan-2 | S |
| S_Dinka-1 | E2a |
| S_Gambian-1 | E1b1a1a1f |
| S_Ju_hoan_North-1 | B2b1b |
| S_Mandenka-1 | E1b1a1a1g1 |
| S_Mbuti-1 | E1b1a1a1g1 |
| S_Yoruba-2 | E1b1a1a1f1b1 |
| A00 | A00 |
| A00-1 | A00 |
| A00-2 | A00 |

**Table S7.2.** Branch counts and estimates of mutation rate and TMRCA between an African lineage and a panel of 13 non-African Y chromosomes. All quantities represent averages across all non-African Y chromosomes (Table S7.1). Counts in columns *a* to *f* represent counts of site patterns as shown in Figures S7.1 and S7.2. *“Total”* represents the number of sites out of the total 6.9 Mb of target sequence available for the analysis. The last two columns represent the inferred mutation rate based on the *Ust’-Ishim* branch-shortening and the average TMRCA between a given African and a panel of non-Africans calculated from the length of the 7 + 9 branch as shown in Figure S7.1.

| African | a | b | c | d | e | f | total | mutation rate<br>(bp<sup>-1</sup> × year<sup>-1</sup>) | TMRCA<sub>AFR</sub><br>(years ago) |
| --- | --- | --- | --- | --- | --- | --- | --- | --- | --- |
| A00 | 1060.2 | 1.2 | 8.3 | 214.5 | 14.2 | 1103.1 | 6,064,477 | 7.34e-10 | 249211.63 |
| A00-2 | 1054.5 | 1.2 | 8.2 | 214.7 | 14.6 | 1092.7 | 6,045,632 | 7.35e-10 | 247074.54 |
| A00-1 | 1004.2 | 1.2 | 8.1 | 207.2 | 14.2 | 1054.1 | 5,855,414 | 7.32e-10 | 246756.41 |
| S_Ju_hoan_North_1 | 261.5 | 2.7 | 8.5 | 216.3 | 14.5 | 439.2 | 6,100,839 | 7.35e-10 | 98443.62 |
| S_Mbuti_1 | 116.5 | 1.2 | 8.9 | 217.5 | 14.5 | 344.7 | 6,123,504 | 7.37e-10 | 76782.21 |
| S_Dinka_1 | 110.8 | 2.1 | 8.7 | 217.4 | 14.5 | 321.2 | 6,090,467 | 7.40e-10 | 71573.49 |
| S_Mandenka_1 | 120.9 | 1.2 | 8.8 | 218.3 | 14.5 | 321.0 | 6,123,094 | 7.39e-10 | 71226.46 |
| S_Gambian_1 | 118.8 | 1.2 | 8.5 | 218.1 | 14.5 | 320.2 | 6,096,582 | 7.42e-10 | 71127.40 |
| S_Yoruba_2 | 118.9 | 1.2 | 8.7 | 217.6 | 13.5 | 320.9 | 6,121,558 | 7.41e-10 | 71119.46 |

### 7.2. Archaic human-modern human TMRCA (*TMRCA _archaic_*)

Having estimated *TMRCA_AFR_* (section 7.1), we can express the TMRCA of archaic and modern human Y chromosomes (*TMRCA_archanic_*) as a factor of how much older is *TMRCA_archanic_* compared to *TMRCA_AFR_* (Figure S7.4). In mathematical terms, if we call the scaling factor α (following the terminology of (*5*)), we can write

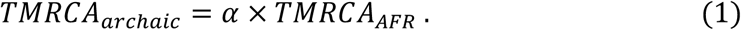

In the remainder of this section, we present two ways of calculating G, first using the original approach of Mendez *et al*. and then using a more straightforward method.

#### Mendez et al. approach to calculate *α*

Based on Figure S7.4, an alternative way to express *TMRCA_archanic_* in addition to equation (1) is

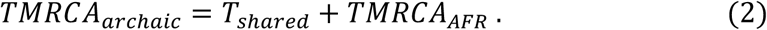

The expressions (1) and (2) define a system of two equations and three variables, which can be solved for *T_shared_* to get

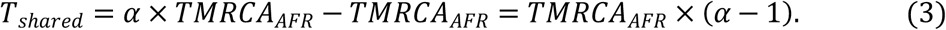

Mendez *et al*. found an expression for *α* by considering a ratio of time shared by hg19 and A00 Y chromosomes after their split from the *El Sidrón 1253* Neandertal (*T_shared_* above) and private branch lengths of both (Figure S7.4), arriving at the following expression:

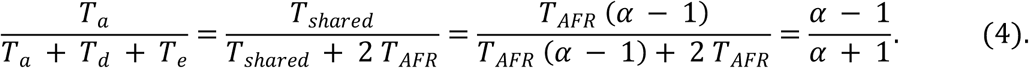

Assuming mutation rate constancy on different lineages, *α* can be found by solving the following equation

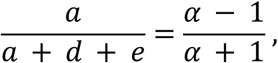

which leads to

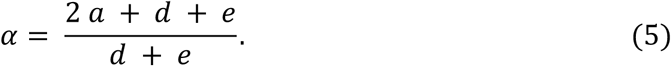

Using this expression for α and the values of *TMRCA_AFR_* estimated in section 7.1, we can calculate *TMRCA_archanic_* for each pair of archaic and non-African Y chromosomes using equation (5) (Figure 7.9B).

#### A more robust α statistic

While investigating the effect of minimum coverage filtering on genotype calling accuracy (section 5.3), we discovered a concerning dependence of the apparent branch lengths on the choice of the minimum coverage cutoff. Under normal conditions, the relative proportions of branch lengths *a*, *d* and *e* (Figure S7.4) should remain constant regardless of coverage. This is crucial because the G estimator proposed by Mendez *et al.* is expressed in terms of proportions of lengths of all three of these branches (equation (5)).

Strikingly, we found that although the proportions of 7 and 9 branch lengths remain relatively stable even for extremely strict coverage filters, the relative length of the O branch (given by the proportion of derived mutations on the private African branch) has increasing tendency (Figures S7.6 and S7.7). Furthermore, although this effect is most pronounced in low coverage samples (Figures S7.6 and S7.7), it is clearly present even in the high coverage *Mezmaiskaya 2*, although at much higher coverage cutoffs (Figure S7.8). Therefore, the issue is clearly not sample-specific but is a common artifact caused by pushing the minimum required coverage close to, or even beyond, the average coverage. Restricting to sites with high number of aligned reads leads to enrichment of regions of lower divergence from the reference sequence, distorting the normal proportions of derived mutations observed on different branches of the tree.

We note that there is a more straightforward way to express the scaling factor α:

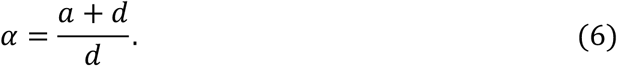

This follows trivially from the definition of α as the factor of how much deeper *TMRCA_archanic_* is compared to *TMRCA_AFR_* and, unlike the original formulation of α (equation (5)), has the advantage of not relying on the relative length of the African branch *e*. This is important not only because of discordant branch proportion patterns (Figures S7.6-S7.8) but also due to known unequal branch lengths observed in African and non-African Y chromosome lineages (Figure S7.3, (*36, 37*)).

For completeness, we note that in a situation without any bias we can assume *e* ≈ *d*. Substituting for *e* in equation (5) then gives

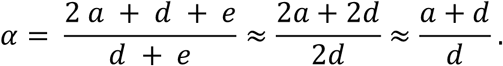

Therefore, under ideal conditions, both approaches to calculate α (equations (5) and (6)) are mathematically equivalent.

Using the new expression for G and the values of *TMRCA_AFR_* estimated in section 7.1, we can estimate *TMRCA_archanic_* for each pair of archaic and non-African Y chromosomes using equation (5) (Figure 2A, Table S7.3). By comparing TMRCA results based on the two formulations for α, we found similar estimates for most of the archaic human Y chromosomes in our study (Figures S7.9 and S7.10). The only exception are the two Denisovan Y chromosomes, for which we infer slightly higher TMRCA with modern humans using the new α estimation procedure compared to the formulation based on the original method (Figures S7.9 and S7.10). This is a consequence of an increased proportion of the *e* branch relative to the *d* branch in *Denisova 4* and *Denisova 8* at the chosen minimum coverage filter which is evident in Figure S7.7. Because the original method of Mendez *et al.* relies on the *e* count of the derived African alleles (equation (5)), this leads to a slight decrease in the value of the α factor and, consequently, to a lower inferred TMRCA value. In contrast, our new formulation of α (equation (6)) is robust to this artifact and the inferred values of TMRCA are not affected.

Finally, we want to emphasize that although the analyses of branch length discrepancies discussed in this section were mostly based on results obtained using the A00 Y chromosome lineage, the issues we discovered are not specific to a particular choice of an African Y chromosome (Figure S7.8A-C). However, comparisons of TMRCA estimates for the low coverage samples with those obtained for the high coverage samples (which do not show any biases at coverage cutoffs used throughout our study) clearly show that the inferences are most stable when the A00 lineage is used in the calculation of the α scaling factor, even for the samples with lowest coverage (Figure S7.13). Therefore, all main results in our study are based on calculations using the A00 high coverage Y chromosome.

**Figure S7.4.**
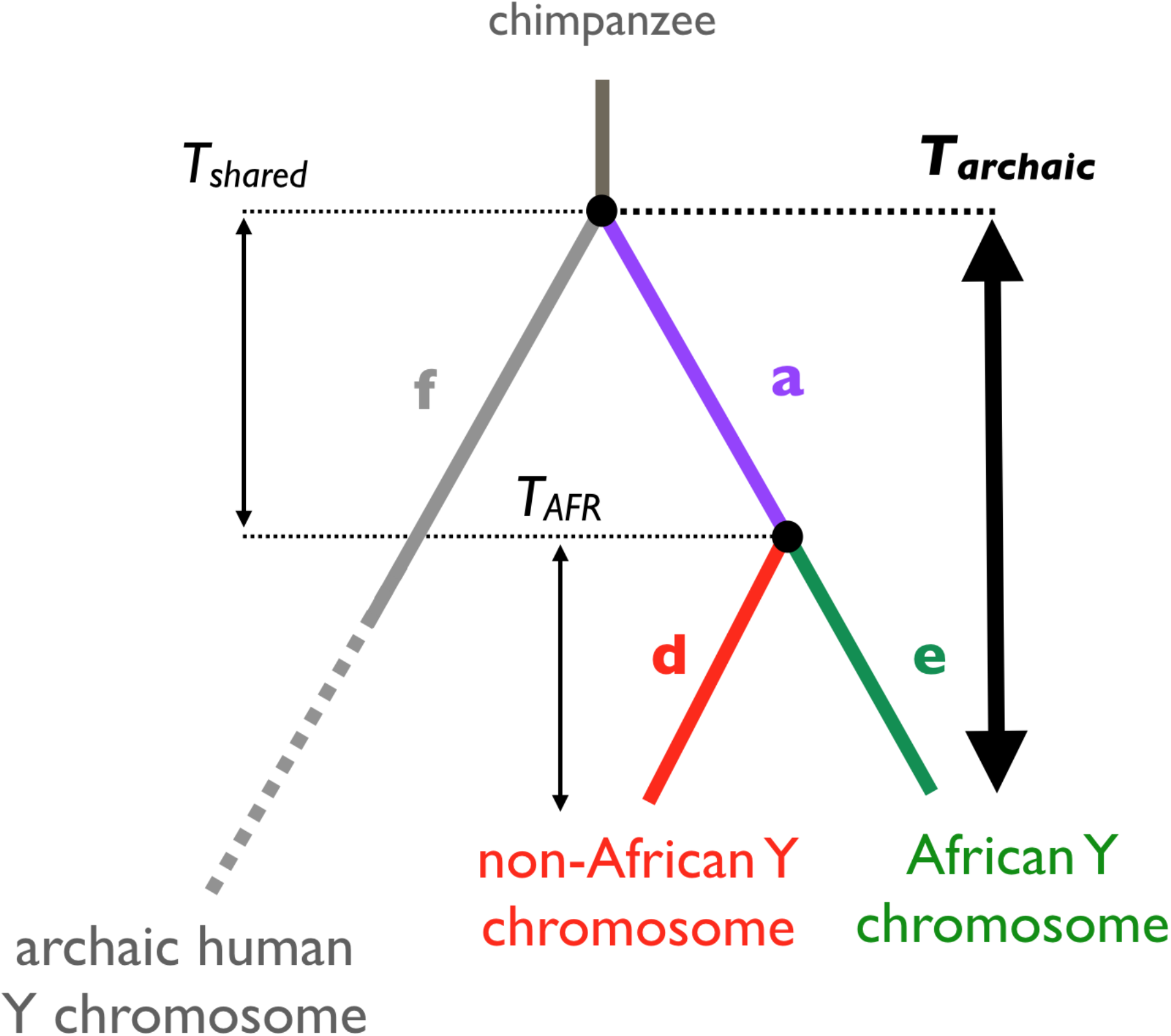
Branch-counting method to estimate the TMRCA of archaic and present-day human Y chromosomes. As explained in section 7.2, we can decompose the quantity of interest (*T_ARCH_*, thick arrow) using two quantities *T_shared_* and *T_AFR_* and express it simply as a factor of *T_AFR_*, which can be accurately estimated using high-quality modern human Y chromosome sequences (section 7.1).

**Figure S7.5.**
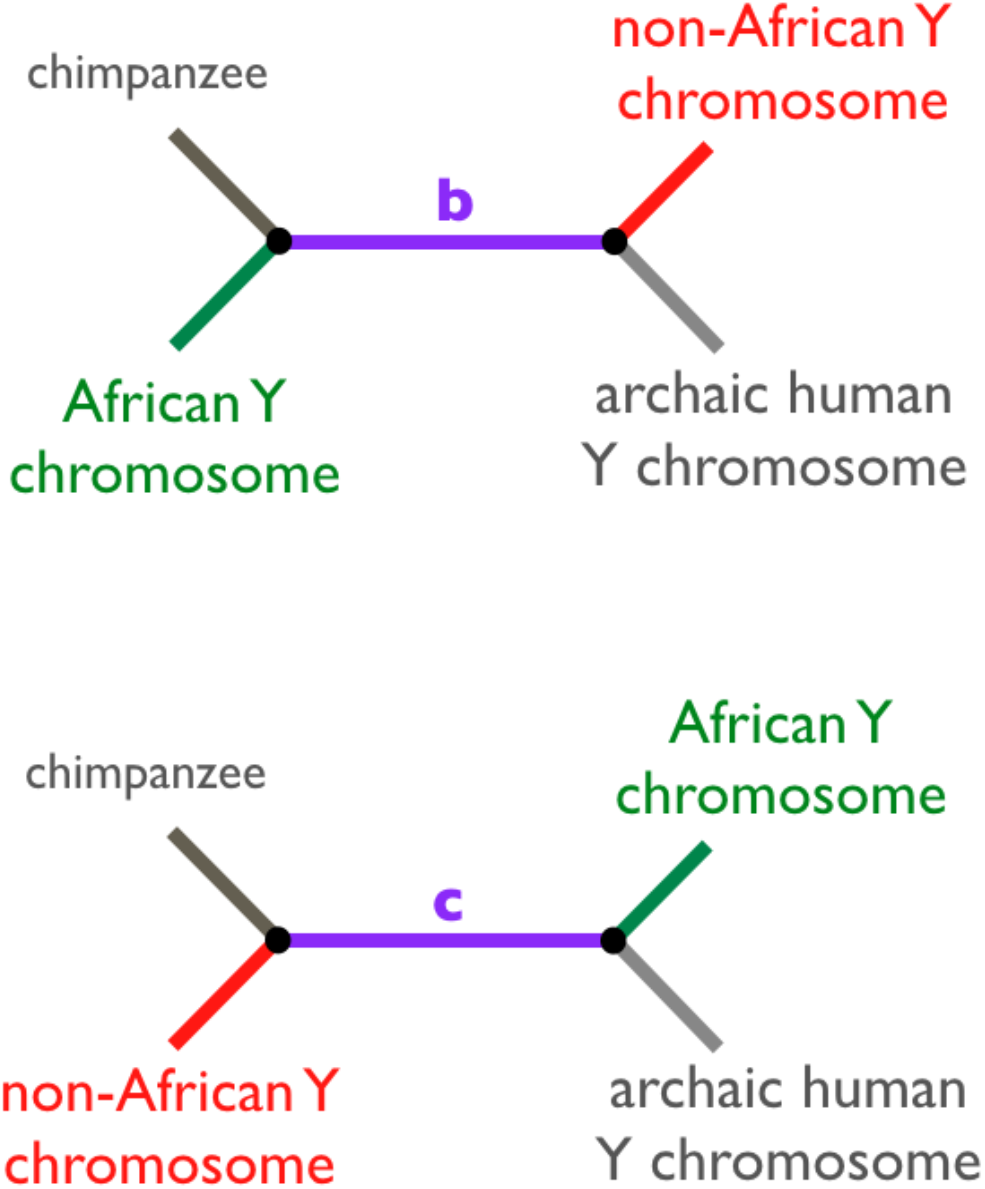
Alternative tree topologies with two additional possible branch counts *b* and *c*. These topologies are incongruent with the true phylogenetic trees and the branch counts *b* and *c* are most likely a result of back mutations or sequencing errors.

**Figure S7.6.**
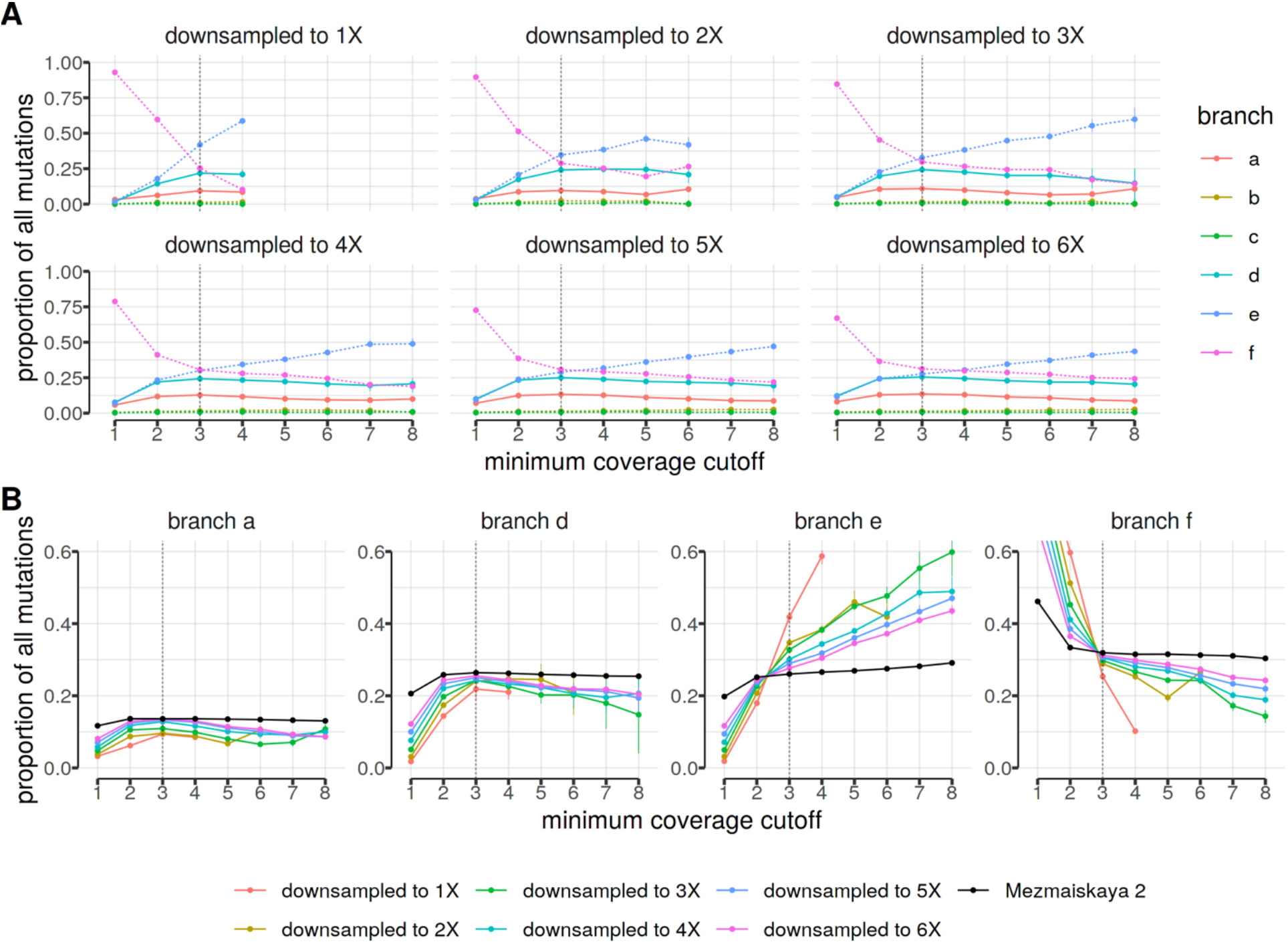
Relative proportion of branch lengths in downsampled *Mezmaiskaya 2* data as a function of minimum coverage cutoff. **(A)** Panels show results for 14.3X *Mezmaiskaya 2* downsampled down to 1X, 2X, … 6X coverage. **(B)** Same as in panel (A) but partitioned per branch. Black solid lines show expectations based on the full *Mezmaiskaya 2* data. Increased relative proportions of the *f* branch lengths are due to false polymorphisms at low coverage cutoffs. Branch length proportions (labeled as in Figure S7.4) were calculated as 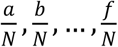, where *N* = *a* + *b* + ⋯ + *f*. Vertical dotted lines indicate a 3X lower coverage cutoff used throughout our study (section 5.3). Branch counts were averaged over pairs of 13 non-Africans and the A00 African Y chromosome. Analysis is based on all classes of polymorphisms.

**Figure S7.7.**
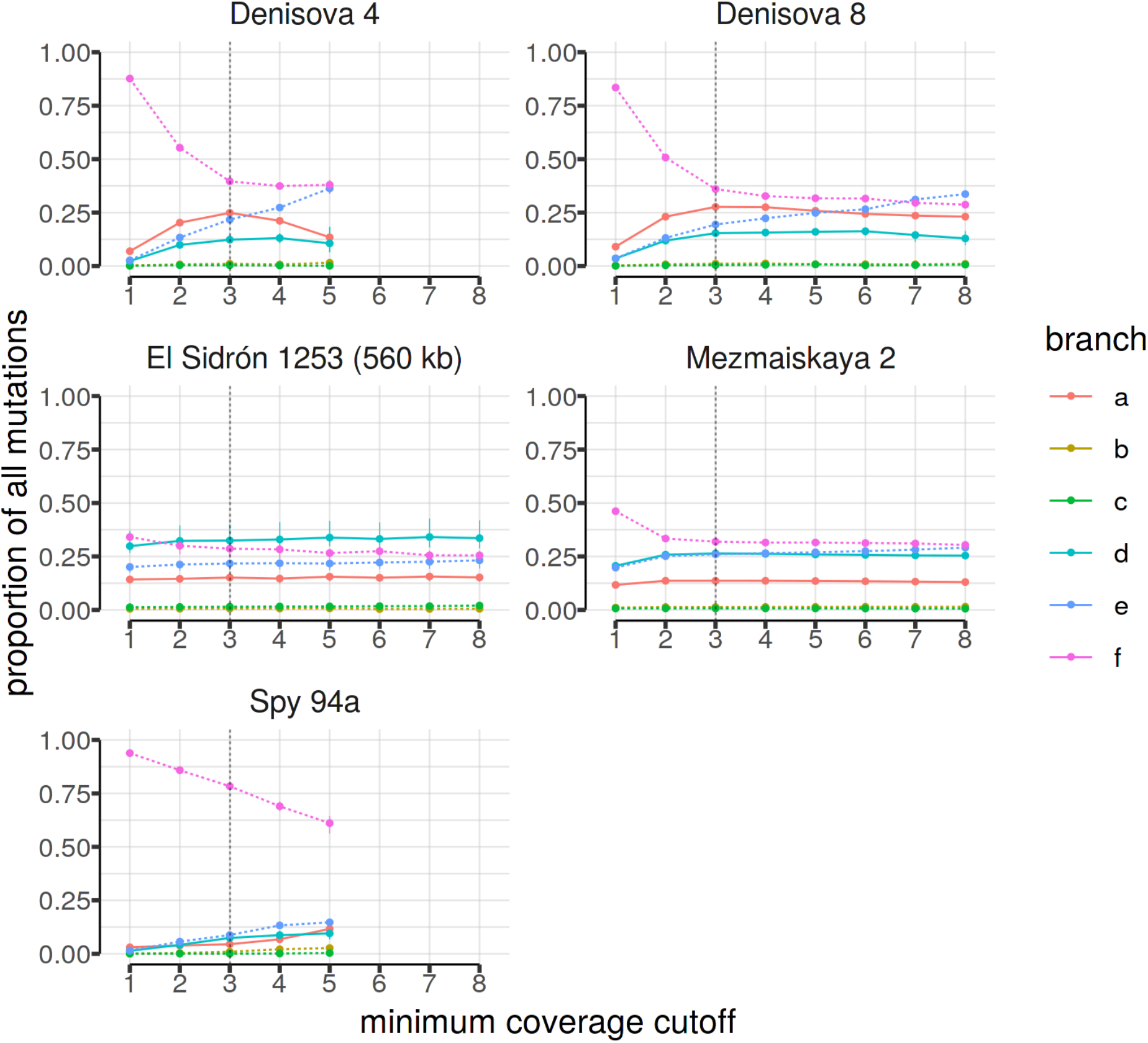
Relative proportion of branch lengths as a function of minimum coverage support required for each genotype call. Branch length proportions (colored lines) were calculated as 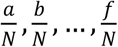, where *N* = *a* + *b* + ⋯ + *f*. Vertical dotted lines indicate a 3X lower coverage cutoff used throughout our study (section 5.3). With the exception of *El Sidrón 1253,* all panels show results for the 6.9 Mb capture data. Analysis is based on all polymorphisms.

**Figure S7.8.**
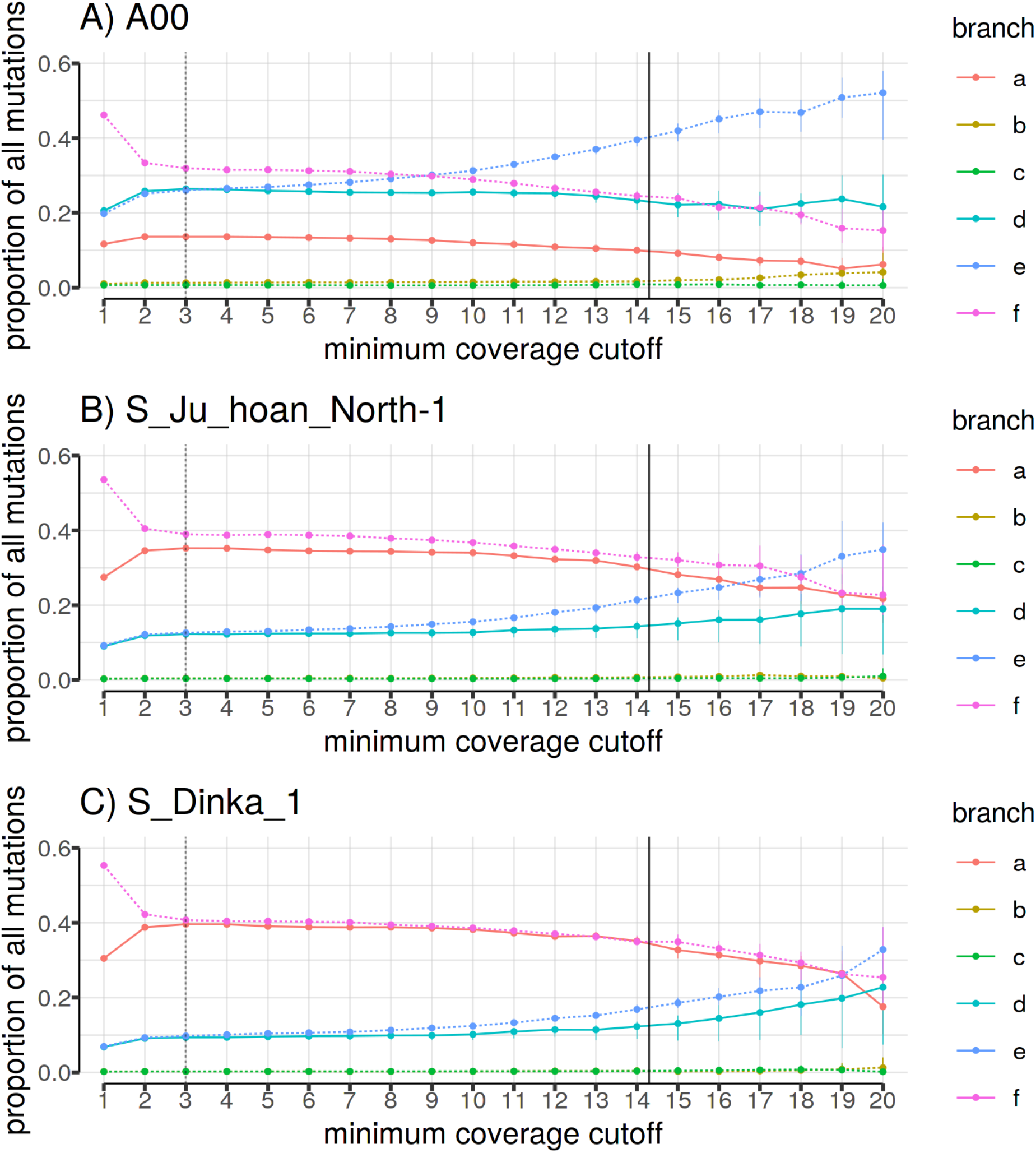
Relative proportions of branch lengths in the 14.3X *Mezmaiskaya 2* capture data as a function of minimum coverage support required for each genotype call. *Mezmaiskaya 2* was chosen for this example because its high coverage makes the branch proportion patterns stand out more clearly. Panels (A), (B) and (C) show results based on three different African Y chromosomes used to define branch *e* (Figure S7.4). Branch length proportions (colored lines) were calculated as 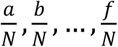, where *N* = *a* + *b* + ⋯ + *f*, using the complete 6.9 Mb capture data of *Mezmaiskaya 2*. Vertical dotted lines indicate a 3X lower coverage cutoff used throughout our study (section 5.3). Analysis is based on all polymorphisms. Vertical solid lines indicate the average coverage in the *Mezmaiskaya 2* individual (14.3X).

**Figure S7.9.**
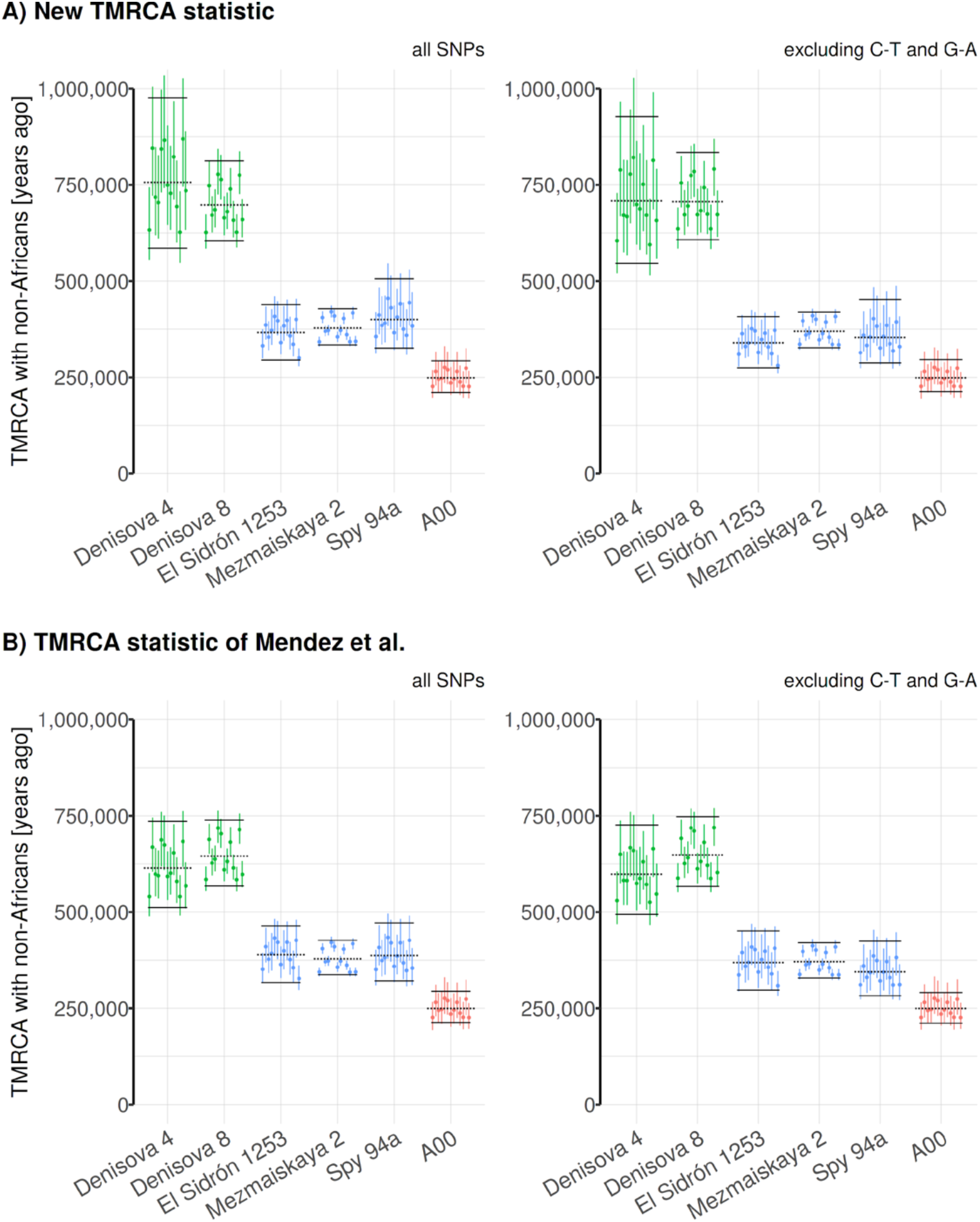
Comparison of TMRCA estimates obtained using the new statistic and the original approach used in the analysis of the *El Sidrón 1253* Neandertal. **(A)** Estimates using our new, more robust TMRCA estimate. **(B)** Original calculation proposed by *Mendez et al.*, 2016 (*5*). Shown are estimates based on all polymorphisms (left panels) and excluding C-to-T or G-to-A polymorphisms which are more likely to be caused by aDNA damage (right panels).

**Figure S7.10.**
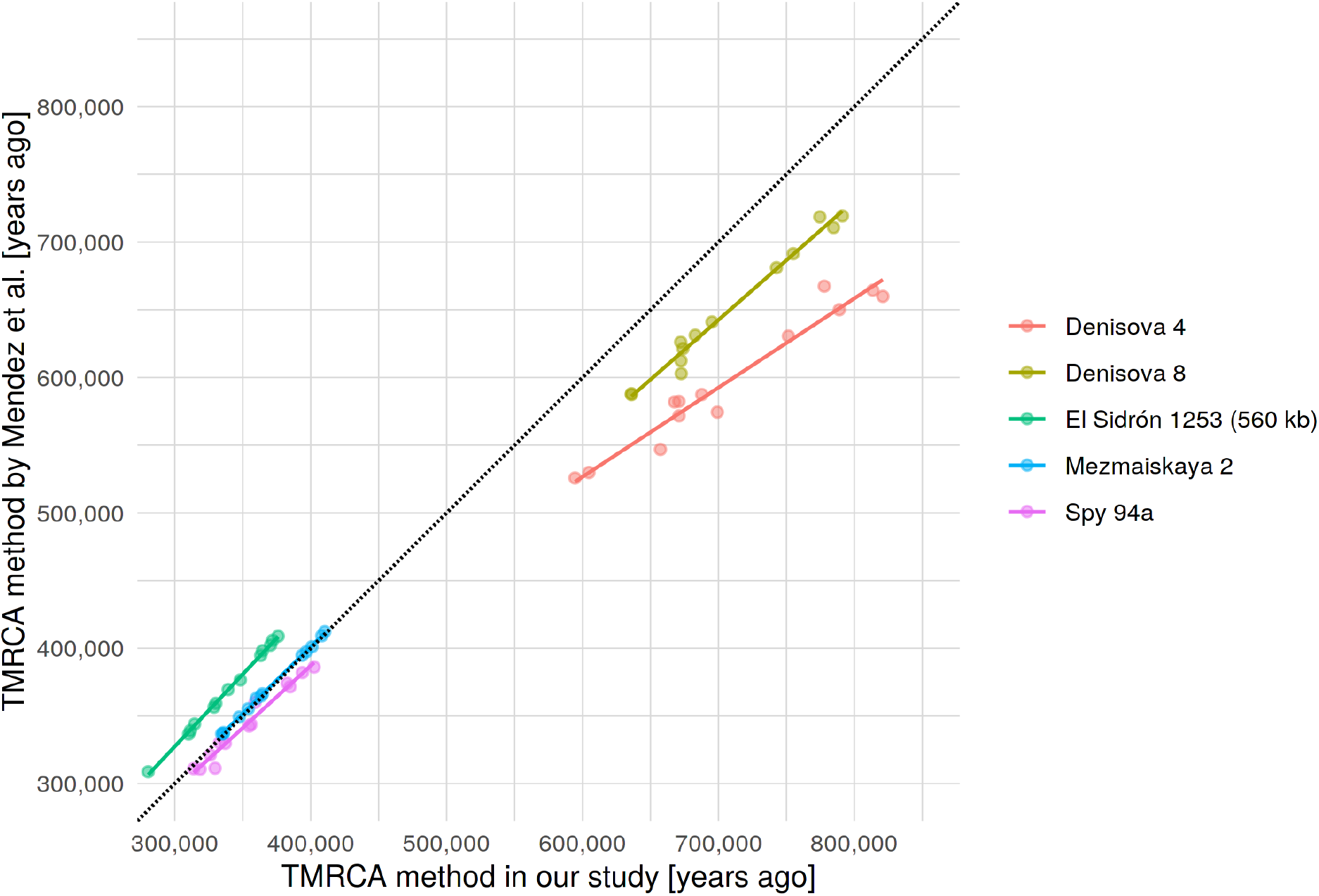
Comparison of TMRCA estimates obtained using a new statistic and the original approach used in the analysis of the *El Sidrón 1253* Neandertal. This figure shows the same data as in Figure S7.9 (panels on the left) but plotted on the same scatterplot for easier comparison. Dotted black line shows the the line of perfect correlation. The TMRCA between the Denisovan individuals and modern human is slightly underestimated due to a bias in the TMRCA estimation procedure proposed in the original study of the *El Sidrón 1253* Neanderthal (*5*). Analysis is based on all polymorphisms except C-to-T and G-to-A changes.

**Figure S7.11.**
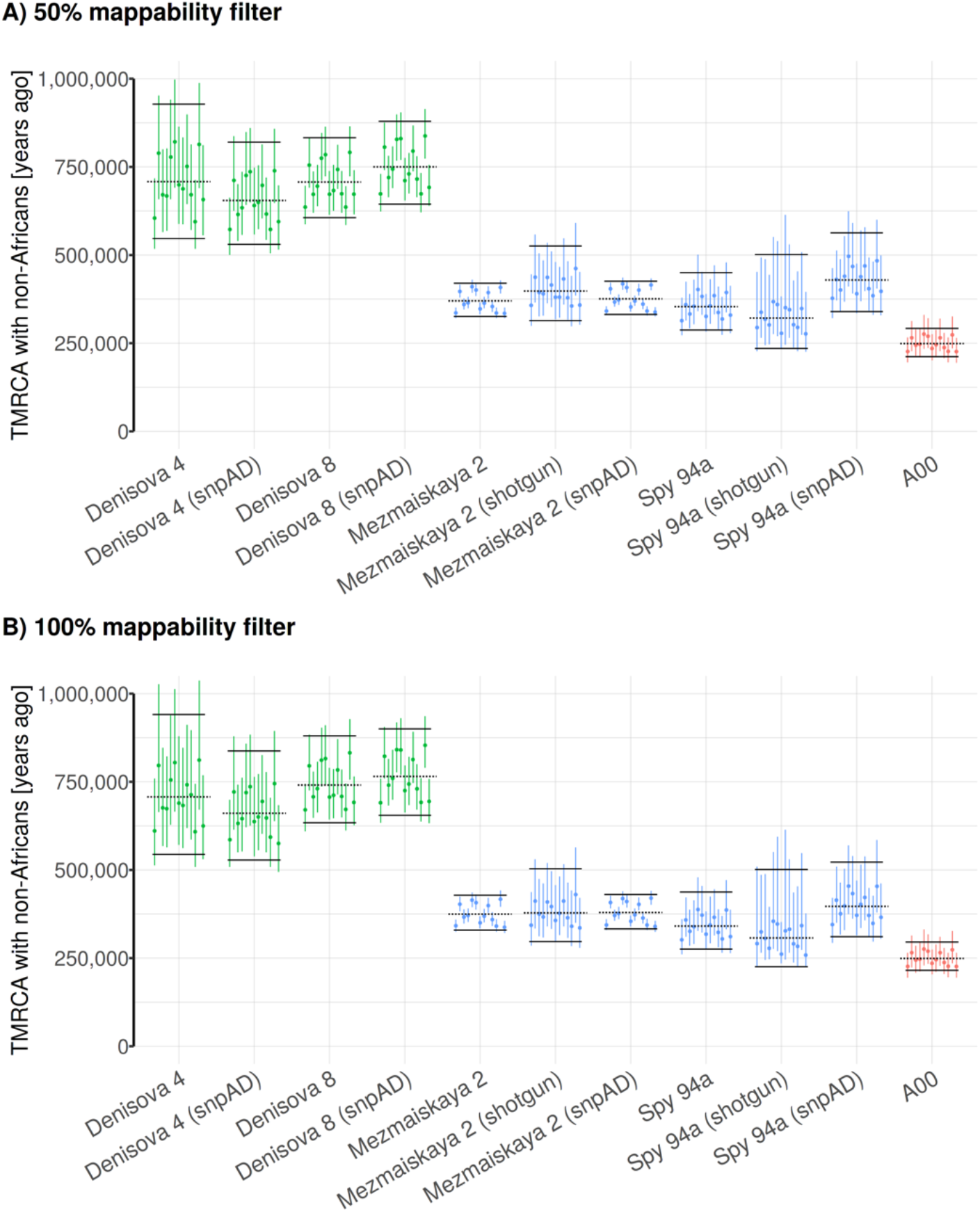
Detailed evaluation for potential technical biases in our TMRCA estimates. Shown are TMRCA results based on different versions of the data (shotgun or capture) or genotype calling methods (snpAD or consensus-based genotype calling method - which is the default). Panels (A) and (B) show results based on two versions of mappability filters - less strict (*“map35_50%”*) and more strict (*“map35_100%”*) filters described in the Altai Neandertal study (*2*). Analysis is based on all polymorphisms except C-to-T and G-to-A changes.

**Figure S7.12.**
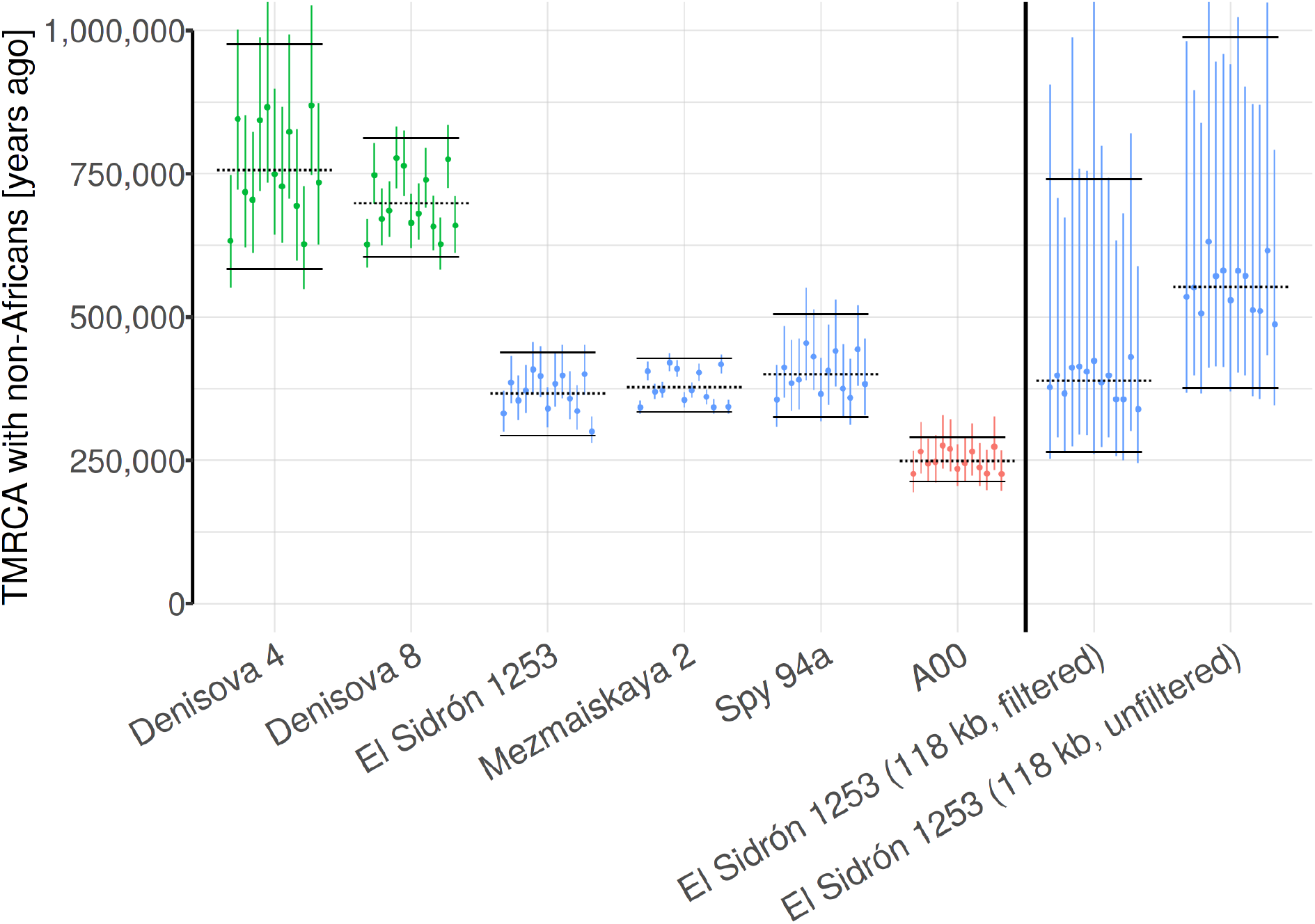
TMRCA between the *El Sidrón 1253* and modern human Y chromosomes. TMRCA estimates obtained for new capture sequence from the *El Sidrón 1253* Neanderthal (∼370 kya) differ significantly from the previously published results based on the 118 kb of the coding capture sequence (∼600 kya, “118 kb, unfiltered” column on the right from the vertical line). We found that applying stricter filtering criteria results in the same TMRCA values we obtain for the new capture data (“118 kb, filtered” column on the right from the vertical line). Analysis is based on all polymorphisms.

**Figure S7.13.**
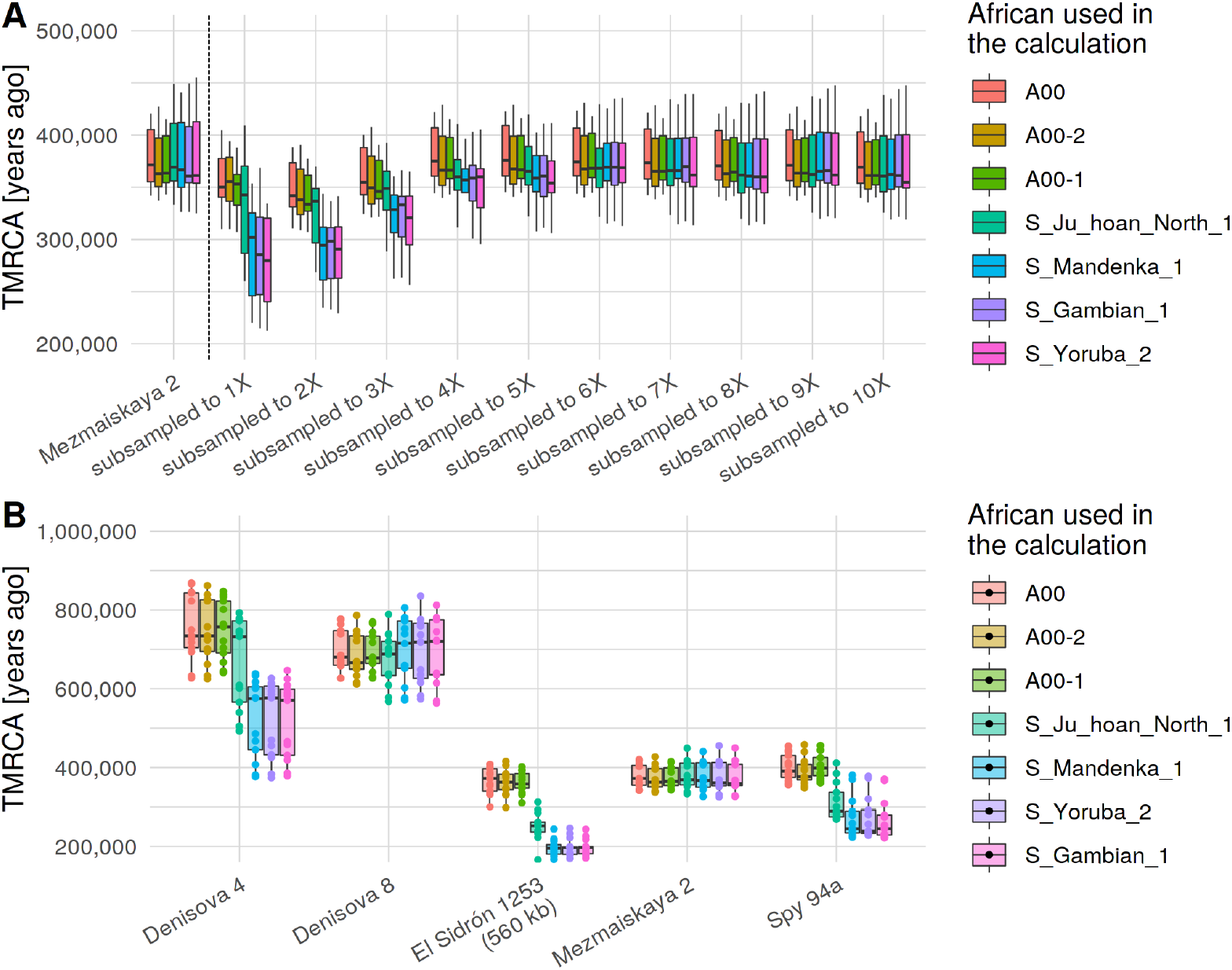
Estimates of *TMRCA_archanic_* for different African Y chromosomes used in the calculation. **(A)** Results for the high-coverage 14.3X *Mezmaiskaya 2* capture data (left of the vertical line) and its subsets generated by downsampling. A00-based TMRCA estimates are quite stable across the entire range of coverage and match those for the full data. In contrast, estimates based on other, less divergent, African Y chromosomes are heavily biased, and this bias is especially strong for low coverage samples. **(B)** A00-based estimates for *Denisova 4*, *El Sidrón 1253* and *Spy 94a* match those for their higher coverage counterparts (*Denisova 8* and *Mezmaiskaya 2*, respectively) as is required by the topology of the phylogenetic tree (Figure 2A). However, estimates based on other African individuals show the same bias shown for low coverage samples in panel (A). Dots show TMRCA estimates based on 13 non-African individuals. Both analyses are based on all polymorphisms.

**Table S7.3.** Observed branch counts and estimates of TMRCA between archaic and modern human Y chromosomes. All quantities represent averages across a panel of 13 non-African Y chromosomes (Table S7.1) and are based on A00-based estimates of mutation rate and *TMRCA_AFR_* (section 7.1). Counts in columns *a* to *f* represent counts of site patterns as shown in Figures S7.4 and S7.5. *“Total”* represents the number of sites out of the total 6.9 Mb of target sequence available for the analysis.

| name | a | b | c | d | e | f | total | TMRCA<br>[years<br>ago] | TMRCA<br>(lower CI) | TMRCA<br>(upper<br>CI) |
| --- | --- | --- | --- | --- | --- | --- | --- | --- | --- | --- |
| Denisova 4 | 142.2 | 2.0 | 2.2 | 77.8 | 125.8 | 165.9 | 1,084,363.6 | 708,133.1 | 549,422.5 | 930,979.7 |
| Denisova 8 | 583.5 | 8.8 | 8.5 | 318.0 | 410.4 | 583.9 | 3,372,262.6 | 706,874.9 | 607,187.2 | 833,211.4 |
| Mezmaiskaya 2 | 301.8 | 13.8 | 16.4 | 625.1 | 609.1 | 651.9 | 5,349,303.5 | 369,637.7 | 326,137.1 | 419,311.0 |
| Spy 94a | 17.4 | 4.5 | 0.2 | 42.0 | 49.5 | 87.6 | 510,735.5 | 353,265.5 | 286,250.7 | 449,185.5 |
| El Sidrón 1253 | 27.9 | 0.0 | 2.5 | 78.8 | 38.2 | 62.3 | 414,420.4 | 339,207.2 | 274,711.4 | 408,161.1 |

### 7.3. TMRCA of *Mezmaiskaya 2* and Spy *94a*

The split time of Neandertal and modern human Y chromosomes estimated in the previous section provides an upper bound for the last time the two populations experienced gene flow. Similarly, the deepest divergence in late Neandertal Y chromosomes represents a lower bound, as the introgressed Y chromosome lineage must have already been present in Neandertals prior to this diversification.

To estimate the deepest TMRCA of the known Neandertal Y chromosomes (*i.e.* the TMRCA of *Mezmaiskaya 2* and *Spy 94a*, Figure 2A), we first defined a set of sites in the ∼6.9 Mb capture target regions which carry a reference allele in the chimpanzee, A00 and French Y chromosomes (the ancestral state) and an alternative allele (the derived state) on the branch leading to the high-coverage Y chromosome of *Mezmaiskaya 2* (Figure S7.14), using the standard filtering used in previous sections (minimum three reads covering each genotyped site, section 5.3). We can calculate the approximate length of this branch using the TMRCA of *Mezmaiskaya 2* and modern human Y chromosomes (∼370 kya, Table S7.3) and the known age of *Mezmaiskaya 2* (∼44 kya, (*11*)) as 370 kya – 44 kya = 326 ky (Figure S7.14). We can then estimate the split time between *Mezmaiskaya 2* and *Spy 94a* Y chromosomes using the proportion of *Mezmaiskaya-*derived sites which show the ancestral allele in *Spy 94a* (Figure S7.14). Specifically, if we let Y be the number of ancestral alleles observed in *Spy 94a* and *X* + *Y* be the total number of sites with genotype calls in *Spy 94a* at positions derived in *Mezmaiskaya 2*, we can express the TMRCA of Y chromosomes of the two Neandertals simply as

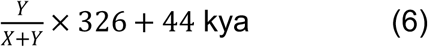

(Figure S7.14). We maximized the amount of data available for the analysis by merging the capture data with previously published shotgun sequences (*11*) and evaluated the robustness of the results to different genotype filtering and classes of polymorphisms. To estimate confidence intervals (C.I.), we re-sampled the *X* and *Y* counts from Poisson distributions with expected values given by the observed counts (Figure S7.14), calculated the TMRCA on the re-sampled counts using equation (6) and then took the 2.5% and 97.5% quantiles of this simulated TMRCA distribution to arrive at the approximate range of 95% C.I.

The TMRCA of *Mezmaiskaya 2* and *Spy 94a* are consistently around ∼100 kya regardless of the filtering criteria used (individual point estimates and 95% confidence intervals shown in Figure S7.15 and Table S7.4). Together with the TMRCA of Neandertal and modern human Y chromosomes, this suggests that the gene flow from an early population related to modern humans is likely to have happened some time between ∼100 kya and ∼370kya. We note that this time window is significantly wider than the one inferred based on a much more extensive set of available Neandertal mtDNA genomes (219-468 kya) (*38*). However, it is likely that future sampling of Neandertal Y chromosome diversity will reveal more basal Y chromosome lineages as has been the case for Neandertal mtDNA (*38*).

**Figure S7.14.**
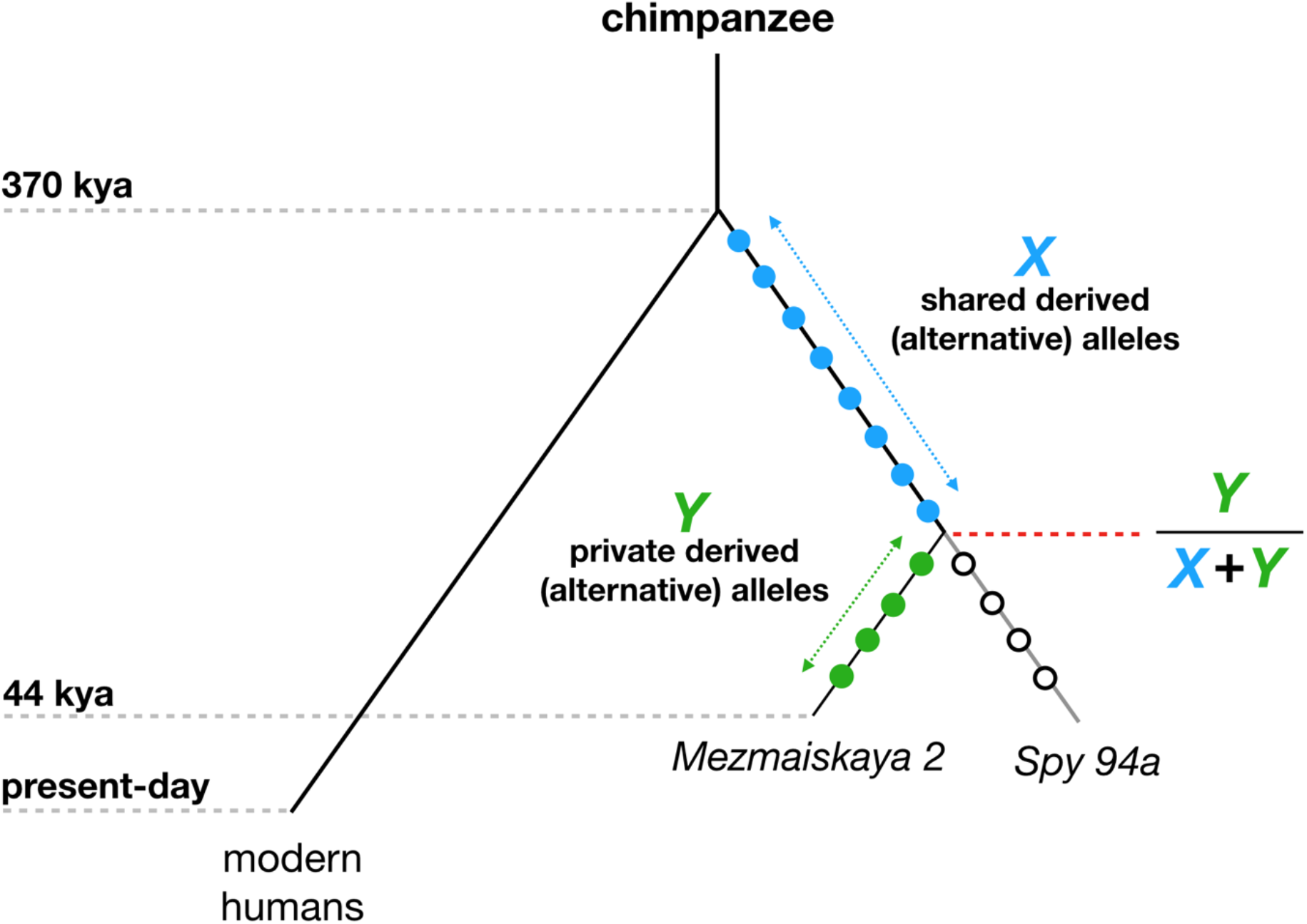
Estimating the TMRCA of Neandertal Y chromosomes. Filled circles represent a set of derived (alternative) alleles on the high-coverage *Mezmaiskaya 2* lineage, and are defined as sites at which *Mezmaiskaya 2* Y chromosome carries a different allele than chimpanzee, A00 and French Y chromosomes. Empty circles represent a subset of such sites which show the ancestral (reference) state in *Spy 94a*.

**Figure S7.15.**
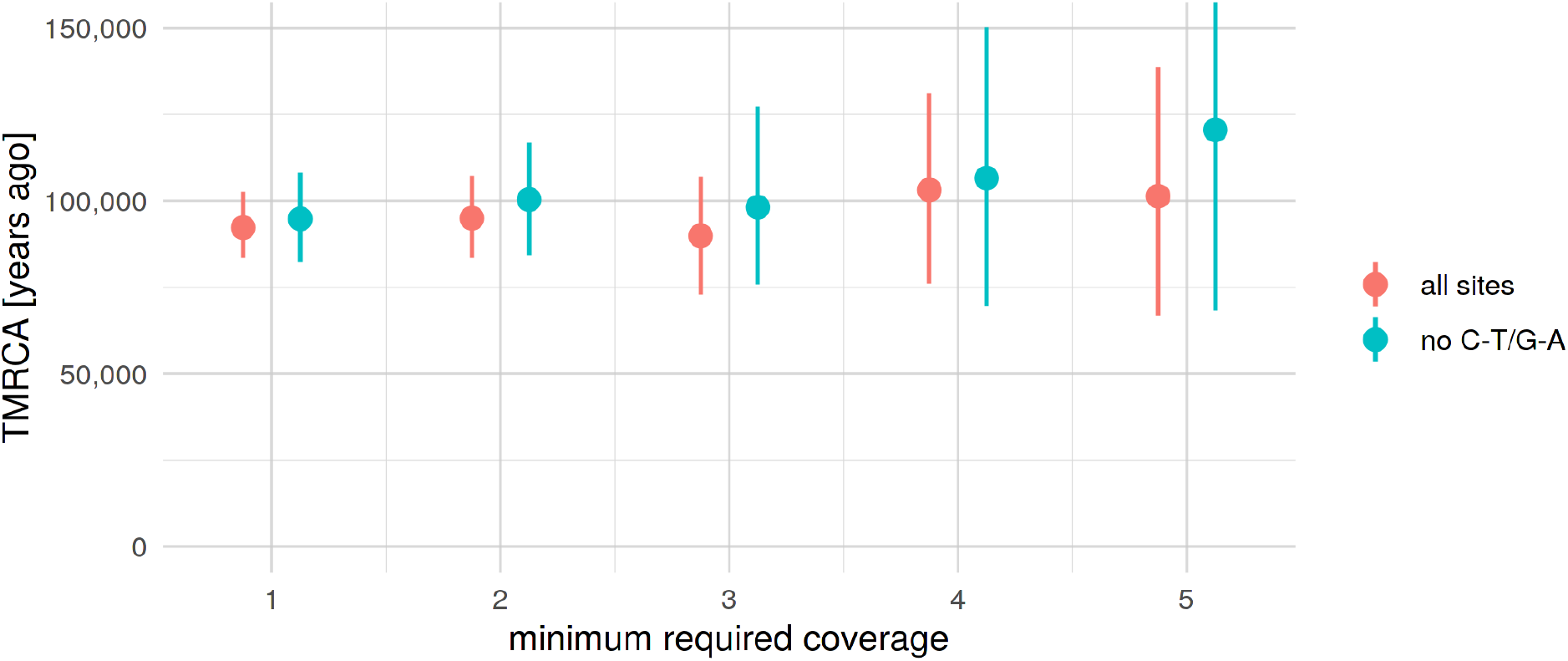
TMRCA of *Mezmaiskaya 2* and *Spy 94a*. Informative positions (derived alleles in *Mezmaiskaya 2*) were defined using genotype calls in *Mezmaiskaya 2* which passed the standard filtering used throughout our study (minimum coverage of at least three reads, maximum coverage less than 98% quantile of the total coverage distribution). We called the genotypes in the *Spy 94a* Y chromosome at these positions and calculated the TMRCA using the equation (6). We tested the robustness of the estimate to genotype calling errors using different minimum coverage filters for *Spy 94a* (x-axis) and two sets of polymorphisms (colors).

**Table S7.4.** Point estimates and 95% C.I. for the TMRCA of *Mezmaiskaya 2* and *Spy 94a* Y chromosomes as shown in Figure S7.15.

| REF<br>count | ALT<br>count | proportion<br>REF | TMRCA | TMRCA<br>(lower CI) | TMRCA<br>(upper CI) | minimum<br>coverage | calculated on |
| --- | --- | --- | --- | --- | --- | --- | --- |
| 71 | 469 | 0.1315 | 92250.32 | 83814.88 | 102157.6 | 1 | all sites |
| 51 | 314 | 0.1397 | 94962.33 | 83315.75 | 107309.6 | 2 | all sites |
| 20 | 141 | 0.1242 | 89862.87 | 74453.89 | 108808.4 | 3 | all sites |
| 13 | 66 | 0.1646 | 103130.37 | 77184.06 | 129130.5 | 4 | all sites |
| 7 | 37 | 0.1591 | 101332.33 | 67250.74 | 141779.7 | 5 | all sites |
| 43 | 266 | 0.1392 | 94775.67 | 82515.76 | 108105.1 | 1 | no C-T/G-A |
| 32 | 173 | 0.1561 | 100347.68 | 83703.34 | 117665.3 | 2 | no C-T/G-A |
| 13 | 74 | 0.1494 | 98152.86 | 75463.15 | 125756.6 | 3 | no C-T/G-A |
| 7 | 33 | 0.1750 | 106565.56 | 69559.13 | 149511.3 | 4 | no C-T/G-A |
| 5 | 18 | 0.2174 | 120510.02 | 67229.10 | 180578.4 | 5 | no C-T/G-A |

### 7.4. Confidence intervals

Under the assumption that mutations on each branch of a tree (Figures S7.1 and S7.4) accumulate independently, the observed counts of mutations can be understood as realizations of independent Poisson processes (mutation counts in Tables S7.2 and S7.3). To quantify the uncertainty in our TMRCA estimates, we used a simulation-based bootstrapping approach. For each set of branch lengths used to calculate scaling factor *α* (branches *a*, *d* and *e*, Table 7.3), we generated 1000 sets of simulated counts by randomly sampling from a Poisson distribution with the parameter λ set to values observed from the data. In other words, we simulated “trees” implicitly by generating a set of Poisson-distributed branch lengths. We then used the simulated counts to estimate the corresponding TMRCA values, obtaining a distribution of TMRCA consistent with the observed data. Finally, we took the lower 2.5% and upper 97.5% quantiles of the simulated distribution as the boundaries of bootstrap-based 95% confidence intervals.

To estimate the confidence interval for TMRCAs across the whole panel of 13 non-African Y chromosomes (black dotted horizontal lines in TMRCA figures in our study such as Figure 2B), we followed the same procedure but pooled all simulated counts together (i.e., 1000 simulated counts for each of the 13 Y chromosomes). Then, we took the lower 2.5% and upper 97.5% quantiles of TMRCA estimates calculated from the pooled counts.

### 8. Simulations of introgression under purifying selection

To investigate the expected frequency trajectories of Y chromosomes introgressed from modern humans into Neandertals, we adapted a modeling approach previously used to study negative selection against Neandertal DNA in modern humans (*39, 40*). Briefly, this model assumes lower effective population size (*N_e_*) in Neandertals than modern humans as inferred from comparisons of whole-genome sequences (*2*). Under nearly-neutral theory, such differences in *N_e_* are expected to increase the genetic load in Neandertals compared to modern humans through an excess of accumulated deleterious mutations due to lower efficacy of purifying selection. Therefore, after introgression from Neandertals into modern humans, Neandertal haplotypes would be under stronger negative selection compared to modern humans haplotypes, causing a rapid decrease in proportion of genome-wide Neandertal ancestry (*39, 40*).

In the context of evidence for Neandertal Y chromosome replacement in our study, we were particularly interested in the dynamics of introgression in the opposite direction, from modern humans into Neandertals. Specifically, given that nearly-neutral theory predicts that Neandertal Y chromosomes would carry a higher load of deleterious mutations compared to modern human Y chromosomes, how much is natural selection expected to favor introgressed modern human Y chromosomes compared to their original Neandertal counterparts?

To address this question, we used a forward population genetic simulation framework SLiM (version 3.3) (*41*) to build an approximate model of modern human and Neandertal demographic histories, following a strategy we used to study the long-term effects of Neandertal DNA in modern humans (*40*). To simulate differences in *N_e_* of both populations, we set *N_e_ = 10,000* in modern humans and *N_e_ = 1,000* in Neandertals after the split of both lineages from each other at 600,000 years ago. Given the non-recombining nature of the human Y chromosome, we implemented a simplified model of a genomic structure in which the only parameter of interest is the total amount of sequence under selection (“functional” sequence, ranging from 100 kb to 2 Mb in steps of 100 kb), and set the recombination rate to zero. Furthermore, because the amount of deleterious variation accumulated on both lineages is directly related the time they have been separated from each other, we simulated gene flow from modern humans into Neandertals between 150,000 to 450,000 years ago in steps of 25,000 years (this time range for gene flow encompasses the times inferred by (*38, 42, 43*) and our own study), assuming a fixed split time of 600,000 years ago. We ran 100 independent replicates for each combination of parameters described above, including an initial burn-in phase of 70,000 generations (7 × ancestral *N_e_* of 10,000) to let the simulations reach the state of mutation-selection-drift equilibrium.

In our previous study, we found evidence for different modes of selection in different classes of functionally important genomic regions, suggesting that the fitness consequences of mutations vary significantly according to the position of their occurrence (*40*). Realistic modeling of negative selection and introgression would thus require precise information about the distributions of fitness effects (DFE), dominance and epistasis for coding, non-coding and regulatory regions. Unfortunately, with the exception of DFE of amino acid changing *de novo* mutations affecting autosomal genes (*44, 45*), little is known about fitness consequences of non-coding and regulatory mutations on the Y chromosome. Furthermore, the impact of Y chromosome structural variation in the context of male fertility is highly significant, but still relatively poorly understood in terms of its DFE (*46, 47*). These issues, as well as the large parameter space of all relevant demographic and selection factors make analyzing the model dynamics quite challenging. To make our results easier to interpret, we scored each simulation run (i.e., each introgression frequency trajectory) with the ratio of fitness values of the average Neandertal Y chromosome and the average modern human Y chromosome generated by the simulation, just prior to the introgression. This way we collapse many potential parameters into a single relevant measure (how much worse are Neandertal Y chromosomes compared to modern human Y chromosomes in terms of evolutionary fitness) while, at the same time, generalizing our conclusions to other potential factors we do not model explicitly.

To calculate the fitness of simulated Y chromosomes, we used the fact that mutations in SLiM behave multiplicatively, i.e. each mutation affects any individual’s fitness independently of other mutations. Following basic population genetics theory (*48*), if we let the fitness of an individual be *W* and the fitness of each mutation be *w_i_*, we can Define *W* as *W* = Π*w_i_* = Π(1 − *s_i_*), where *i* runs across all mutations carried by this individual and *s_i_* is the selection coefficient of the *i*-th deleterious mutation. We can then transform multiplicative interaction into log-additive interaction by

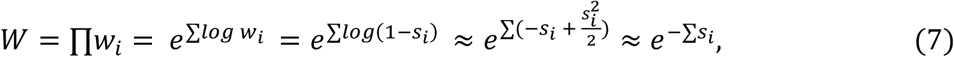

using simple rules of Taylor expansion under the assumption that we are dealing with weakly deleterious mutations whose selection coefficients (*s_i_*) are expected to be very small (*48*).

In practice, we let each simulation replicate run until modern human introgression into Neandertals, at which point we saved all Neandertal and modern human Y chromosomes and their mutations to a SLiM population output file. After introgression, which we simulated at 5%, we tracked the frequency of modern human Y chromosomes in Neandertals for 100,000 years, saving the frequency values at regular time intervals to another output file.

For the downstream analyses, we calculated the fitnesses of all Neandertal and modern human Y chromosomes from the population output files saved in the first step using equation (7) and calculated the ratio of the mean fitness values in both populations – this measure quantifies the expected decrease in fitness of Neandertal Y chromosomes compared to modern human Y chromosomes. The distribution of simulated fitness ratios across a two-dimensional parameter grid is shown in Figure S8.1. As expected, longer times of separation between Neandertals and modern humans and larger amounts of mutational target sequence increases the average genetic load of Neandertals. To analyze the dynamics of introgression in Neandertals over time, we scored the simulated modern human Y chromosome frequency trajectories with the calculated fitness decrease obtained from the simulation (Figure S8.2). Finally, we estimated the expected probability of replacement of the original Neandertal Y chromosomes over time by counting the proportion of the simulated trajectories (over all trajectories in each fitness bin) in which the introgressed modern human Y chromosomes reached fixation in each time point (Figure S8.2). These trajectories of replacement probabilities are shown in Figure 3B. Similarly to Figure S8.1, Figure S8.3 shows these probabilities expanded from the single fitness reduction score back on the two-dimensional parameter grid.

We note that although the simulation setup presented here is specific to Y chromosomes, the conclusions based on the abstract measure of fitness reduction can be generalized even to the case of mtDNA introgression (Figure 3B).

**Figure S8.1.**
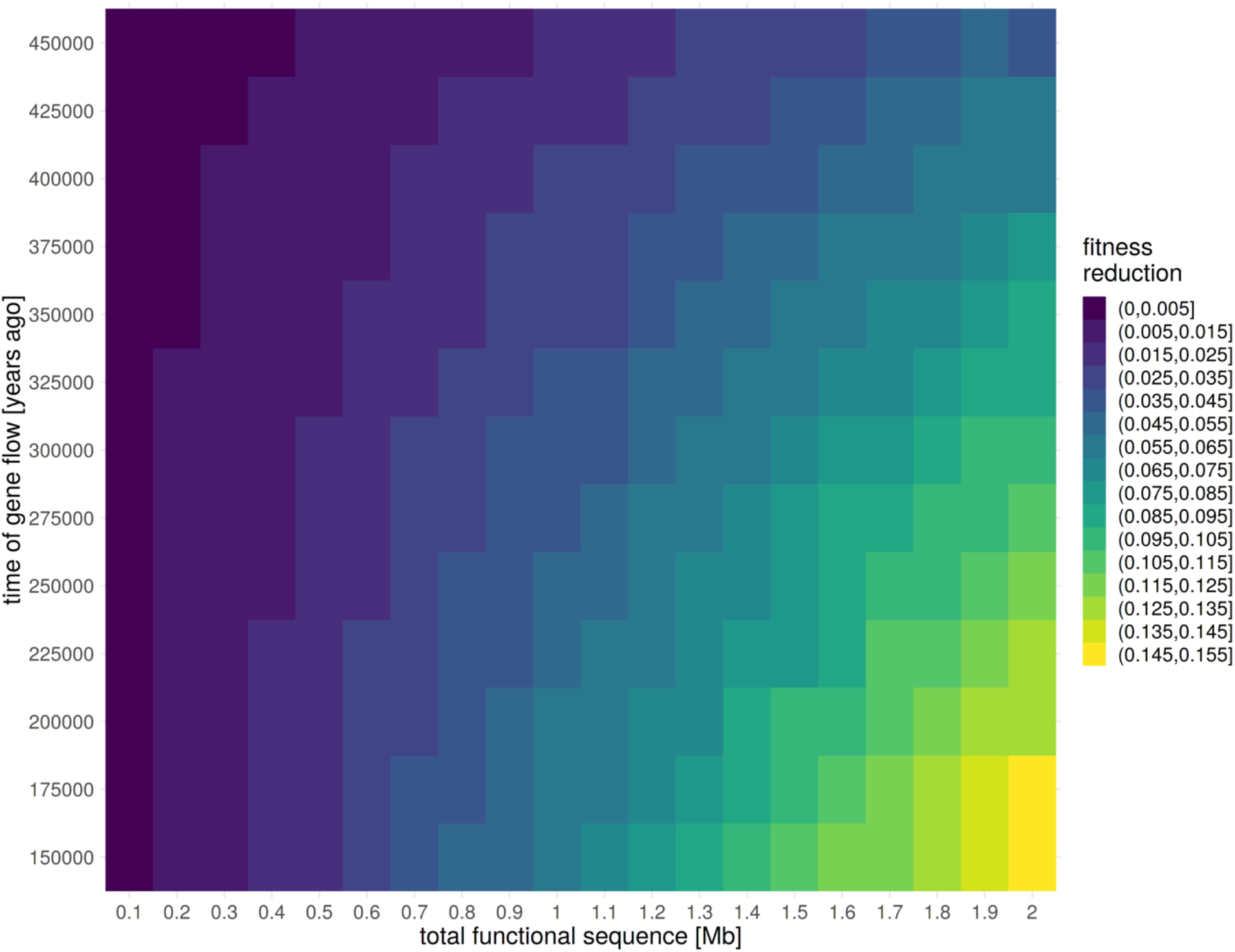
Expected decrease in fitness of an average Neandertal Y chromosome compared to an average modern human Y chromosome. Fitness decrease values were averaged over 100 independent simulation replicates on a grid of two parameters as the ratios of the mean fitness of Neandertal Y chromosomes to the mean fitness of modern human Y chromosomes (calculated using equation 7). Lighter colors represent lower fitness of Neandertal Y chromosomes compared to modern human Y chromosomes.

**Figure S8.2.**
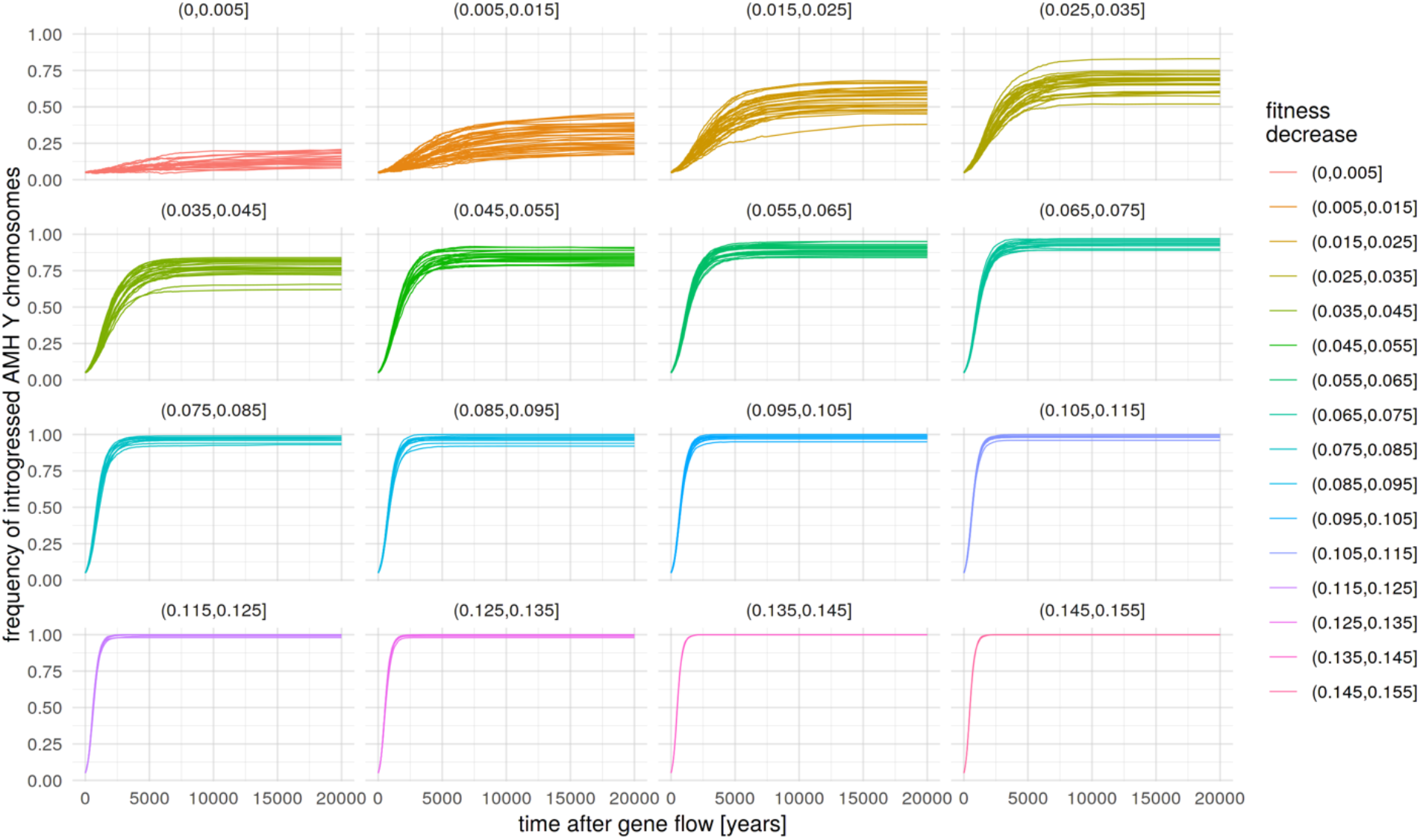
Frequency trajectories of introgressed modern human Y chromosomes in Neandertals, partitioned by the fitness decrease of Neandertal Y chromosomes compared to modern human Y chromosomes.

**Figure S8.3.**
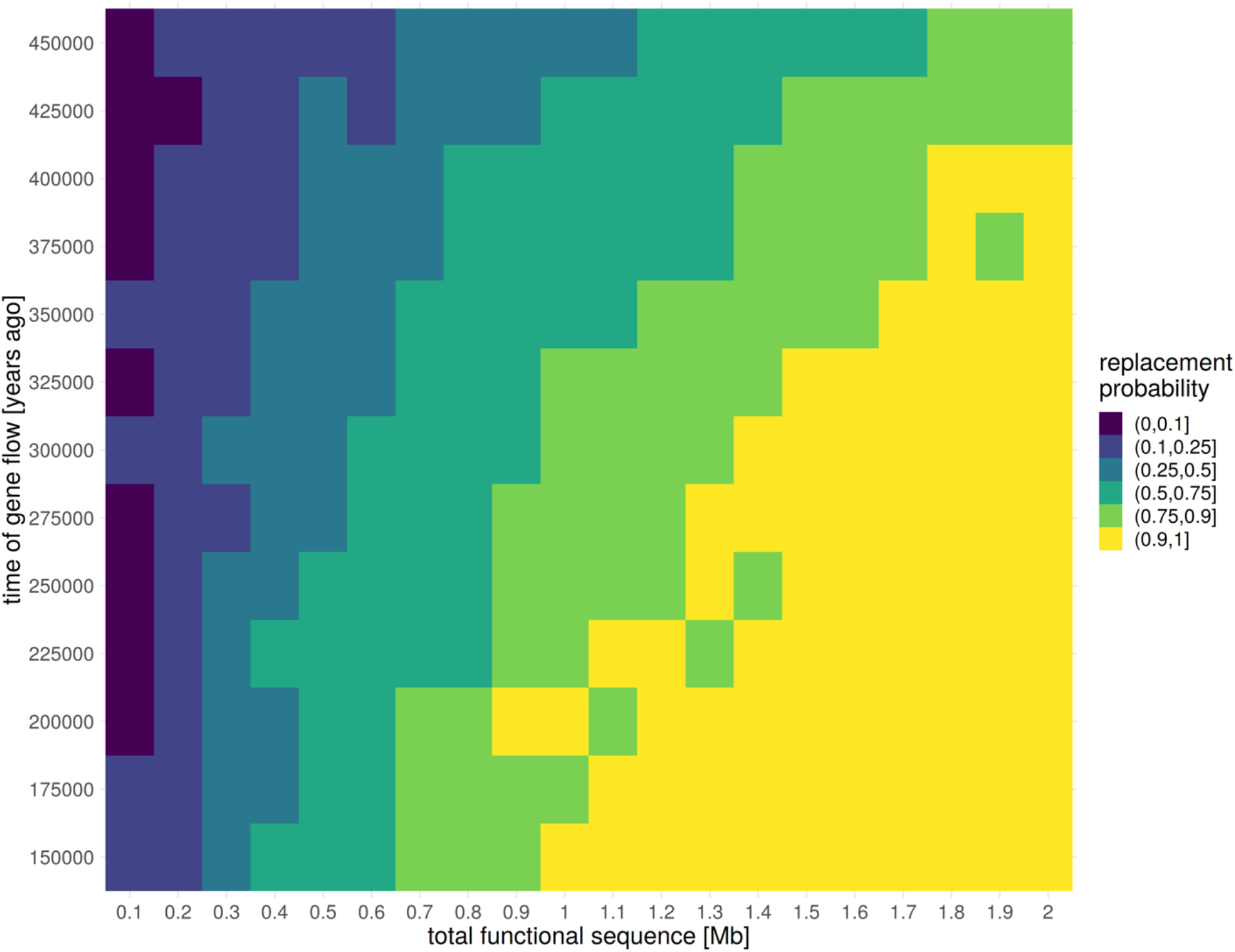
Probability of replacement of a Neandertal Y chromosome at 20 thousand years after gene flow from modern human. Probabilities represent the proportion of introgressed modern human Y chromosome trajectories that reached fixation in the Neandertals after 20 thousand years after gene flow, out of the total 100 simulation replicates performed for each combination of two-dimensional parameters (Figure S8.2).

---

**Table S2.1.**
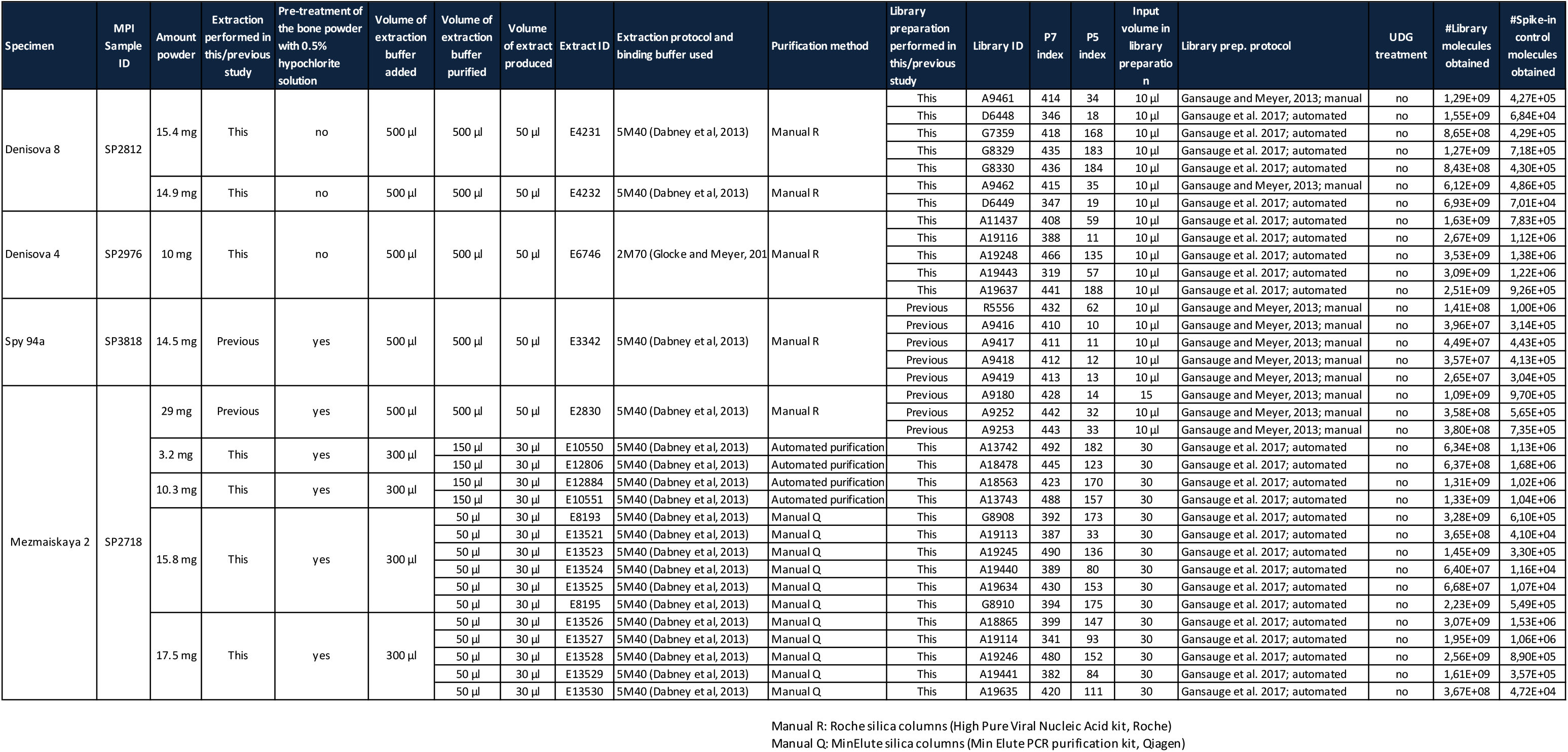
DNA library information.

